# Dynamics of bacterial operons during genome-wide stresses is influenced by premature terminations and internal promoters

**DOI:** 10.1101/2023.08.24.554578

**Authors:** Rahul Jagadeesan, Suchintak Dash, Cristina S.D. Palma, Ines S.C. Baptista, Vatsala Chauhan, Jarno Mäkelä, Andre S. Ribeiro

## Abstract

Bacterial gene networks have operons, each coordinating several genes under a primary promoter. Half of the operons in *Escherichia coli* have been reported to also contain internal promoters. We studied their role during genome-wide stresses targeting key transcription regulators, RNAP and gyrase. Our results suggest that operons’ responses are influenced by stress-related changes in premature elongation terminations and internal promoters’ activity. Globally, this causes the responses of genes in the same operon to differ with the distance between them in a wavelike pattern. Meanwhile, premature terminations are affected by positive supercoiling buildup, collisions between elongating and promoter-bound RNAPs, and local regulatory elements. We report similar findings in *E. coli* under other stresses and in evolutionarily distant bacteria *Bacillus subtilis, Corynebacterium glutamicum*, and *Helicobacter pylori*. Our results suggest that the strength, number, and positioning of operons’ internal promoters might have evolved to compensate for premature terminations, providing distal genes similar response strengths.

## Introduction

All organisms must coordinate the activities of their genes in time and strength. In bacteria and archaea species, those sets of genes are commonly placed adjacently in the DNA, in structures named operons (*1*), where they can be transcribed as single units. For this, upstream from the structural genes is (at least) one strictly regulated promoter (*2*), controlling their expression. This promoter is both where transcription initiates, as well as where most regulation of gene expression is exerted (*3*, *4*).

The dynamic correlation between genes in the same operon (*5*, *6*) can benefit the assembly of multi-subunit protein complexes (*7*) and the integration of laterally transferred genetic information (*8*). It may also assist co-regulating functionally related genes (*2*). In agreement, genes in the same operon are commonly of the same functional class (*9*, *10*).

Although genes in an operon share the same upstream promoter (named here ‘*primary promoter*’), some genes can nevertheless be differentially expressed from others (*11*), due to *internal* promoters, placed after the coding region of the first (or more) gene(s) of the operon (*12*). For example, the *cmk-rpsaA-ihfB* operon in *Escherichia coli* has a primary promoter, *cmkP*, and a few internal promoters (*12*). The most downstream of the internal promoters, *ihfBp2*, has preference for σ^38^, while the others have preference for σ^70^. Thus, while most genes of *cmk-rpsaA-ihfB* are preferentially expressed during exponential growth, *ihfB* is preferentially expressed during the stationary phase.

Another cause for differential expression between genes in the same operon is likely the occurrence of premature terminations of RNA polymerase (RNAP) transcription elongation events (*13*). There are several natural sources of premature terminations, such as pause-prone sequences, particularly when caused by RNA hairpin loops (*13*). Arrests can also influence, when ternary complexes remain intact, while further RNA synthesis is blocked (*14*). Homopolymer A/T tracts in the DNA (*15*), other attenuator sequences (*16*), RNAP collisions, and cAMP can also affect premature terminations (*17*, *18*). In the *gal* operon, in the presence of galactose, adding glycerol causes low transcriptional polarity, while adding glucose causes higher polarity, by enhancing premature terminations (*19*). Finally, environmental factors, such as temperature shifts, can affect premature dissociation rates in some operons, due to operon-specific mechanisms (*20*).

During genome-wide stresses, such as media shifts perturbing hundreds of genes at the same time (*21*), the responsiveness from internal promoters could have complex effects on the divergency between upstream and downstream genes. Further, these effects will likely depend on several factors, including premature termination rates, the locations of internal promoters in the operon, and the response strengths of primary as well as internal promoters. To study this, we measured the transcription dynamics of the operons of *E. coli* (Figure 1A) during genome-wide stresses targeting RNAP and gyrase (Figure 1B), since disturbing them should influence (quasi-synchronously) the activities of several hundred transcription start sites (TSSs) (*22*, *23*). From the data, we analyzed how the response strength of each gene, in each operon, relates to their position *P* in the operon (illustrated in Figure 1C). We also analyzed how the response dynamics of genes in the same operon differ with the distance *L_G_* between them (measured in ‘gene’ units as illustrated in Figure 1C). Finally, we evaluated the influence of the distances *L_P_* between consecutive promoters in the same operon (also measured in ‘gene’ units as illustrated in Figure 1C). In the end, we explore if our findings on the role of internal promoters are general or stress– and/or species-specific, using recently published data on *E. coli* and three other bacteria: *Bacillus subtilis*, *Corynebacterium glutamicum*, and *Helicobacter pylori*.

**Figure 1:**
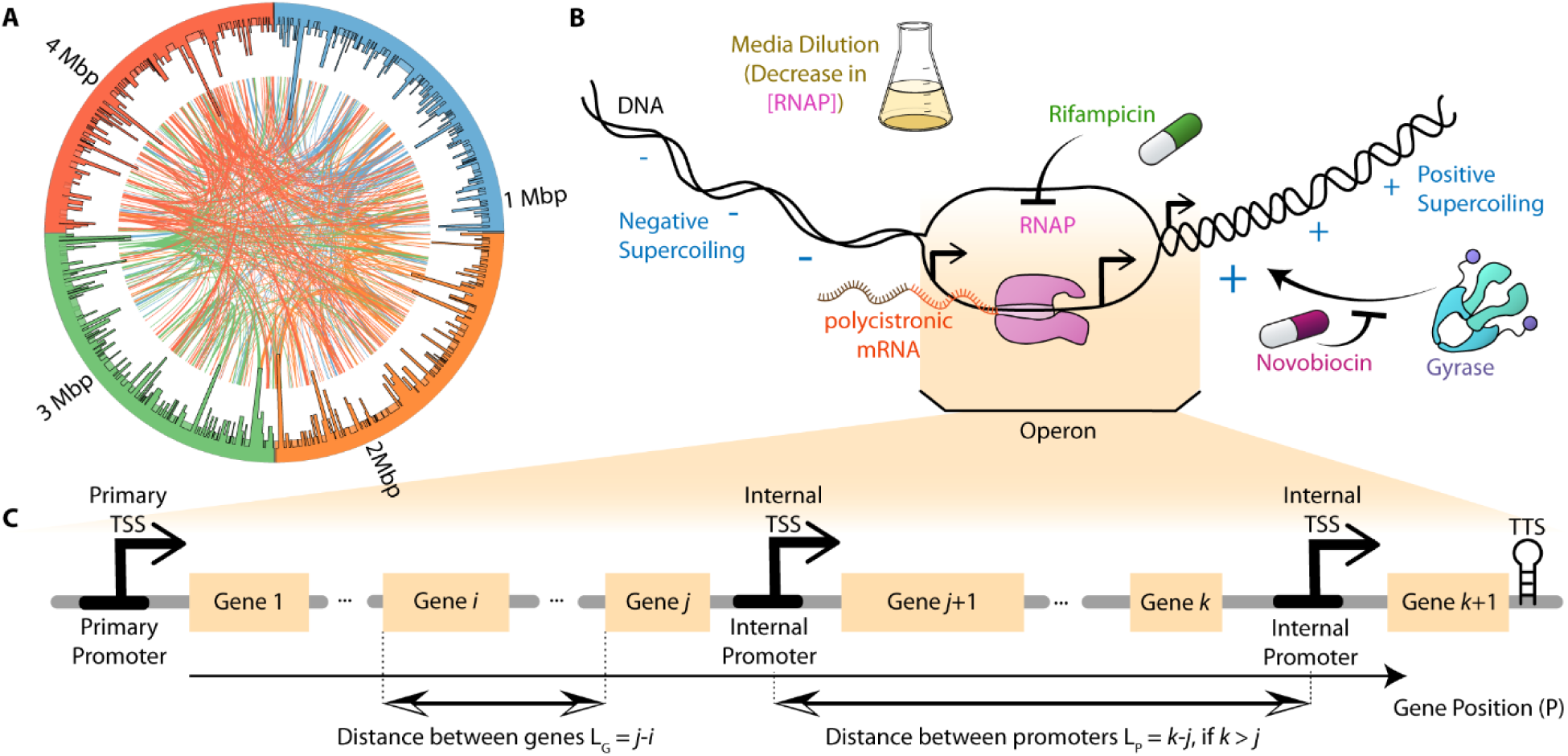
Illustrative summary of our study of the dynamics of operons during genome-wide stresses. **(A)** Illustration of the location of operons in the DNA was constructed using Circos. The external histogram shows the nucleotide length of each operon. Each curved line is a TF interaction between a gene in an operon and another gene. Colors of individual genes are DNA location-based, to facilitate distinguishing interactions. **(B)** Illustration of the three transcription-related, genome-wide stresses studied, specifically caused by media dilution (which reduces RNAP concentration), rifampicin (which blocks RNAP activity, and novobiocin (which inhibits gyrase activity), respectively. Positive and negative signs illustrate the accumulation of positive and negative supercoiling downstream and upstream the RNAP, respectively. **(C)** Illustration of the structural features of operons here considered. Shown are gene positions, *P*, (used to label genes) and the distances between genes in the same operon, *L_G_*. Also shown are the distances, *L_P_*, between consecutive promoters (specifically, between their TSSs). Here, if *k* is not larger than *j*, *L_P_* = 0. All distances and positionings are measured in ‘gene’ units. Finally, a transcription termination site, TTS, is illustrated at the most downstream location of the operon.

## Results

### Operons of *E. coli* have complex internal structure

*E. coli* has 833 operons, which account for 2708 of the 4724 genes (MG1655 strain (*12*)). The distributions of nucleotide lengths of operons and genes composing them are shown in Supplementary Figures S1A and S1B. Meanwhile, from Supplementary Figure S1C, in general, genes in operons are closely spaced, being separated by ∼50 nucleotides on average, regardless of their distance from the primary promoter (Figure 1C).

The number of known promoters in operons is 1174, while the number of transcription termination sites (TTSs) is only 174 (RegulonDB (*12*)). This suggests that, in operons, there are more opportunities to regulate initiations than terminations. In this regard, in RegulonDB, the location of each promoter in the DNA is defined by the positioning (in the DNA) of its nucleotide where transcription initiates, i.e. its TSS.

Meanwhile, 51% (422) of all operons have known transcription factor (TF) regulation. For comparison, only 24% (483) of the genes outside operons have TF regulation. This suggests that operons might be more subject to regulation under specific conditions.

We also studied the distance (in number of nucleotides) between operons and the next gene in the DNA, downstream from the end of the operon. (Supplementary Figure S1D). The average distance is 2× longer if there is no TTS in between (Supplementary Figure S1D). This suggests that there are substantial premature termination rates during elongation in non-coding regions of the DNA that ‘suffice’ to separate the dynamics of the operon from the closest downstream gene.

Many operons have more than one TSS (Supplementary Figure S1E). Namely, of the 1174 TSSs existing in operons, 532 (∼45%) are internally located. I.e., they are downstream from the most upstream gene of the operon.

Also, genome-wide, the numbers of internal TSSs are positively correlated to the nucleotide lengths of the operons (Supplementary Figure S1F). Since this correlation would also occur if internal TSSs were uniformly distributed in the DNA of operons, we constructed a null model of the same operons as *E. coli*, where the internal TSSs were spatially distributed uniformly.

Comparing the model with the empirical data, we find that the longer operons of in *E. coli* have more internal TSSs than expected by chance (Supplementary Figure S1F), while shorter ones have less. Thus, internal TSSs likely play a key role in the transcription dynamics of longer operons.

### The average response strengths of genes inside and outside operons are similar

To compare the stress responses of genes in operons with other genes, we exposed cells to three prominent stresses known to affect hundreds of genes by disrupting the transcription initiation process (directly or indirectly), and, thus, RNA production rates.

The first stress was caused by novobiocin. This antibiotic perturbs DNA supercoiling levels by partially inhibiting Gyrase (*23*, *24*), which is a protein responsible for regulating positive supercoiling buildup in the DNA (*25*, *26*). Consequently, novobiocin influences the TSSs’ availability to start transcription (*27*), RNAP elongation rates (*28*), and the dynamics of open complex formation (*27*). Since DNA supercoiling differs between DNA topological domains (*29*), the effects of novobiocin on operons differ with their DNA location. We studied both ‘strong’ (own data, Methods section ‘RNA-seq’) as well as ‘weak’ stresses (data from (*23*). We classified a stress as ‘strong’ when the resulting growth rates differ from the control condition (Supplementary Figure S2).

The second stress was caused by placing cells in poorer medium (created by diluting LB medium) (Methods section ‘RNA-seq’). This provokes RNAP concentration to reduce systematically and rapidly, without altering growth rates substantially (*4*, *22*). The data is from (*22*) at two time-points: i) following partial changes in RNAP levels and initial gene responses (∼120 minutes after the media shift); and ii) following the changes in RNAP levels and in TF levels, which further affect RNA levels (∼180 minutes after the media shift, during late exponential growth).

The final stress was caused by rifampicin (own data, Methods section ‘RNA-seq’), an antibiotic that specifically binds to RNAP, hampering promoter escape, by blocking transcription elongation after the initial 2-3 nucleotides of the RNA are added (*30*). Rifampicin also affects DNA replication and triggers general stress responses (*31*).

Supplementary Figure S3 shows the genome-wide distributions of single-gene response strengths to each stress (calculated by the log2 fold changes, *LFC*, of RNA read counts between the control and stress condition). Also shown are the corresponding means (*μ_LFC_*) and absolute means (*μ_|LFC|_*). Only the absolute means increase with the stress strength. On average, genes in operons do not differ in absolute response strength from other genes (Supplementary Figure S3).

### Gene response strengths do not show a consistent trend as a function of their position in the operon

We next studied the genes’ stress response strength, *µ_|LFC|_,* as a function of their positions in the operons, *P* (Figure 1C). We searched for a simple trend, common to all stresses. However, in some stresses *µ_|LFC|_* decreases weakly with *P*, while in other stresses it does not (Supplementary Figure S4). Moreover, *µ_|LFC|_* fluctuates with *P*.

Thus, using the same data, we instead studied the mean absolute differences in response strengths between genes in a given position, *P*, and the other genes of the same operon, *µ_|ΔLFC_*_|_, as illustrated in Supplementary Figure S5. From the same figure, *µ_|ΔLFC_*_|_ changes with *P* non-monotonically. In general, it first increases with the distance. However, for longer distances (∼5 or more genes), it starts decreasing with the distance. This distance (∼5) is similar to the average distance between promoters in the same operon (∼4), suggesting that the two results might be related.

Moreover, we find that operons differ widely in how their genes’ responsiveness differs with their positioning. In some operons, the genes exhibit changes with *P*, while in other operons they do not. This suggests that the changes in genes response strengths with *P* depend on the operons’ internal structure and, potentially, on external regulators.

### Differences in response strengths differ non-monotonically with the distance between genes in the same operon

Based on the results above, we next considered how the distance (in gene units) between genes in the same operon, *L_G_*, relates to the differences in response strengths, *µ_|ΔLFC|_*, as illustrated on the top of Figure 2.

**Figure 2:**
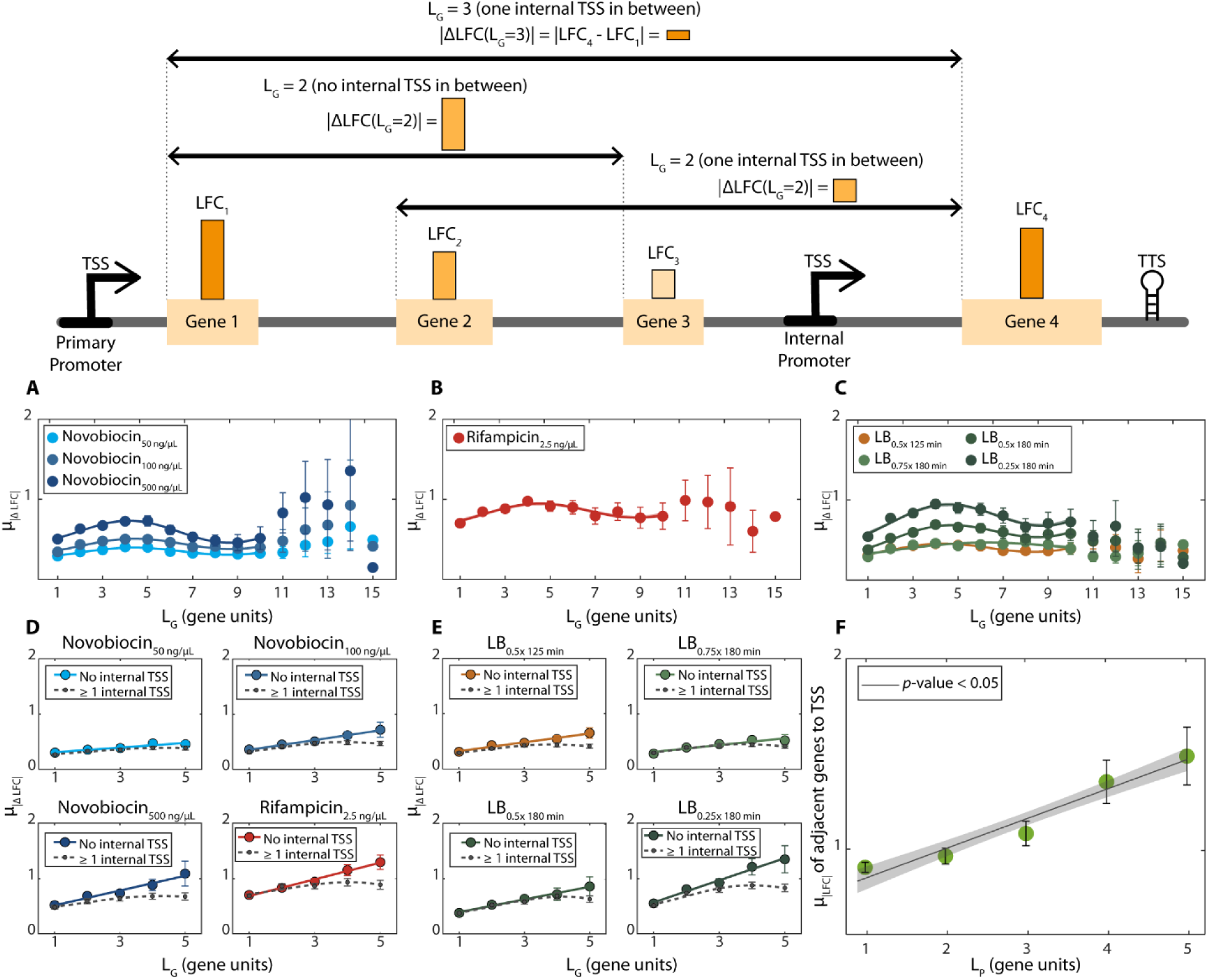
Average absolute differences in the response strengths of genes in the same operon (*µ_|ΔLFC|_)* as a function of the distance between genes, *L_G_*. Data from all *E. coli*’s 833 operons. Top: illustration of data used in Figures 2A-2F. Primary and internal promoters (TSSs shown) in operons respond to genome-wide stresses, while premature elongation terminations decrease responsiveness. Mean differences in between genes responses, *µ_|ΔLFC|_*, should first *increase* with *L_G_* due to premature terminations, but then *decrease* as internal promoters enhance downstream responses. Figures 2A-2F show corresponding empirical data. **(A)** Novobiocin-based stresses. **(B)** Rifampicin-based stresses. **(C)** Shifts in RNAP concentration due to shifting media richness. (**A-C**) *µ_|ΔLFC|_* from all pairs of genes in the same operon versus *L_G_*. Error bars are the standard error of the mean. Best fits are of the form of *Eq. 1*. Supplementary Table S1 shows fitting parameter and R^2^ values. **(D, E)** Average absolute difference between LFCs of genes in same operon *(µ_|ΔLFC|_)* with and without internal TSS’s in between (dotted and solid lines, respectively). Data only for *L_G_* < 6 as the number of pairs of genes without TSSs for *L_G_* > 5 is too small. See Supplementary Table S7, for the best fitting parameter values. **(F)** Average absolute response strengths, *µ_|LFC|_*, of genes adjacent (downstream) to TSSs, plotted against the distance (in ‘genes’) from their TSS to the nearest downstream TSS in the same operon, i.e. *L_P_* (illustrated in Figure 1C). In figures A, B, C, and F, the shadow areas are the 95% confidence intervals (not visible in most cases). In Figures D and E, the dashed lines correspond to the cases (≥ 1 TSS). In (D-F) we also show best linear fits, along with their *p*-values (Methods section ‘Statistical tests’).

From Figures 2A to 2C, in all stresses, the response strengths of genes in the same operon differ with the distance between them according to a wavelike pattern. Consequently, *µ_|ΔLFC|_(L_G_)* can be well fitted by wavelike functions, such as (but not exclusively) a sine function of the form:

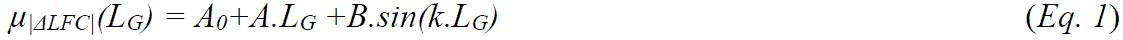

Here, *A_0_* stands for the y-axis intercept, while *A* and *B* are, respectively, the coefficient of the linear term and the amplitude of the best fitting function. Meanwhile, k = *2π.f* is the wavenumber (number of oscillations per unit length), where *f* is the spatial frequency of the sine function. Supplementary Table S1 shows the parameter values of each fitted curve.

As expected, for all stresses tested, the amplitude increases with the stress strength.

Meanwhile, the spatial frequency, *f*, differs little between stresses (ranging only from 0.09 to 0.11, with a mean of 0.10 and a small standard deviation of 0.01). Thus, the wavenumber might be largely independent from both the mechanism(s) affected by the stress, as well as the stress strength.

We observed a similar behavior, not previously reported, in single-cell protein numbers (*32*). We studied 444 different proteins in *E. coli* cells moved from an overnight culture into a fresh medium (*32*), and extracted 207 absolute differences between the protein numbers of genes in the same operon *(_|_ΔP_n_|).* Similar to mRNA levels, *µ_|ΔPn|_* follow a wavelike pattern with the distance between the genes coding for the proteins (*f* ∼ 0.17) (Supplementary Figure S6, Supplementary Table S2). For comparison, for the same 207 pairs of genes, a sine function fitted to *µ_|ΔLFC|_(L_G_)* has an average *f* ∼ 0.10 and a standard deviation of 0.01 (averaged from the stresses above).

Potentially, the wavelike patterns emerge from two or more influential factors, perhaps with opposite effects. Moreover, the impacts of these factors are likely to change differently with the distance between genes. Contrarily to the amplitudes of the fitted wave functions, the wavenumbers are very similar for all stresses (and stress strengths). This suggests that the wavenumbers might be mostly dependent on the structural properties of the operons, such as their length and internal promoters, among other.

### The sinusoidal curves of **µ_|ΔLFC|_(L_G_)** cannot be explained by transcription factor regulation

Next, we investigated if transcription factors (TFs) are necessary for observing the wavelike functions in Figures 2A-2C. For this, we plotted *separately µ_|ΔLFC|_(L_G_)* for the genes in the 427 operons that are *not* subject to TF regulation (*12*) (Supplementary Figure S7 and Supplementary Table S3) and for the genes in the 406 operons that *are* subject to TF(s) regulation (Supplementary Figure S8 and Supplementary Table S4), respectively.

Both operon cohorts exhibit wavelike patterns that can be well fitted by similar wave functions whose spatial frequencies for the same stresses differ by less than ∼10%. The only difference is ∼15% higher amplitude in the waves for operons subject to TF regulation. This can be explained by stronger response strengths, likely due to the responses being enhanced at subsequent times by changes in the numbers of the TFs regulating them (reported in (*22*).

We further plotted the differences in response strengths of the 56 operons that express one or more self-regulator transcription factors (“self-TFs”) (Supplementary Figure S9 and Supplementary Table S5). Again, sine functions describe well the behavior (frequencies differ by 20% from the operons above).

Overall, transcription factor interactions (at the genome-wide level) do not influence the wavelike patterns substantially. This holds true even when the responses of genes directly interacting by TFs exhibit correlation, e.g., following media dilutions (Supplementary Figure S10). On the other hand, this result does not imply that TFs do not influence the stress responses of genes in operons.

Namely, below we show that, locally, TFs influence differences in response strengths of adjacent genes in the same operon.

### The distances between genes in the same operon relate to their mean absolute differences in response strengths

Next, we studied if the differences in the response strengths of genes in the same operons relate to the distance between them. For that, first, we used *in silico* null models, where either the genes’ positioning in the operon, or the operon that the genes belong to, or both are randomized.

Using these computational models, we found that when randomizing the genes positions in their operons *in silico*, wave patterns are no longer visible (Supplementary Figure S11). The same occurs when randomizing the operon that each gene belongs to (Supplementary Figure S11). Finally, the same occurs when randomizing the position as well as the operon of each gene (Supplementary Figure S11). We thus hypothesized that the wavelike patterns of *µ_|ΔLFC|_(L_G_)* during the stress responses could be related to premature RNAP fall-offs from the DNA.

Finally, we observed that in natural operons, the differences in response strength between the most downstream gene in each operon and the first gene downstream the end of the operon is ∼2× higher (Supplementary Table S6) than between adjacent genes in the same operon (Figures 2A-2C). We conclude that the amplitudes of the waves are also influenced by the dynamic correlations between genes in the same operon.

### In the absence of internal TSSs, the mean absolute differences in response strengths of genes in the same operon, µ**_|ΔLFC|_**, are linearly correlated with the distance between them

We investigated how premature terminations of RNAP elongation influence the wavelike patterns in Figures 2A-2C. For that, we studied the differences in response strengths, *µ_|ΔLFC|_*, between genes *without* internal TSSs in between. For these, the *µ_|ΔLFC|_* increases linearly with the distance, *L_G_* (Figure 2D and 2E, solid lines & Supplementary table S7). Comparatively, having one or more internal TSSs in between causes *µ_|ΔLFC|_* to increase less with *L_G_* (Figures 2D and 2E, dotted lines).

To test the impact of where internal promoters are located, we used null models. First, we randomly switched the information on which pairs of genes have internal promoters in between. As we increased the fraction of pairs switched, *µ_|ΔLFC|_*(*L_G_*) differed less and less between the two cohorts of gene pairs. Having flipped 30% of the gene pairs, in all stresses, statistically significant differences in *µ_|ΔLFC|_*(*L_G_*) of the two cohorts disappeared (Supplementary Figure S12).

Next, we hypothesized that if premature terminations occur at significant rates and if internal promoters influence downstream genes, then, on average, promoters controlling more downstream genes should have stronger responses, so that all of them are responsive. To test this, we plotted the response strengths, µ_|*LFC*|_, of genes adjacent (downstream) to TSSs against the distance (in ‘genes’) from the nearest upstream TSS to the nearest downstream TSS (*L_P_*) in the operon (Figure 1C). As hypothesized, the response strength increases linearly with the distance from the next downstream promoter (Figure 2F). Meanwhile, randomly shuffling the position of the internal promoters, removes any correlation between µ_|LFC|_ and *L_P_* (Supplementary Figure S13).

Finally, to support these conclusions, we calculated autocorrelations (Methods Section ‘Statistical tests – Autocorrelations’) between the responses of genes in the same operon, as a function of their positions in the operon. In operons without internal promoters, if elongating RNAPs frequently terminate prematurely during stresses, the autocorrelations should gradually decrease with increasing distance between genes and the upstream promoter. Meanwhile, we expect this trend to be disrupted by internal promoters. Finally, in operons lacking internal promoters, we also do not expect to find any trend if the information on the genes’ position is randomized. Supplementary Figure S14 A, B, and C, respectively, validated all three predictions.

Overall, we infer that one main role of internal promoters might be to override the effects of natural premature terminations of the transcriptional machinery during genome-wide stresses, strengthening downstream genes more distanced from primary promoters (and, thus, homogenizing the response strengths of genes in the same operon). This should be particularly relevant in operons of many genes that need to be co-expressed, e.g., to execute complex adaptations to a genome-wide stress that involves several proteins.

### Premature terminations influence differences in response strength of genes in the same operon

Results in Figures 2D and 2E can be explained by differences (between control and stress conditions) in the rates of premature terminations of elongating RNAPs and/or the rates of transcription initiation at the promoter regions. If true, there should exist a substantial difference between the numbers of RNA reads from the starting and the ending regions of each gene (illustrated in Figure 3A). Moreover, these differences should differ between control and stress conditions, as well as between stress conditions (also illustrated in Figure 3A).

**Figure 3:**
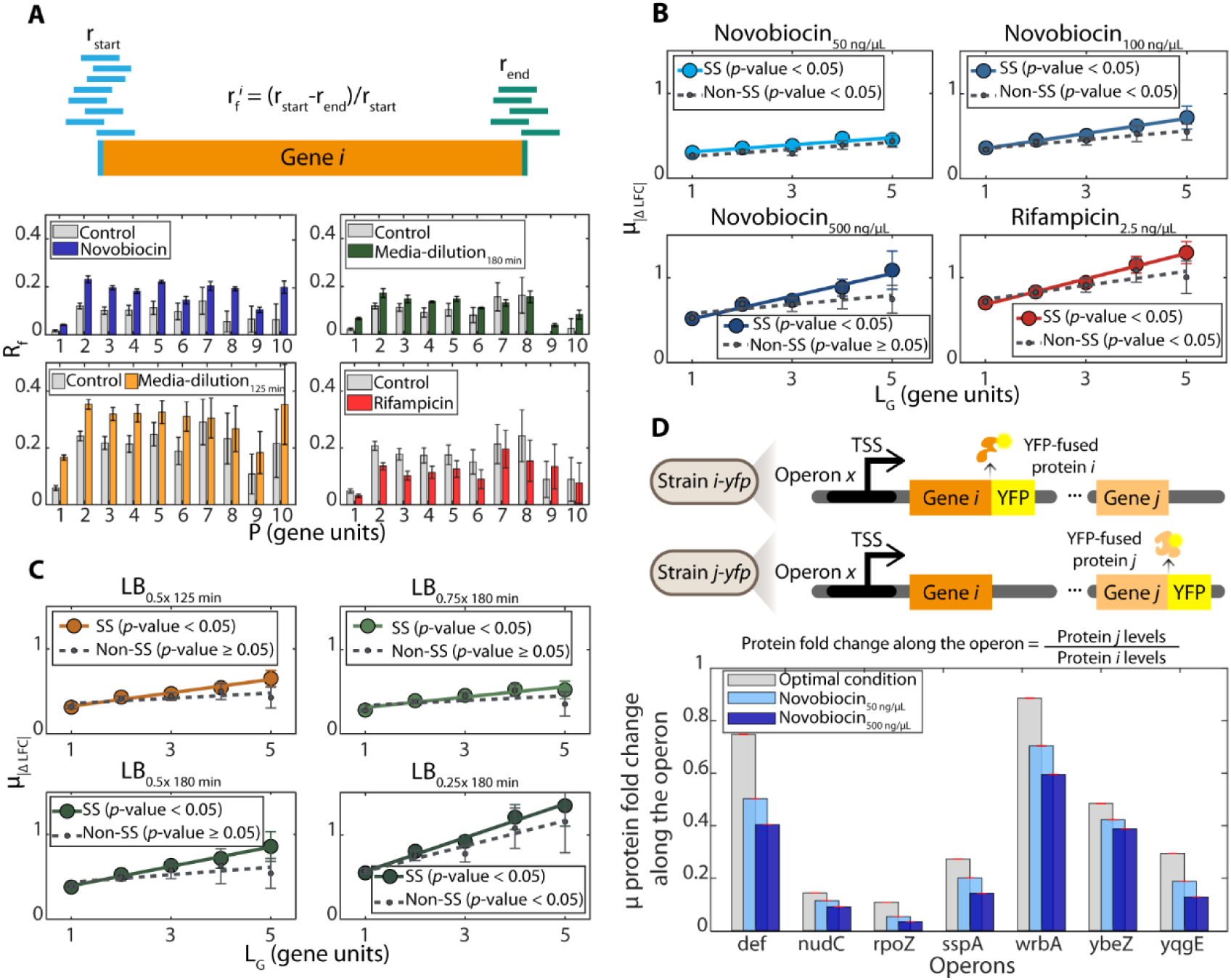
Effects of premature terminations on operon’s genome wide stress responses. **(A)** Illustration of raw data used to calculate, *r_f_*, which is the difference between numbers of RNA reads from the starting (blue lines) and ending (green lines) regions of a gene, normalized by the former. Below are the corresponding empirical data on genome-wide average values of *r_f_* (*R_f_*) as a function of the genes’ position *(P)*, relative to the primary TSS. Error bars are standard errors of the sum. **(B** and **C)** Average absolute difference between LFCs of genes in same operon *(µ_|ΔLFC|_)* with (colored) and without (grey) highly positive supercoiling buildup sensitive (‘SS’) genes in between them. Data for *L_G_* < 6 since there are not enough pairs of genes without TSSs in between for *L_G_* > 5. Solid lines correspond to pairs of genes in operons without internal TSSs. Dotted lines correspond to pairs of genes in operons without internal TSS and with positive supercoiling buildup sensitive (SS) genes. At 50 ng/µL, the responsiveness of genes with and without supercoiling sensitivity cannot be visually distinguished. The *p*-value of each line is from a statistical test of whether the linear fit differs from a horizontal line, using ‘fitlm’ (MATLAB). **(D)** Illustrated in the top are two strains differing in which gene of the same operon ‘x’ is fused with YFP, allowing protein detection by flow-cytometry, and then comparing levels as a function of positioning (and thus distances) in the same operon (Methods section ‘Flow cytometry’). Only pairs of genes without promoters in between were considered. The graph below shows the empirical data for several operons obtained using the YFP strain library (*32*). Strains are listed in Supplementary Table S18. Red error bars are the standard error of the fold change.

Using RNA-seq data, we estimated the average differences in the numbers of RNA reads from the starting and the ending regions of each gene, *R_f_* (normalized by the former). Values were obtained as a function of the positions of genes (Methods section ‘Estimating genome-wide differences in reads between the start and end of genes in operons’). Figure 3A shows that: i) the read counts aligned to the start and end regions of the gene coding regions differ substantially, suggesting non-negligible rates of premature terminations. Also, ii) *R_f_* does not differ with the position, suggesting that premature terminations do not occur at a few specific positions. Finally, iii) the premature termination rates differ between stresses and the control, which plays an important role in the formation of wavelike patterns in Figures 2A-2C. As a side note, we could not explain why *R_f_(P)* is much lower for *P* = 1.

As a side note, the existence of substantial premature terminations of transcription elongation could help explain why the presence of TTSs between operons and downstream genes in *E. coli* is more frequent if the two are closer to each other (Supplementary Figure S1D).

### Genes with high sensitivity to positive supercoiling buildup in standard conditions can be associated with higher premature termination rates

A past study showed that positive supercoiling buildup can force elongating RNAPs to pause (*33*). It can also destabilize DNA-bound proteins (*33*). In agreement, we observed that novobiocin, a gyrase inhibitor, increases the genome-wide rates of premature terminations, *R_f_*, regardless of gene positions (Figure 3A and Supplementary Figure S15). Thus, positive supercoiling buildup may partially explain some premature terminations. If true, genes in DNA regions more sensitive to positive supercoiling buildup should have higher differences in response strengths to the stresses.

To test this, we studied the differences in response strengths, *µ_|ΔLFC|_(L_G_)*, between genes, considering whether one of the two genes (or one of the genes in between them) has been classified as being highly sensitive to positive supercoiling buildup, based on their responsiveness to novobiocin (50 ng/µL), in cells originally in standard growth conditions (control) (*23*) (Methods 4.8). We only consider genes from operons without internal TSSs, since TSSs would interfere with the estimations of premature termination rates.

We find that the differences in response strengths is higher when genes have high sensitivity to positive supercoiling buildup (Figures 3B and 3C). Moreover, the difference increases with novobiocin concentration (Figure 3B and Supplementary Table S8). Finally, the differences are large enough to cause protein levels to also be distinct (Figure 3D and Supplementary Figure S16). In addition, we find similar results under the other stresses studied (Figures 3B and 3C and Supplementary Table S8). This suggests that the same DNA regions are more prone to premature terminations under these two stresses as well. Overall, positive supercoiling buildup likely partially influences the differences in response strengths of genes in the same operon, likely by influencing premature termination rates.

Meanwhile, from Figure 3A, when considering all operons (with and without TSSs), we observe that the normalized rates of premature terminations, R_f_, are smaller when cells are subject to rifampicin. This can be explained by the expected smaller number of elongating RNA polymerases, as they fail to escape transcription start sites.

On the other hand, in general, *R_f_* is stronger than the control under both media dilutions, similarly to when under Novobiocin. This may be partially related to positive supercoiling buildup. Namely, *E. coli* cells in poor media are known to have reduced internal energy levels, which can reduce their capacity in curating positive supercoiling buildup (*27*, *34*). In agreement, we found that the ATP levels of cells under medium dilutions were much lower than in the control (Methods section ‘Cellular ATP levels’ and Supplementary Figure S17). E.g. at 180 minutes after moving cells to increasingly diluted media, ATP levels were 15%, 25%, and 47% lower, respectively. Differences of such magnitudes are expected to hamper Gyrase ability to curate positive supercoils (*23*), likely disturbing premature termination rates, particularly in DNA regions with higher positive supercoiling buildup.

### Synthetic constructs suggest that internal promoters can cause premature terminations in all stress conditions

Contrarily to when subject to novobiocin and media dilutions, cells under rifampicin mostly exhibited premature termination rates lower than the control (Figure 3A). Meanwhile, rifampicin blocks RNAP escape from TSSs (*30*). As such, genome wide reductions of transcription rates are expected. Consequently, under this stress, positive supercoiling buildup might be lesser influential on the premature termination rates.

Instead, because RNAP escape from internal promoters is hampered, collisions between RNAPs elongating from upstream promoters with RNAPs bound to downstream promoters should become more frequent (thus, influential). I.e., we expect that some of those collisions will cause premature terminations (most likely of the elongating RNAPs but, in some cases, also of the bound RNAP, or even of both colliding RNAPs). Thus, internal promoters might not only enhance the responsiveness of downstream genes, but also enhance premature terminations. Their contributions to both phenomena likely differ between internal promoters and between stress conditions.

To test this, we use recently engineered synthetic genetic constructs of the P_tetA_ and the P_LacO3O1_ promoters in non-overlapping tandem formation (upstream and downstream, respectively) controlling mCherry expression (*35*). This construct, named ‘P_tetA-PLacO3O1_’, is used as a proxy for an operon with one internal TSS. The expected events following collisions between RNAPs leading to premature terminations are illustrated in Figure 4A. We use constructs with the same promoters in single formations for comparison (Supplementary Table S9).

**Figure 4:**
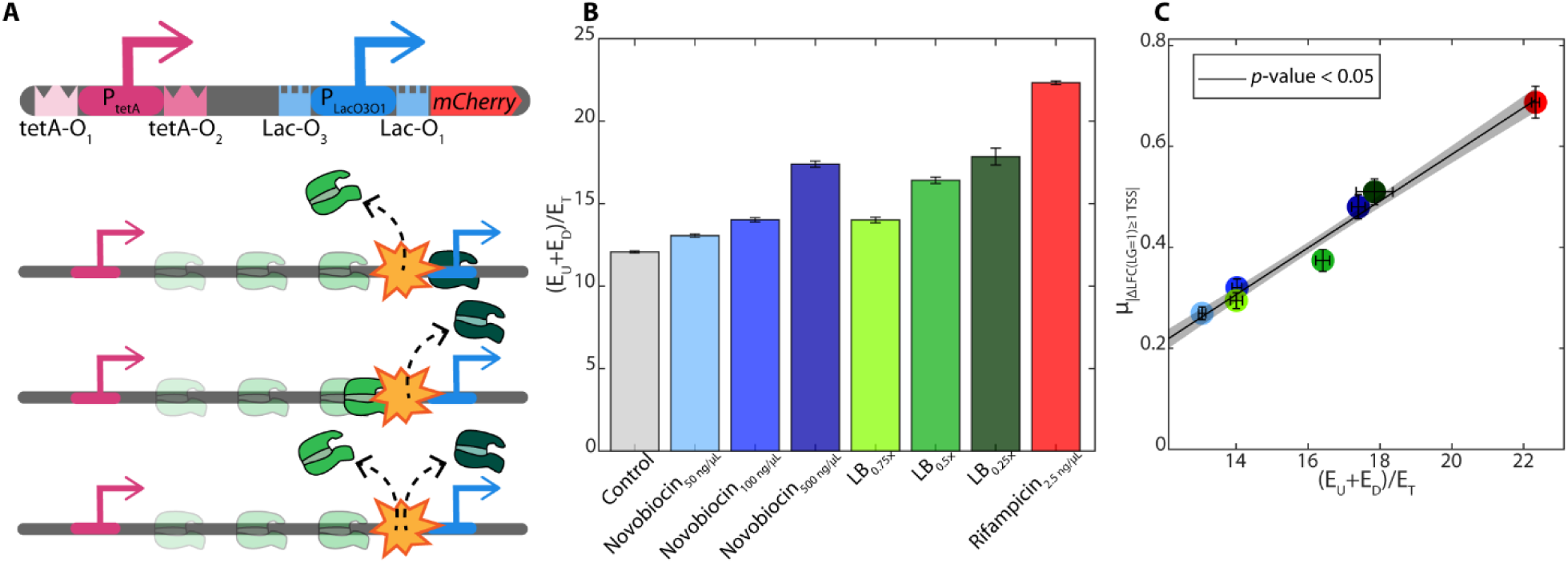
Synthetic genetic construct carrying two, non-overlapping promoters in tandem formation. (A) Illustration of the construct, including their transcription start sites (arrows) and transcription factor binding sites (‘TetA-O’ binding sites for anhydrotetracycline, aTc, in the case of the upstream promoter, P_TetA_, and ‘Lac-O’ binding sites for LacI, in the downstream promoter, P_LacO3O1_). Also illustrated are the expected possible premature termination events, following collisions between RNAPs elongating from the upstream promoter with RNAPs bound to the downstream promoter. (B) Summed expression levels of the two promoters in single formation relative to the expression of the non-overlapping promoters in tandem formation, for each stress condition and the control. (C) Mean absolute differences in response strength between pairs of genes with one internal TSS in between (µ_|ΔLFC|_(L_G_ ***=*** 1)_≥ 1 TSS_) in each stress condition plotted against the values in (B).

If RNAP collisions between elongating RNAPs and RNAPs (or transiently bound repressors) at internal promoters can cause premature terminations, the summed expression levels of P_tetA_ and P_LacO3O1_ (E_U_+E_D_) in single formations should be higher than of the non-overlapping tandem construct P_tetA_-P_LacO3O1_ (E_T_), even in control conditions. Moreover, increasing the time that RNAPs spend bound to the downstream promoter, e.g., by adding rifampicin, should increase the ratio between summed single expressions (E_U_+E_D_) and the expression of the tandem construct (E_T_). Results in Figure 4B support these hypotheses, and further suggest that the influence from premature terminations due to these collisions differs between stress conditions.

To assess if these results are quantitatively in line with the genome-wide data on the operons’ stress responses, we compared the constructs’ behavior under each stress, with the corresponding mean absolute differences in response strength between pairs of genes with one internal TSS in between (µ_|ΔLFC_|(*L_G_* = 1, with 1 TSS in between), as they are relatively similar in structure to the tandem construct. From Figure 4C, we find a strong linear correlation between the behaviors of the synthetic construct and the natural operons, suggesting similar underlying mechanisms and functioning.

We conclude that, while internal promoters in operons enhance the stress responses of their downstream genes, they nevertheless also increase premature terminations. When these terminations are of RNAPs elongating from upstream promoters, they could increase differences in response strengths of upstream and downstream genes. In general, Figures 2A-2C show that, during genome-wide stresses, their contribution to decreasing differences in responses outweighs their contribution to increasing those differences.

### Local DNA properties and other regulatory factors influence differences in the response strengths of adjacent genes

In this section, we evaluated the influence of other variables on local differences in the response strengths of adjacent genes in operons.

First, we investigated sites reported to be prone to premature transcription terminations (*pTTS’s*) reported in (*36*). We found that their presence is consistent with higher differences in response strengths between adjacent genes, due to causing higher local premature termination rates (Supplementary Section ‘Sites reported to be prone to premature transcription terminations’, Supplementary Figures S18 & S19).

Similarly, we found that the local differences between adjacent genes are influenced by the promoters’ σ factor preferences (Supplementary Figure S20), regulatory non-coding RNAs binding sites (Supplementary Figure S21), genes sensitive to (p)ppGpp (Supplementary Figure S22), and internal TFs binding sites (Supplementary Figure S23). For details, see Supplementary Section ‘σ factor preferences, sRNA regulation, (p)ppGpp, and transcription factor binding sites.

Contrarily, for specific RNAP pausing sequences, RNA degradation rates, distances from the origin of DNA replication, TSSs in the opposite DNA strand, or promoters AT-richness, we failed to find evidence that they influence the differences in response strengths between adjacent genes. For details, see Supplementary Section ‘RNAP pausing sequences, RNA degradation rates, distances from the origin of DNA replication, TSSs in the opposite DNA strand, and promoters AT-richness’, Supplementary Figures S24-S28 & Supplementary Table S10.

### Other stresses cause similar genome-wide responses in *E. coli*

If the genome-wide stress response dynamics of operons observed in Figures 2A-2C is largely due to premature terminations and internal TSSs, the same responsiveness should be detectable in past data on other stress responses by *E. coli* cells. To test it, we collected publicly available, processed RNA-seq data in NCBI for *E. coli* MG1655 subject to stresses (mutations not included), namely: changes in the microaerobic environment (*37*), heat-shock (*38*), hydrostatic pressure (*39*), chloramphenicol (*40*), erythromycin (*40*), and Mercury (*41*). Further, we studied anaerobic-to-aerobic shifts (at 0.5, 1, 2, 5, and 10 min, after the shift) (*42*).

Analyzing the data accounting for the operon structure of *E. coli*, in all cases, the differences in response strengths between genes in the same operon (*µ_|ΔLFC|_*(*L_G_*)), are well fit by a sine function of the form of *Eq. 1* (Supplementary Figure S29 and Supplementary Table S11). Moreover, the mean and standard deviation of the spatial frequency *f* of the best fitting lines are also 0.10 and 0.01, respectively, as in (Figures 2A-2C). Finally, as expected, the amplitudes also differ with the nature of perturbations (as in Figures 2A-2C).

### Other bacteria also exhibit similar genome-wide stress responses

Other bacteria have their genome organized in operons and their sizes (gene numbers) have been reported (*1*) (albeit the numbers and positioning of internal TSSs have not). Noteworthy, their operon size distributions (Supplementary Figure S30) cannot be distinguished statistically from the size distribution of *E. coli* (Supplementary Table S12). We thus hypothesized that, during genome-wide stress responses, the differences in the responses of the genes in operons, *µ_|ΔLFC|_*, should differ with the distance between them, *L_G_*, as in *E. coli*.

To test this, we collected data on various stress responses of *B. subtilis* (*43–49*), *C. glutamicum* (*50*, *51*), and *H. pylori* (*52*). From Figure 5, in all cases and species, *µ_|ΔLFC|_(L_G_)* is well fit by a sine function of the form (*Equation 1*). The best fitting spatial frequencies *f* (mean and standard deviation) for each stress and species are shown in Supplementary Table S13-15.

**Figure 5:**
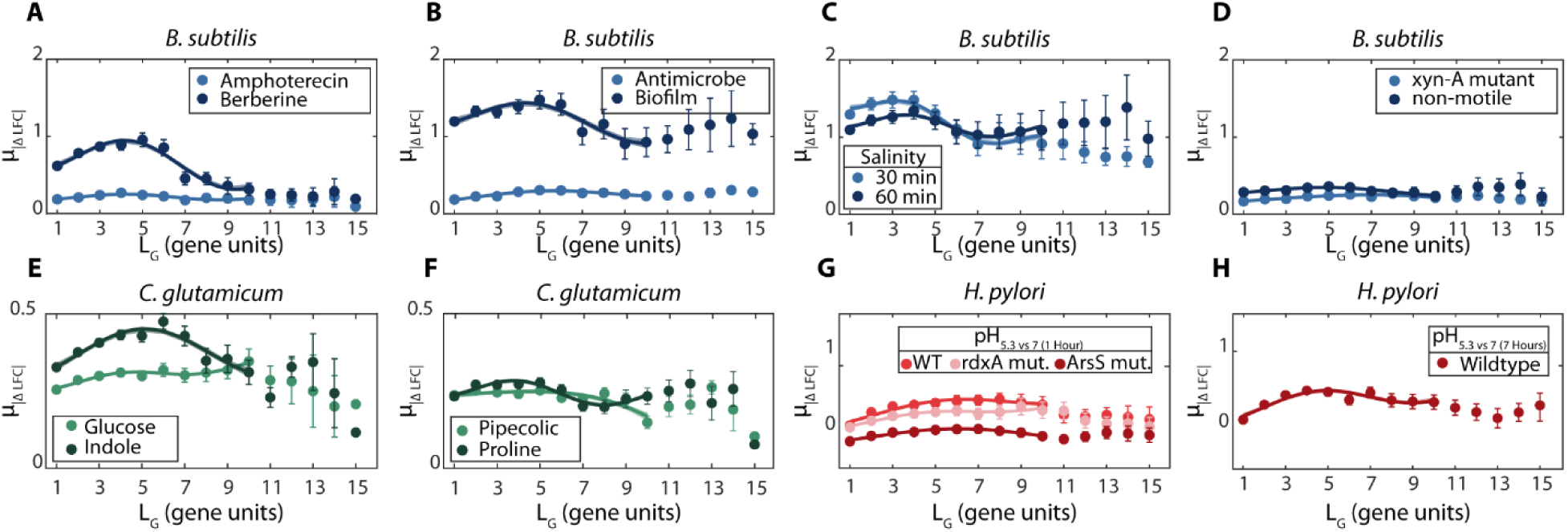
Average absolute differences in response strengths between genes in the same operon *(µ_|ΔLFC|_)* as a function of the distance between genes in the same operon, *L_G_*, in *B. subtilis, C. glutamicum, and H. pylori*. Each data is the *µ_|ΔLFC|_* of all pairs of genes of the same operon distanced by *L_G_* genes in between. Error bars are the standard error of the mean. Also shown are the best fits of functions of the form of *Eq. 1*. Their R^2^ values are shown in Supplementary Table S13-15 along with the best fitting parameter values. Data from (*43–52*). **(A-H)** Data from a specific species and conditions, listed in the figures.

While similar, the spatial frequencies differ more between species, than between different stresses in the same species. There are many possible reasons for this, including differences in how the internal TSSs of each species are arranged or in how energy is needed for the RNAP processes, among others. Meanwhile, as expected, amplitudes differ between species and stresses (Supplementary Table S13-15). Overall, the response dynamics of the operons of *E. coli* to genome wide stresses (Figures 2A-2C) is also observed in *B. subtilis*, *C. glutamicum*, and *H. pylori*.

### A stochastic model of operon dynamics regulated by internal promoters and premature terminations alone can reproduce the empirical data

We used a stochastic model of transcription in operons to test if the genome-wide behavior in Figure 2 can be explained by internal TSSs combined with changes during stresses in the rates of premature terminations of elongation. The model (Reactions S1-S7) is described in detail in the Supplementary Section “Stochastic model”.

Briefly, Reaction S1 accounts for multi-step transcription initiation, starting with RNAP binding to a TSS (primary or internal) leading to the sequential events of closed complex formation, open complex formation, and promoter escape. Reaction S2 accounts for successful elongation events that produce complete RNAs. Reaction S3 models spontaneous premature terminations of elongating RNAPs and its rate can differ between control and stress conditions. The premature terminations compete with successful elongation events. Finally, Reactions S4-S7 model various outcomes (premature terminations) due to collisions of elongating RNAPs with RNAPs in closed or in open complex formation. Premature terminations due to RNAP pausing were not included since we found no correlation between them and known pause sequences (Results section ‘Local DNA properties and other regulatory factors influence differences in the response strengths of adjacent genes’).

We simulated the stochastic model using *SGN Sim* (*53*, *54*). For each condition, operons were simulated in 2 states per condition (1000 runs lasting 10^6^ seconds each). In one state, the cells are assumed to be in the control condition and the rate constants have the values shown in Supplementary Table S16. In the other state (stress), the transcription initiation rate constants *k_cc_* and *k_oc_* of all promoters were multiplied by two, to mimic changes in transcription initiation. In addition, RNAP fall-off rates were set independently for each gene of the operon as described in Supplementary Table S16, so as to model the observed variability in RNAP fall-off rates between genes (Figure 3A).

First, we studied the effects of changing the number of internal TSSs (per gene) of an operon. From Supplementary Figure S31A, in all models, *µ_|ΔLFC|_* differs with *L_G_* according to a sine-like function, in agreement with the empirical data (Figures 2 and 5). To validate this empirically, we searched for all operons whose primary and nearest downstream promoters are separated by 5 and only 5 genes. Then, we obtained the corresponding *µ_|ΔLFC|_(L_G_)* for each stress condition and fitted *Eq. 1* to estimate the best fitting spatial frequency, which equaled ∼0.135. The genome wide spatial frequency equaled ∼0.105 (less ∼30%, Supplementary Table S1) for an average distance between the promoters of ∼4. Based on the model, we conclude that the spatial frequency differs with the distances between internal promoters in operons.

Next, we studied the effects of changing premature termination rates in order to account for changes during the stress conditions. Again, in all models, *µ_|ΔLFC|_* differs with *L_G_* according to a sine-like function (Supplementary Figure S31B).

Moreover, we tested a model without internal TSSs. In this case, the results (Supplementary Figure S31C) match instead the data in Figures 2D and 2E, as predicted. Thus, according to the model, internal TSSs and premature terminations alone can explain *µ_|ΔLFC|_*(*L_G_*) values of *E. coli*, *B. subtilis*, *C. glutamicum*, and *H. pylori*.

The model also suggests that increasing the number of internal TSSs (while maintaining the gene length of the operon constant) decreases the spatial frequency of *µ_|ΔLFC|_(L_G_)* (Supplementary Figure S31A). To test this prediction, we considered the empirical data on *µ_|ΔLFC|_(L_G_)* solely from operons with more than 3 genes (a total of 423 operons). Then, we separated them into 2 cohorts, one with operons of less than 3 TSSs and the other with the operons of 3 or more TSSs. As predicted, the spatial frequency of the best fitting function of *µ_|ΔLFC|_(L_G_)* of operons with 3 or more TSSs is higher (Supplementary Table S17). Only for ‘Media richness depletion (0.75× at 180 min)’ is this not observed. However, this condition is the one producing the weakest signals.

Moreover, from Supplementary Figure S31B, the models predict that increasing the rate of premature terminations should increase the amplitude of *µ_|ΔLFC|_(L_G_).* In agreement, these amplitudes increase with increasing concentrations of novobiocin (Supplementary Table S1), which increasingly disturbs positive supercoiling buildup and, thus, transcription elongation and likely premature termination rates (*33*).

Finally, we explored with the model the effects of internal promoter(s) whose responsiveness increases (rather than decreases) the divergence between the stress responses of upstream and downstream genes, respectively. For this, contrary to the above (where both promoters are equally activated by the stress), we next assume that only the downstream promoter is activated by the stress. This may be a realistic scenario, e.g., when one of the promoters is responsive to a stress-specific global regulator.

From Figure 30D (compared to Figure 30A), if the promoters differ in responsiveness, upstream and downstream genes will differ more in responsiveness. For all *L_G_*, *µ_|ΔLFC|_(L_G_)* is higher and only at a longer distance from the primary promoter will downstream genes exhibit the same responsiveness as the first gene of the operon (when premature terminations reduce sufficiently the differences in response strengths).

### The biological functions of genes in operons are related with *L_P_*

We investigated if the biological functions associated to the genes in operons relate with their positioning, using information on gene ontology (*55*, *56*)(Methods section ‘Gene Ontology’). We found that, on average, the closer two genes are to each other in an operon, the more likely it is that they are involved in the same biological processes (Supplementary Figure S32). This conclusion reinforces past findings (*9*) using other classification methods.

We also studied the number of genes between consecutive promoters with specific biological functions as a function of the distances from the closest upstream and the closest downstream TSSs (Table 1). We considered gene ontology (GO) terms that are direct descendants (child) of the general GO term GO:0008150 “biological process” (*62*). Table 1 shows the 8 (of 23) terms considered (rows 1 to 8). The other terms were not considered since we found less than 100 genes with such feature(s).

**Table 1.** Distances from the closest upstream TSSs and the closest downstream TSSs of genes with specific biological functions. Shown are the average (µ) and the standard deviation (STD) of these distances between consecutive internal TSSs (*L_P_*, in gene units) of gene cohorts sharing a higher order biological function (based on the gene ontology classification (*55*, *56*)). Also shown are the number of genes of each cohort. Gene functions are based on the gene ontology classification in (*55*, *56*), except for cohorts 10 to 11, which involve ‘all genes’ in operons and ‘essential genes’, which are based on the classification in (*57*). For additional information, see the distribution of operon genes lengths in Supplementary Figure S12 (top left), with a median of 3. In addition, we show the *p*-value obtained from a KS test comparing the distributions of *L_P_* values of the various operons with specific biological functionalities against the distribution of all *L_P_* values of *E. coli*’s genome. For *p*-values smaller than 0.05, we discard the null hypothesis that the two distributions come from the same distribution.

| Functionality | | $\mu(L_P)$ | STD($L_P$) | Genes in operons |
| --- | --- | --- | --- | --- |
| 1 | Positive regulation of biological process ** | 0.6 | 1.22 | 70 |
| 2 | Localization * | 1.09 | 1.31 | 449 |
| 3 | Metabolic process | 0.92 | 1.42 | 1225 |
| 4 | Regulation of biological process *** | 0.55 | 0.99 | 291 |
| 5 | Response to stimulus | 0.86 | 1.37 | 475 |
| 6 | Biological regulation *** | 0.59 | 1.00 | 358 |
| 7 | Cellular process | 0.95 | 1.38 | 1598 |
| 8 | Negative regulation of biological process *** | 0.36 | 0.63 | 110 |
| 9 | All genes not in 1-8 | 0.93 | 1.38 | 597 |
| 10 | All genes in operons | 0.96 | 1.40 | 2262 |
| 11 | Essential Genes * | 0.59 | 0.89 | 222 |

\* $P \leq 0.05$ . \*\* $P \leq 0.01$ . \*\*\* $P \leq 0.001$ .

From Table 1, 6 of the 9 cohorts have distributions of *L_P_* values that statistically differ from the genome-wide distribution. Moreover, the mean, *µ(L_P_)*, and the standard deviation, *STD(L_P_)*, differ widely between the cohorts of genes with specific biological functions. *E.g.*, genes associated with ‘regulation of biological processes*’* (rows 1, 4, 6, and 8) have small *µ(L_P_)* = 0.52, while the remaining cohorts have *µ(L_P_)* = 0.95, similar to the average of other genes in operons (rows 9 and 10). This suggests that the distance of genes in operons from the closest upstream promoter may be influenced by their biological function.

Finally, we considered 236 ‘essential’ genes, i.e., genes that, if deleted, cells do not survive in optimal control conditions (*57*). Given this definition and the existence of premature terminations during elongation, essential genes in operons should be relatively close to their upstream promoters (to ensure strong expression). In agreement with this prediction, the *µ(L_P_)* of essential genes equals 0.59, which is much smaller (statistically) than the genome-wide average (Table 1). This suggests that essential cellular processes may be amongst those that need to be more quickly and efficiently adjusted during stresses.

Finally, also in agreement with our hypothesis, this appears to contribute to higher mean transcription rates in essential genes, in all conditions. Specifically, essentials genes located in operons have higher normalized read counts with a mean of 846 and STD of 3117, while other genes have a mean of 141 and STD of 658 (as measured by the transcripts per million method, TPM (*58*)) under all stresses. This agrees with protein measurements in cells in optimal conditions (*32*).

## Discussion

Many open questions remain about the general principles governing the formation and evolution of operons. Studies have characterized several individual operons and identified clear phenotypic advantages of placing specific genes under the control of the same promoter(s), due to related biological functions (*6*, *10*, *12*). For example, the *rplKAJL-rpoBC* operon codes for proteins RplK, RplA, RplJ, and RplL, which are all 50S ribosomal subunits (*59*). The operon further codes for RpoB and RpoC, which are the subunits β and β’ of RNAP (*12*) (*59*). Having the production of any of these proteins under the control of another operon would probably decrease the efficiency with which *E. coli* forms RNAPs and ribosomes. Such need for producing sub-units of complex proteins at a similar time, location, and quantities provides one general explanation for why genes coding for protein subunits are integrated into the same operons.

Other studies have identified benefits of having internal promoters in some operons (*60*). For example, *rplKAJL-rpoBC* has three promoters, which allow, in some conditions, to produce the protein pairs (RplK and RplA), (RplJ and RplL), and (RpoB and RpoC) with relatively distinct dynamics (*61*). This may be advantageous when, e.g., ribosomes and RNAPs should not be produced at the same rate (*62*). Meanwhile, decreasing the physical stability of mRNAs and increasing failures in transcription elongation might limit the size of operon lengths (*63*). Thus, we hypothesized that, in some conditions, some internal promoters may assist in harmonizing the production of upstream and downstream genes, while also allowing diverging their expression levels in other conditions.

Here, we studied the behavior of genes in operons in the context of genome wide perturbations that, by targeting RNAP and gyrase (*22*, *23*), affect both primary, as well as internal promoters of most operons, quasi-synchronously. We analyzed this data on the response dynamics while accounting for information on the composition of the operons of *E. coli*, whose genome is likely the most dissected one. This allowed accessing the degree to which operon lengths and internal promoters can be correlated to the dynamics of RNA production within operons.

We found that differences in response strengths between genes in the same operon are well matched by relatively ‘smooth’ wave (sine) functions of the distance between the genes. This was unexpected, since they emerge from combining the data on changes in gene expression from all operons, which differ in many ways (e.g. in the number, positioning and response strengths of internal promoters, σ factor preferences, transcription factor regulators, etc.). Nevertheless, the wavelike patterns emerge from the activity of internal TSSs to enhance the expression of their downstream genes, combined with relatively frequent premature transcription terminations of elongating RNAPs, which are generated by several processes. Contrarily, in operons lacking internal TSSs (and in transcription units), the differences in responsiveness of the component genes, as a function of the distance between them, are better matched by lines with positive slope, in agreement with the occurrence of tangible, genome-wide premature termination rates of elongating RNAPs. Finally, from recently published data, we observed the same phenomenon in *E. coli* under various other stresses, and in three other bacterial species (one gram-negative and two gram-positive). As such, this may be a prevalent evolutionary constraint across bacterial species.

Internal promoters should be able to exert complex, condition-depending influences on operon dynamics. First, they can enhance the response strengths of downstream genes, with the enhancement differing with the strength and positioning of the promoter along the operon. Second, they can combine their actions with the actions of other promoters in the same operon. Third, given that open complex formations and promoter escapes are rate-limiting steps that can take longer time than elongation (*64*, *65*), internal promoters should generate substantial numbers of collisions between RNAPs bound to them and RNAPs elongating from upstream promoters (as supported by the kinetics of the synthetic constructs here reported). Moreover, the dynamics of these collisions should be adaptive and evolvable, given the adaptability and evolvability of transcription initiation kinetics (*64*, *65*). These observations were reproducible by a simple stochastic model of transcription assuming RNAP elongation events emerging from a primary as well as internal promoters, along with frequent RNAP premature termination events (due to collisions between sitting and elongating RNAPs and due to other events).

At least three results support the hypothesis that the number and positioning of internal promoters can fine tune transcription elongation kinetics in operons during the genome-wide stresses. First, the cohort of operons whose primary and nearest downstream promoters are separated by 5 genes (the genome-wide average distance is ∼4) have a best fitting spatial frequency for *µ_|ΔLFC|_(L_G_)* that is ∼30% higher than the genome-wide value. Second, consistently, operons with more TSSs ( ≥ 3) nearly always have higher spatial frequency (*f*) than cohorts with less TSSs (<3) (Supplementary Table S17). Third, in all stresses, the operons subject to self-TF regulation exhibited higher spatial frequency than the genome-wide average (Supplementary Table S5). To study why, we determined the average distance between the primary and nearest downstream internal promoter in those operons. We found that it equals ∼3.1, which is also smaller than the genome wide average.

Based on the above, assuming that the relationship between spatial frequency and downstream promoters positioning is valid for the three other organisms studied, from their spatial frequencies (Supplementary Tables S13, S14, and S15), we infer that the genome distances between primary and nearest downstream promoters in *B. subtilis* are similar to *E. coli*. Meanwhile, they are likely smaller in *H. pylori*, and bigger in *C. glutamicum*. Future mapping of their promoters in operons should allow testing this prediction.

Next, we studied which processes can best explain the changes in premature terminations of RNAP elongation. First, we provided evidence that positive supercoiling buildup can partially explain some changes in premature terminations, in that genes known to be highly responsive to positive supercoiling buildup in standard growth conditions are related to the strongest differences in response strengths along operons. Additionally, in poor media conditions we also observed low cellular ATP levels which, similarly to novobiocin, can hamper Gyrase activity (*23*). Second, we used a synthetically engineered pair of promoters in non-overlapping tandem formation, to show that collisions between RNAPs elongating from an upstream promoter with RNAPs bound to a downstream promoter can cause high rates of premature terminations. Moreover, this phenomenon differs in rates of occurrence with the nature and the strength of the genome-wide stresses studied. Similar events likely influence the RNA productivity of natural operons with internal promoters. Third, we showed that sequences commonly found at 3’ ends of RNA, σ factor preferences of the internal promoters, small regulatory RNAs, (p)ppGpp, and transcription factor binding sites are also related to high premature termination rates.

Future studies should allow discerning the contribution of each of the above phenomena to different operons and stress conditions. Moreover, it should also be feasible to investigate the influence from changes in global translation rates during the stresses, as these may affect the stability of RNAP elongation complexes, leading to premature termination (*66*, *67*). Also, the changes in the ATP levels likely have indirect effects on cellular physiology, which might be influential in ways other than the changes in supercoiling levels here considered. Another factor not considered is the changes in the local 3D structure of the DNA, which likely differs between stresses. These efforts could be assisted by the engineering of long synthetic operons in the chromosome, or *in vitro*, where the number of unknown variables is likely smaller, and where various parameters could be more easily controlled without disturbing cellular homeostasis.

So far, the main biological function associated with internal promoters has been enhancing divergency between upstream and downstream genes, in response to specific signals (*19*), including antibiotics (*68*). However, under the genome-wide stresses studied here, overall, they decreased divergency, counteracting the effects of premature terminations. This and the strong correlation between the internal promoters’ response strength and number of downstream genes prior to other internal promoters (Figure 2F) suggest that premature termination rates may have partially shaped the number, positioning, and strength of internal promoters in operons.

Another way the gene network of *E. coli* may have been shaped by premature terminations is the number and location of TTSs. There are only 174 known TTSs associated with operons in *E. coli*, while operons contain more than 2500 genes (*12*). However, a tangible premature termination rate could allow more distanced genes to become uncorrelated, even in the absence of a TTS. In agreement, adjacent genes in the DNA, but located in different operons, that are more distanced from each other are also less likely to have a TTS in between (Supplementary Figure S1D). This implies that more distanced genes have their activity made independent by natural premature terminations.

The findings reported here could be of use to the Bioindustries, including in Synthetic Biology, and to Biomedicines. Rearranging the positioning of genes and TSSs in operons may allow fine-tuning natural transcriptional programs of stress response. I.e., re-positioning of genes in operons should allow tuning their response strengths to certain stresses, which could alter certain cellular processes, in order to, for example, minimize resources consumption, and thus improve the yield and sustainability of bioreactors. Similarly, adjustments of gene positioning in operons and the use of internal TSSs could also become useful for tuning the kinetics of synthetic genetic circuits. Finally, if bacteria are placing vital genes (e.g. essential ones) immediately downstream to primary and internal promoters, to ensure their responsiveness to genome-wide stresses, we could potentially use this to search for specific target genes for antibiotics.

Overall, our results provide explanations for the naturally evolved lengths of prokaryotic operons, and for the number, strength, and location of their internal promoters. The findings may be of use to predict and dissect the same features in the operons of other bacteria.

## Materials and Methods

### Bacterial strains and genome-wide stresses

*E. coli* (MG1655 strain) is our model organism because, first, its TFN is likely the best mapped one (*12*), which facilitates determining the influence of TF interactions in genome-wide stresses (*23*). Moreover, sequence-dependent phenomena, e.g., transcriptional pauses and other elongation-related events are relatively well known (*13*, *69*). Finally, many biological functions of most proteins have been studied at a genome-wide level (*55*, *56*).

To study how the genome wide stress responses of *E. coli* are shaped by the internal structure and components of its operons, we relied on existing annotations, e.g., on the location of internal TSSs, σ factor preferences, etc. (*12*). While, for *E. coli*, the accumulated knowledge is relatively extensive, it is not yet complete and may contain biases. As these annotations continue to improve, it should be possible to further improve the level of precision of our results.

RNA-seq data on genome-wide stresses is partially curated from (*22*, *23*) and partially produced for the present study. In all cases, changes in gene expression were measured by RNA-seq by estimating the genes’ Log fold changes (LFC) relative to RNA-seq when not adding novobiocin, not diluting the medium, and not adding rifampicin, respectively. A total of 4573 unique combination of genes separated by a certain distance was obtained for genes in the same operon (with > 1 genes). For novobiocin, we have rejected 1015 pairs (unknown differences in LFC). So, we have considered only 3558 pairs of genes, for whose LFC is known and calculated the absolute Delta Fold change between them. Same as novobiocin, we have rejected 1631 gene pairs for Media-dilution as one we don’t know at least one of their LFC, and only 2942 gene pairs were considered (FC of 653 genes is unknown). For rifampicin we have considered 3897 pairs of genes and rejected 676 pairs.

We also collected data on protein levels by flow-cytometry from cells subject to novobiocin stresses (50 and 500 ng/μL), respectively (methods section ‘Flow cytometry’). For all stresses, we measured the optical density at 600 nm (OD_600_ nm) to monitor the cell growth, using a Bioteks Synergy HTX Multi-Mode Reader.

For simplicity and readability, in the figures, the concentrations are all in μL^-1^. Finally, in Supplementary Table S19, we list all variables used in the study, and their definitions.

### Estimating genome-wide differences in reads between the start and end of genes in operons

To demonstrate the existence of substantial *premature* termination rates during elongation, even in the absence of TTSs, we obtained read counts, ‘*reads*’, of the starting as well as of the ending DNA coding ‘regions’ of each gene, in each operon (including genes coding for small RNAs. The start and end regions of a gene are defined by the corresponding RNA reads that are aligned to the genomic coordinates of the gene’s start and end nucleotide, respectively (annotated in sequence ID: NC_000913.3.). These regions and corresponding counts are illustrated in Figure 3A. For this, we used the MATLAB ‘featurecount’ function, and then normalized all read counts using the Transcripts per million method (TPM) (*58*).

Next, for each gene, we applied a t-test to evaluate if the read counts from each of the three biological replicates (for each stress condition) differ statistically. Then, we estimated the normalized premature termination rate, *r_f_*, of each gene in position *i* of operon *j* as follows:

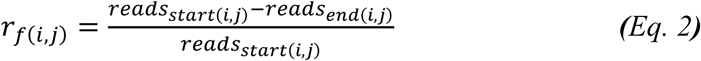

From this, the operon-wide average rate of premature terminations, *R_f_*, as a function of the position of the genes is given by:

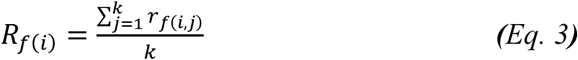

where *k* is the total number of genes in each position in the genome.

Finally, we removed from this analysis the 12 genes (in operons) whose coding regions are reported in RegulonDB to have TTSs (*12*), because we aimed to quantify termination rates *not* caused by TTSs.

### RNA-seq

#### a) Sample preparation

The RNA-seq data used for the four media dilutions and for the condition with novobiocin at 50 ng/μL were published in (*22*) and (*23*), respectively.

From (*22*), starting from standard LB medium, we produced tailored media, denoted as ‘LB_1.0×_’, ‘LB_0.75×_’, ‘LB_0.5×_’, and ‘LB_0.25×_’. Their composition for 100 mL (pH of 7.0) are, respectively: (LB_1.0×_) 1 g tryptone, 0.5 g yeast extract and 1 g NaCl; (LB_0.75×_) 0.75 g tryptone, 0.375 g yeast extract and 1 g NaCl; (LB_0.5×_) 0.5 g tryptone, 0.25 g yeast extract and 1gNaCl; and (LB_0.25×_) 0.25 g tryptone, 0.125 g yeast extract and 1 g NaCl.

To collect the data in other conditions (listed in Figures 2A-C), cells in the mid-exponential growth phase (OD_600_ = 0.3) were subject to, respectively, 100 ng/μL novobiocin, 500 ng/μL novobiocin, and 2.5 ng/μL rifampicin. Two hours later, for each condition (including a control strain, not subject to antibiotic stress), samples were collected from 3 independent colonies. This time lag reduces cell-to-cell diversity due to, e.g., different antibiotic absorption times. After collecting the samples, 5 mL of the culture was immediately treated with a double volume (10 mL) of RNA protect bacteria reagent (Qiagen, Germany) for 5 minutes at room temperature, to prevent RNA degradation. Next, treated cells were pelleted and frozen at –80 °C overnight. The next morning, total RNA was extracted using the RNeasy kit (Qiagen, Germany).

#### b) Sequencing

Extracted RNA was treated twice with DNase (Turbo DNA-free kit, Ambion, USA) and quantified using Qubit 2.0 Fluorometer RNA assay (Invitrogen, Carlsbad, CA, USA). The total RNA quality was determined using a 1% agarose gel stained with SYBR Safe (Invitrogen, Carlsbad, CA, USA), where RNA was detected using UV in a Chemidoc XRS imager (Biorad, USA). RNA integrity was measured by the Agilent 4200 TapeStation (Agilent Technologies, Palo Alto, CA, USA).

RNA library preparations, sequencing, and quality control analysis of sequenced data were conducted at GENEWIZ, Inc. (Leipzig, Germany). In detail, ribosomal RNA depletion was performed using Ribo-Zero Gold Kits (Bacteria probe) (Illumina, San Diego, CA, USA), while the RNA sequencing library was prepared using the NEBNext Ultra RNA Library Prep Kit.

The sequencing libraries were multiplexed and clustered on one lane of a Flowcell, which was loaded on an Illumina NovaSeq 6000 instrument. In both instruments, the samples were sequenced using a single-index 2×150 Paired-End (PE) configuration. Image analysis and base calling were conducted by the NovaSeq Control Software v1.7 (Illumina NovaSeq). The raw sequence data (.bcl files) was converted into “fastq” files and de-multiplexed using Illumina bsl2fastq v.2.20. One mismatch was allowed for index sequence identification.

#### c) RNA-seq data analysis pipeline

We analyzed RNA-seq data in four steps:

*Step 1:* RNA sequencing reads were trimmed to remove possible adapter sequences and nucleotides with poor quality using Trimmomatic v.0.39 (*70*).

*Step 2:* To generate BAM files, trimmed reads were mapped to the reference genome *E. coli* MG1655 (NC_000913.3) using the Bowtie2 aligner v.2.3.5.1 (*71*).

*Step 3:* Unique gene hit counts were calculated using ‘featureCounts’ from the ‘Rsubread’ R package (v.2.8.2)(*72*). Genes with less than 5 counts in more than 3 samples, as well as genes whose mean counts were smaller than 10 were removed from further analysis.

*Step 4:* The DESeq2 R package (v.1.34.0) (*73*) was used to calculate the log_2_ fold changes (LFC) of RNA read counts between conditions. Also, it calculates the corresponding *p*-values, using Wald tests (function ‘nbinomWaldTest’).

### Flow cytometry

We used an ACEA NovoCyte Flow Cytometer to measure cell fluorescence. Cells were diluted (1:10000) in 1 mL of PBS solution, and then vortexed for 10 seconds. For each condition, we performed three biological replicates and collected data from 50 000 cells from each replicate using the Novo Express software (ACEA Biosciences Inc.). The flow rate was set to 14 µL/minute.

For detecting YFP, we used a blue laser (488 nm) for excitation and the fluorescein isothiocyanate detection channel (FITC-H) (530/30 nm filter) for emission. We set a core diameter of 7.7 μM and a PMT voltage of 600. To remove interference from particles, a detection threshold of 5000 was set in FSC-H. Abnormal cells were identified using the nonparametric outlier detection method ‘Tukey fences’ (*74*). The method establishes boundaries beyond the first and the third quartiles by 1.5 times the interquartile range. Data beyond these boundaries are classified as outliers

### Statistical tests

#### a) Curve and line fitting and selecting best fitting models

To model the empirical relationship between the distances between genes in the same operon, *L_G_*, and the mean absolute differences in their response strengths, *µ_|ΔLFC|_*, we used a nonlinear model (Eq. 1). The model has five unknown parameters to be estimated from the data (*A_0_, A, L_G_, B, and f)*.

To estimate, we used the MATLAB 2019 ‘fit’ function, to which we provided the empirical values of *L_G_* and *µ_|ΔLFC|_* in order to best fit function (Eq. 1). We repeated the fits 100 times per stress condition, to eliminate any bias from the random initial conditions assumed by the ‘fit’ function. We tried increasing the number of runs, but it did not change the solutions.

Noteworthy, we used the same protocol to fit linear regression models. In both cases, to choose the best fit, first, we selected the fit(s) with the highest R^2^ value. If only one model remained, we kept that model as the best fitting one. Else, to select from the remaining models, we calculated the Bayesian information criterion (BIC) and selected the model with both lowest BIC. Finally, we further tested the solution by applying the Akaike information criterion (AIC) and selected the model with the lowest AIC. In all cases, the AIC and BIC made the same selection.

#### b) 2-sample t-test and 2-sample KS-test

We used 2-sample t-tests to investigate the null hypothesis that two samples come from normal distributions with equal means and equal, but unknown variances. The MATLAB function *ttest2* was used for this. We rejected the null hypothesis at a 5% significance level.

We also used 2-sample Kolmogorov-Smirnov (KS) tests using the MATLAB function *kstest2*. This function returns a decision for the null hypothesis that the data from two data sets are from the same continuous distribution. We rejected the null hypothesis at a 5% significance level.

#### c) Standard Error of the Fold change

The standard error of the fold changes was calculated using the standard formula:

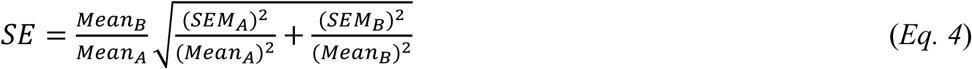

where ‘*A’* and ‘*B*’ represent each of the two sets of data whose fold change is being analyzed.

#### d) Autocorrelations

We calculated the spatial autocorrelation between genes in operons as a function of their distance from the primary (most upstream) promoter. For this, we used the MATLAB’s function ‘xcorr’ with unbiased autocorrelation. This unbiased method adjusts for the decreasing of the number of overlapping data points as the distance increases, so as avoid underestimating the correlation values at larger distances. The process was repeated for all operons. Then, we calculated the mean normalized autocorrelation as a function of *P*, for each stress condition. Operons with one or more genes with an unknown log fold change were removed from the analysis.

### Promoters AT-richness

We considered all promoters identified in RegulonDB (*12*). We then discarded 37% of them since their DNA sequence is not available. Of the remaining 1057 promoters, we consider their DNA sequence starting from 60 nucleotides upstream of the TSS up to 20 nucleotides downstream of the TSS.

Assuming the TSS in position +1, from the nucleotides in positions –60 to +20 (*12*, *75*), we calculated the fractions of A, C, G and T, respectively. The AT-richness of the promoter was estimated by summing the fractions of A and T in this region.

### Identifying premature terminations and pause sequences

Premature termination-prone and pause-prone sequences were searched using the MATLAB *localalign* function. From the whole genome sequence of *E. coli* MG1655 strain in FASTA format, we searched for specific pause sequences reported in (*69*) and for premature termination sequences, such as homopolymetric A’s or T’s, reported in (*76*).

To eliminate randomness or bias from having different starting points during the optimization process, a gap open penalty of 1 has been counted for each misaligned read and a maximum of 4000 alignments have been performed. Finally, we counted the number of aligned sequences for different degrees of freedom (i.e. number of errors allowed), namely, for 0 and 1 degrees. Finally, we normalized their rate of occurrence by the total number of operons.

### Classification of pairs of genes as ‘highly sensitive to positive supercoiling buildup’

Reference (*23*) used the RNA-seq data on the stress caused by novobiocin at 50 ng/µL to identify which genes are highly sensitive to positive supercoiling buildup. Namely, these were the genes whose *absolute* log fold change with the stress was above 0.4 (for a *p*-value < 0.05). Thus, initially highly/weakly expressing genes in the control condition were strongly responsive if their expression under the stress condition was weak/high (above the threshold), respectively. Based on that, we then used the following rule to classify pairs of genes in the same operon as ‘highly sensitive to positive supercoiling buildup’. Specifically, a pair of genes was highly sensitive to positive supercoiling buildup if either of the two genes, or any gene located between them, is sensitive. All other pairs of genes were classified as non-sensitive.

### Gene ontology

To study how Gene Ontology (GO) relates to the operons’ spatial organization, we determined all GO biological processes terms that each gene in *E. coli* is known to be involved in (*55*, *56*). Next, for all 4573 pairs of genes in the same operon, we extracted the number of GO terms that the two genes have in common. Also, we calculated the percentage of common GO terms for each pair. This percentage, *P_c_*, equals the ratio between the number of common terms between two genes, *n_c_*, and the number of possible combinations (*N_i_×N_j_*), given the number of GO terms of each of the two genes, *N_i_* and *N_j_*, respectively (*55*, *56*):

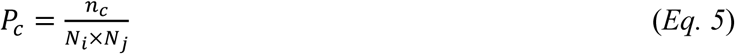

We discarded 29% of the gene pairs, since at least one of the genes was not associated with any GO term.

As a null model, we performed 1000 random shuffles of the pairs that each gene belongs to. This created 4573000 pairs, but then we removed random pairs having one or more genes without GO term associations. We also removed pairs with genes from the same operon. From the remaining pairs, we calculated the percentage of common GO terms. Overall, 37% of all random pairs were discarded.

### Cellular ATP levels

QUEEN-2m cells (kind gift from Hiromi Imamura (*77*) were grown in ‘LB_1.0×_’, ‘LB_0.75×_’, ‘LB_0.5×_’, and ‘LB_0.25×_’ (methods section ‘RNA-seq’). We tracked their ATP levels (Supplementary Figure S17) using a Biotek Synergy HTX Multi-Mode Reader. The solution was excited at 400 nm and emission was recorded at 513 nm. Similarly, the solution was re-excited at 494 nm and emission was recorded at 513 nm. The ratio between the 513 nm emission intensities at the two excitation wavelengths, which was denoted as ‘400ex/494ex’, is used as a proxy for cellular ATP levels as proposed in (*77*).

## Acknowledgments

We would like to acknowledge M. Bahrudeen and B. Almeida for suggestions regarding the statistical analysis. We also thank all referees and HT Jacobs for their valuable suggestions.

## Funding

Work supported by

The Jane and Aatos Erkko Foundation [10-10524-38 to A.S.R.]

The Sigrid Jusélius Foundation [A.S.R.]

Suomalainen Tiedeakatemia [R.J. and S.D.],

Finnish Cultural Foundation [00222452 to I.S.C.B., and 00210818 to C.S.D.P.]

Tampere University Graduate Program [V.C.]

EDUFI [TM-21-11655 R.J.].

Academy of Finland [349116 to J.M.].

The funders had no role in study design, data collection and analysis, decision to publish, or preparation of the manuscript.

## Author contributions

Conceptualization: ASR, RJ, CP, SD

Methodology: RJ, ASR, CP, SD, IB

Investigation: RJ, VC

Visualization: RJ, IB, JM

Supervision: ASR, RJ

Writing—original draft: ASR, RJ, SD, IB

Writing—review & editing: ASR, RJ, CP, JM, SD, IB

Funding acquisition: ASR, RJ, JM

Resources: RJ, SD

Project Administration: ASR

Formal Analysis: RJ, IB, CP

Software: RJ, IB, CP

Validation: RJ, SD, IB, CP Data Curation: RJ

## Competing interests

Authors declare that they have no competing interests.

## Notes

### Competing Interest Statement

The authors have declared no competing interest.

### Summary of Updates

New sections have been added following the suggestions of the referees. Some of the main manuscript figures have been moved to supplementary files.

