## Supplementary material for "Dynamics of bacterial operons during genome-wide stresses is influenced by premature terminations and internal promoters"

Rahul Jagadeesan et al

**This PDF file includes:**

Supplementary Text
Figs. S1 to S32
Tables S1 to S19

### Supplementary Text

#### Sites reported to be prone to premature transcription terminations

Recently, in reference (*36*) in the main manuscript, it was reported that many coordinates of RNA 3´ ends in 5´ untranslated regions (UTRs) are prone to Rho-dependent and factor-independent premature transcription terminations. The information on each of these sites is available in the form of the specific nucleotide positioning in the DNA of E. coli MG1655 (reference genome NC_000913.3, wild type).

We tested if these sites influenced our measured normalized premature termination rates ($R_f$). For this, we considered the 429 *pTTSs* reported in (*36*) to be located in operons. Then, we investigated if the mean differences in response strengths ($\mu_{|\Delta LFC|}$) between adjacent genes in the operons (i.e., for $L_G = 1$) are locally influenced by *pTTSs*.

For this, first, we studied $\mu_{|\Delta LFC|}(L_G = 1)$ of pairs of adjacent genes in operons when at least one of the genes (or a region between the genes) contains at least one *pTTS* (Supplementary Figure S18). We found that $\mu_{|\Delta LFC|}(L_G = 1)$ is consistently ~40% higher in pairs of genes with a *pTTS*, in all stress responses (compared to all other pairs). Second, we studied if $R_f$ also correlate with *pTTSs* locations. We found that $R_f$ of genes in operons with at least one *pTTS* in their coding region is higher (Supplementary Figure S19).

In summary, the *pTTSs* reported in reference (*36*) in the main manuscript enhance mean absolute differences in response strengths between adjacent genes in operons by increasing premature terminations of elongating RNAPs.

#### σ factor preferences, sRNA regulation, (p)ppGpp, and transcription factor binding sites

Several regulatory mechanisms of gene expression in *E. coli* influence the dynamics of expression of operons. We evaluated how some of these mechanisms influence differences in response strengths to genome-wide stress between adjacent genes in operons.

First, we studied the influence of promoters' σ factor preferences since σ factors are expected to exert control on stress-specific responsiveness. For this, we estimated the mean absolute differences in response strength ($\mu_{|\Delta LFC|}$) between adjacent genes in operons that are separated by an internal promoter whose σ factor preference differs from the preference of its upstream promoter. Visibly, different σ factor preferences in promoters of the operon can cause differences $\mu_{|\Delta LFC|}$ between adjacent genes (Supplementary Figure S20). $\sigma^{38}$ factor preferences cause the highest differences between adjacent genes, but this is likely due to the nature of the stresses applied.

We also investigated regulatory non-coding RNAs (sRNA's), by comparing mean differences in response strengths between adjacent genes in operons ($\mu_{|\Delta LFC|}(L_G = 1)$) when at least one of the two genes has a known sRNA binding site region (Supplementary File S2**)**. From Supplementary Figure S21, when these sites are present, $\mu_{|\Delta LFC|}$ is higher.

Similarly, we examined the potential influence of sensitivity to (p)ppGpp, since a large number of genes in operons has been reported to be sensitive to (p)ppGpp (listed in Supplementary File S3 using information in (*12*)). From Supplementary Figure S22, the $\mu_{|\Delta LFC|}$ between adjacent genes when one or both are sensitive to (p)ppGpp is higher than when neither gene is sensitive.

Finally, we observed that the presence of internal TFs binding sites (TFBSs, listed in Supplementary File S4) also enhances $\mu_{|\Delta LFC|}$ between adjacent genes in operons (Supplementary Figure S23). Potentially, these three forms of regulation may enhance $\mu_{|\Delta LFC|}$ between adjacent genes by increasing the rates of premature termination of RNAPs elongating from upstream promoters (e.g. due to collisions with the regulatory factors or other forms of physical interference with the elongation process). Another possibility is the enhancement of the expression of the downstream genes, by recruiting RNAPs to those regions.

Overall, we conclude that differences in response strength between adjacent genes in operons are sensitive to these natural, local, sequence dependent gene regulatory mechanisms of *E. coli*.

**RNAP pausing sequences, RNA degradation rates, distances from the origin of DNA replication, TSSs in the opposite DNA strand, and promoters AT-richness**

We investigated if we could relate premature termination rates to specific DNA sequences. First, we searched for 14 DNA sequences (12 nucleotide each) (Methods section 'Identifying premature terminations and pause sequences', main manuscript) known to enhance the propensity for transcriptional pausing (ref (*69*), main manuscript), since these events could promote premature terminations (e.g. by enhancing RNAP collisions). Also, we searched for homopolymetric tracts of A's or T's with 9 nucleotides each, as they can promote RNAP slippage and premature terminations (ref. (*76*), main manuscript) Their rate of occurrences in the DNA of *E. coli* (operon regions) are shown in Supplementary Table S10.

We tested if the frequency of occurrence of each of the sequences correlates with the average normalized rates of premature terminations, $R_f(P)$. From Supplementary Figure S24, we found no correlation in any case.

Next, we obtained information on TSSs known to be located on the opposite strand of regions coding for operons (ref. (*12*), main manuscript), since these sites could potentially enhance RNAP collisions ((*78*, *79*), main manuscript). However, we did not find such TSSs in sufficient numbers to test if they are locally influential (only 18 are listed and most were located at the primary promoter regions and, thus, one cannot test if they influence $\boldsymbol{\mu_{|\Delta LFC|}}$ of adjacent genes).

We also considered the possibility that $\mu_{|\Delta LFC|}(L_G)$ in Figure 2D and 2E in the main manuscript could be influenced by differences in RNA degradation rates of genes in the same operon. However, we found no correlation between the two features (Supplementary Figure S25). Similarly, we obtained differences in the degradation rates of RNAs coded by adjacent genes from ref. (*80*) of the main manuscript and compared with the corresponding $\mu_{|\Delta LFC|}$. From Supplementary Figure S26, under no stress are differences in RNA degradation rates statistically correlated with the differences in response strengths, $\mu_{|\Delta LFC|}$, between adjacent genes in same operon.

Next, we considered whether DNA replication could influence premature termination rates during the genome-wide stresses. Thus, we assessed if the average $\mu_{|\Delta LFC|}$ of adjacent genes in the same operons correlates with the distance of the operon from the origin of DNA replication. Since internal promoters compensate for premature terminations, we only considered pairs of genes without internal promoters in between. From Supplementary Figure S27, using t test statistic, in general, we failed to reject the null hypothesis that the line does not differ from a horizontal line (except for the rifampicin stress).

Finally, we studied if the AT-richness using data from ref. (*81*) in the main manuscript (see also Methods section 'Promoters AT-richness' in the main manuscript) of internal promoters relate with differences in response strength between their (adjacent) upstream and downstream genes. From Supplementary Figure S28, we found no statistical correlation between the internal promoters AT-richness and the |ΔLFC| between their upstream and downstream genes, except in the stress caused by rifampicin (as above). These results may suggest that rifampicin's efficiency to prevent RNAP escape from a promoter is weakly, but tangibly, influenced by the AT-richness of the promoter.

**Stochastic model**

Models of prokaryotic single gene expression (*4*, *82–85*) generally assume that transcription occurs when an RNAP finds a promoter (and corresponding TSS) and initiates transcription. Once escaping the promoter, the RNAP elongates to assemble the RNA coded by the gene. Finally, the RNA is released and the RNAP unbinds from the DNA (*4*, *82*, *83*, *85*).

The initiation consists of a closed and open complex formation, and of promoter escape towards elongation. These events can be delayed or stalled by promoter locking due to positive supercoiling buildup, whose unlocking requires Gyrase intervention (*4*, *23*, *86–88*).

Several events can occur during elongation as well including transcriptional pausing, misincorporation and editing, pyrophosphorolysis, and premature terminations (*64*, *89–94*). Also, it is possible for RNAPs to collide, particularly when one is paused or bound to a TSS of an promoter in complex formation (*78, 79, 95, 96*).

Based past models for individual genes (*65*, *83*, *97*) and given how $\mu_{|\Delta LFC|}$ changes with $L_G$ (Figure 2 in the main manuscript), here we propose a general, genome-wide model of the dynamics of the responses of operons to the genome-wide stresses. The model considers a promoter, i.e., its position in the operon, along with the genes and promoters that are upstream and downstream of it. This allows defining a set of reactions controlling the transcription dynamics of that promoter that account for the other factors (e.g., elongating RNAPs from the upstream promoters), which can then be generalized using a matrix-like formalism.

Finally, to avoid modelling the processes at the nucleotide level, rate constants are tuned based on the frequency of events within the operons that best explain our measurement data.

Consider a model promoter $P_j$, located in the model operon $k$, In general $P_j$ can range from 1 to $n_{k,j}$ which is the number of promoters in operon $k$. This promoter controls the expression of its downstream genes, from gene i to gene $n_{k,i}$, where $n_{k,i}$ is the total number of genes in operon $k$.

We model transcription initiation at the promoter $p$ in operon $j$ as:

$$\mathrm{Pro}^{i,P_j} + R \xrightarrow{K^{P_j}{}_{cc}} \mathrm{CC}^{i,P_j} \xrightarrow{K^{P_j}{}_{oc}} \mathrm{OC}^{i,P_j} \xrightarrow{K^{P_j}{}_{esc}} \mathrm{EC}^{i,P_j} + \mathrm{Pro}^{i,P_j} \qquad \text{(S1)}$$

In (S1), the first step is the finding of the promoter by a free floating RNAP (R) (i.e., not elongating from an upstream promoter) and the subsequent formation of the closed complex at the rate $K^{P_j}{}_{cc}$ (*4, 65, 85, 98*). Since this step is reversible, $K^{P_j}{}_{cc}$ accounts for the failure of several OCs, until one succeeds to proceed to the open complex formation. The second step, at the rate of $K^{P_j}_{oc}$ is the formation of the open complex, which is irreversible (*4*, *65*, *83*, *99*). Finally, the RNAP can escape and release the promoter (*67*) and then form an elongating complex along gene i ($EC^{i,j}$) (*67*, *100*). Using equation (S1) one can implement a model of production of EC's from each promoter in each operon of *E. coli*. Finally, tuning $K^{P_j}_{cc}$ and $K^{P_j}_{cc}$ allows changing these events' kinetics between stress and control conditions.

Next, we consider what can occur to an *EC* as it elongates along gene *i* in operon *k*. The frequency of those events will depend on the upstream and downstream structure of the operon, relative to the position of *EC*. It will also depend on whether there are upstream promoters. For simplicity, we start by assuming an operon with one initial promoter, but no internal promoters. In that case, the following 2 reactions suffice to model elongation.

First, reaction (S2) models $EC^{i,P_j}$ as it progresses from gene *i* to gene *i+1*. In that process, it completes the assembly of $RNA^{i,P_j}$. Next, Reaction (S3) models spontaneous premature terminations of the RNAP (R), while transcribing gene *i*. As noted, this event has many potential causes (e.g., transcriptional pauses or collisions between elongating RNAPs) but is modeled here as one process whose rate of occurrence can be tuned by $k_{fall}$. This rate constant allows changing these events' propensity between stress and control conditions.

$$EC^{i,P_j} \xrightarrow{K^{i}{}_{el(Dnuc)}} EC^{i+1,P_j} + RNA^{i,P_j} \qquad (S2)$$

$$EC^{i,P_j} \xrightarrow{K_{fall}} R \qquad (S3)$$

Finally, the model accounts explicitly for multiple promoters in the operons. For simplicity, here all such promoters are assumed to have the same response strength to a stress. If there are promoters upstream and/or downstream from the promoter of interest, RNAPs will interact and potentially cause premature terminations (other than the spontaneous ones modelled above). E.g., upstream promoters can "send" elongating RNAP's which will cause premature terminations in the OC formation of our promoter of interest or of the elongating RNAP. Similarly, CCs and OCs in downstream promoters can be hampered by an EC (Reactions S4 and S5, respectively) or can hamper the elongating RNAPs (reactions S6 and S7, respectively) from our promoter of interest. We assume that either the RNAP or the EC can be terminated during a collision (with the same probability, for simplicity) but that, in general, the termination of both, in the same event, is sufficiently unlikely to not require modelling. Overall, to account for terminations due to initiation events in promoters, we model the following events:

$$EC^{i+1,P_j} + CC^{i+1,P_{j+1}} \xrightarrow{K^{CC}{}_{col}} R + CC^{i+1,P_{j+1}} \qquad (S4)$$

$$EC^{i+1,P_j} + OC^{i+1,P_{j+1}} \xrightarrow{K^{OC}{}_{col}} R + OC^{i+1,P_{j+1}} \qquad (S5)$$

$$EC^{i+1,P_j} + CC^{i+1,P_{j+1}} \xrightarrow{K^{CC}{}_{col}} R + EC^{i+1,P_j} \qquad (S6)$$

$$EC^{i+1,P_j} + OC^{i+1,P_{j+1}} \xrightarrow{K^{OC}{}_{col}} R + EC^{i+1,P_j} \qquad (S7)$$

Noteworthy, in addition, we did not consider TTS in our model, given only 53 internal TTSs in RegulonDB.

To test our model, we simulated it using SGNSim (*53*) a stochastic system simulator based on Gillespie algorithm. Rate constants used are shown in Supplementary Table S16. From the resulting data, we calculated $\mu_{|\Delta LFC|}$ from all simulations and plotted it as a function of $L_G$ (Supplementary Figure S31). In agreement with the hypothesis, the model with internal promoters follows a sinusoidal curve and the model with no internal promoters follows a linear regression.

### Supplementary Figures

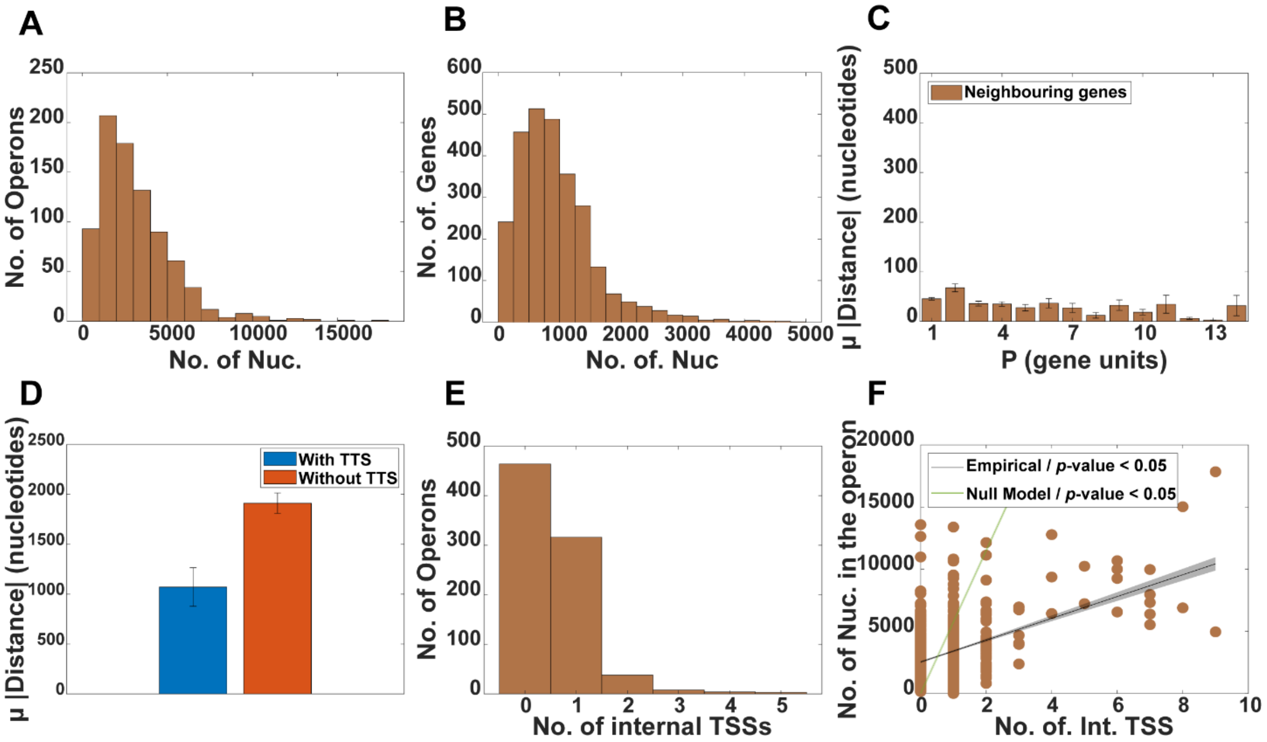


**Figure S1. Structural properties of the operons of *E. coli*. (A)** Histogram of the number of nucleotides ("Nuc.") composing operons. **(B)** Distribution of the number of genes in operons with a given nucleotide length. **(C)** Mean number of nucleotides between nearest neighbor genes in the same operon as function of their position (*P*) relative to the primary promoter. **(D)** Mean number of nucleotides between the end of the operon and the next downstream gene (red bar for cases where there is a TTS in between them and orange bar otherwise). **(E)** Distribution of the number of operons with a given number of TSSs. **(F)** Scattered plot between the number of nucleotides and the number of internal TSSs in the operons. We obtained a best fitting line and its *p*-value under the null hypothesis that the line is horizontal, using 'fitlm' in MATLAB. For t-statistic *p*-values > 0.05, we cannot reject the null hypothesis that the slope is 0. Also shown is a green line. This line is the best fit for the expected number of internal TSSs in operons of the same length, when randomly placed (assuming a model with the same number and length of operons as well as the same number of internal TSSs as E. coli). Data from 1000 independent runs.

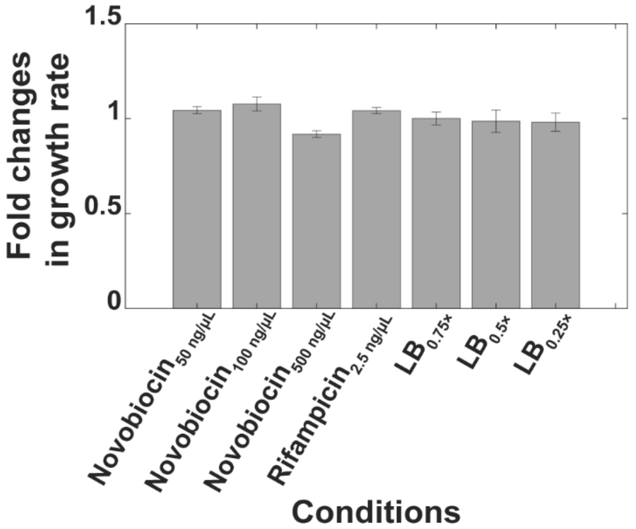


**Figure S2. Average fold changes in growth rates of cells under stress conditions**. Data shown is relative to the control condition (Methods section 'Bacterial strains and genome-wide stresses'). Error bars are the standard error of the mean (SEM) of the fold change of 3 biological replicates. Based on the error bars, the only condition with substantial differences in growth rate from the control condition is 'Novobiocin 500 ng/μL'.

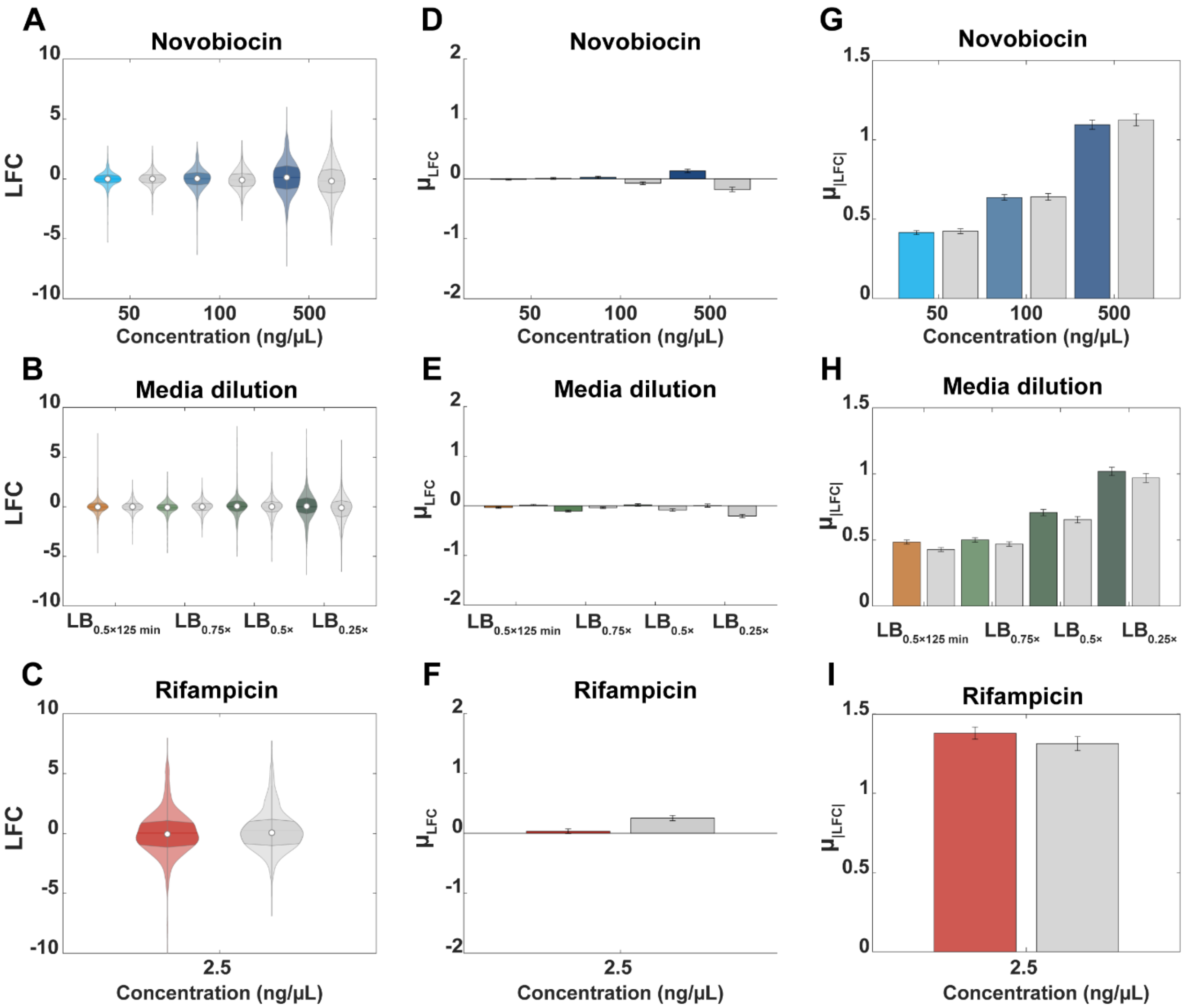


**Figure S3. Genome-wide single-gene stress responses**. (**A-C**) Violin plots (*101*) with the maximum, minimum, median, interquartile ranges, and probability density of the distributions of single gene LFCs, following the stresses. (**D-F**) Mean LFCs ($\mu_{LFC}$) of the distributions in A-C. (**G-I**). Mean absolute LFCs ($\mu_{|LFC|}$) of the distributions in A-C. In all graphs, the error bars are the standard error of the mean (SEM). In some figures, the error bars are not visible. The colored data are from genes in operons, and the grey data are from genes not in operons. The media used for Figures B, E, and H are labeled according to their composition, described in Methods section 'RNA-seq'.

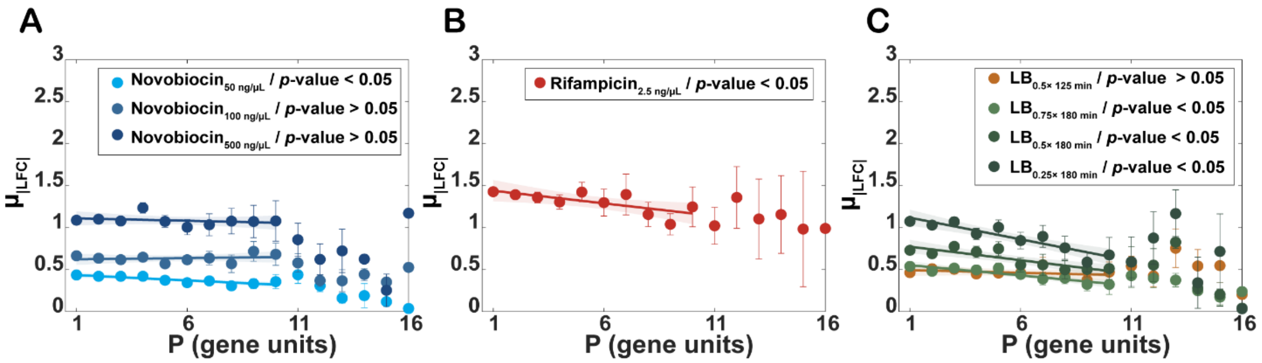


**Figure S4**. **Average absolute response strengths of genes, $\mu_{|LFC|}$, as a function of their position, *P*, in the operon**. Error bars are the standard error of the mean. The *p*-values are from statistical tests of whether the best linear fit differs from a horizontal line, using 'fitlm' in MATLAB. The shadow areas are the 95% confidence intervals. For t-statistic *p*-values > 0.05, we cannot reject the null hypothesis that the slope is 0. **(A**, **B**, and **C)** correspond to data from a specific condition, listed on the legend of the graph. In all cases, for $P > 10$, the number of data points is smaller than 30, which did not suffice to establish a trend.

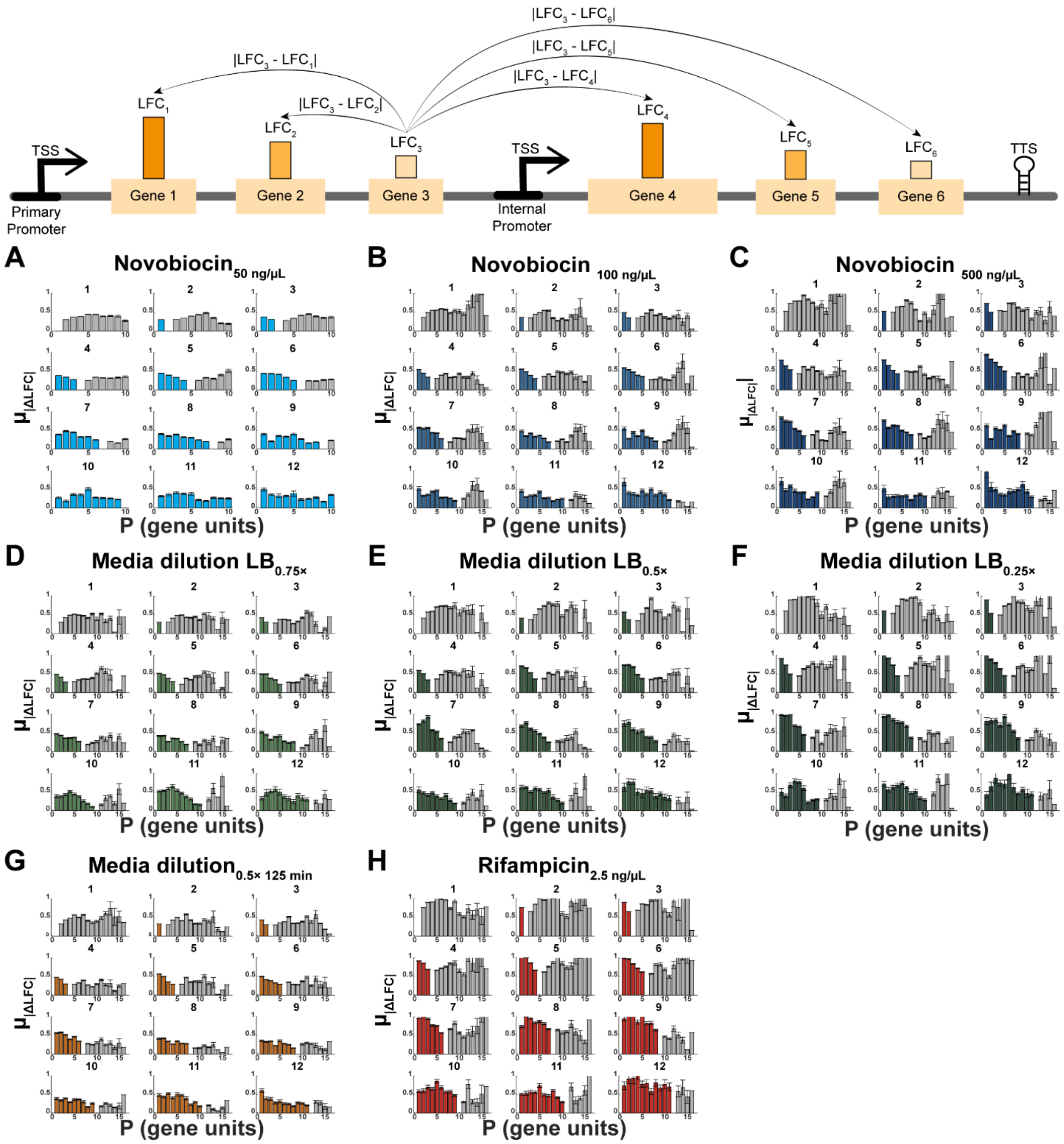


**Figure S5**. **Mean absolute differences between the LFCs, $\mu_{|\Delta LFC|}$, of pairs of genes for each position in the operons**. Top: illustration of how the data in Figures A-H are calculated. For each position ($P = 1, 2, \ldots, 15$) in each operon, we obtained the difference between the response strength of the gene in that position and the response strength of genes in other positions of the same operon. We then average those differences, for each position, over all operons. Colored and grey bars correspond to genes prior and after the position of interest (from whom no bar is shown), respectively. Vertical error bars are the standard errors of the mean (SEM). **(A-H)** correspond to data from a specific condition, listed on top of the graph, specifically. The position of interest is written above each subplot (as 1-12).

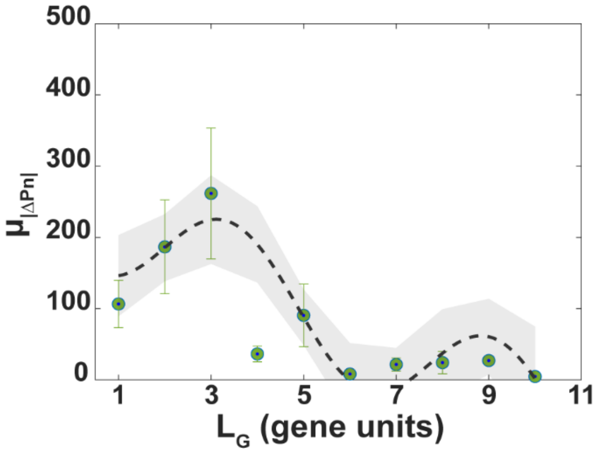


**Figure S6. Average absolute difference between the protein numbers (*$\mu|\Delta_{Pn}|$*) of genes in the same operon reported in** (*32*). Data shown as a function of the distance between the genes, $L_G$, in gene units. Vertical error bars are the standard errors of the mean (SEM). $L_G$ is set to a maximum value of 10, since no pair of genes considered is separated by a larger distance. The parameters of the best fitting curve are shown in Supplementary Table S2.

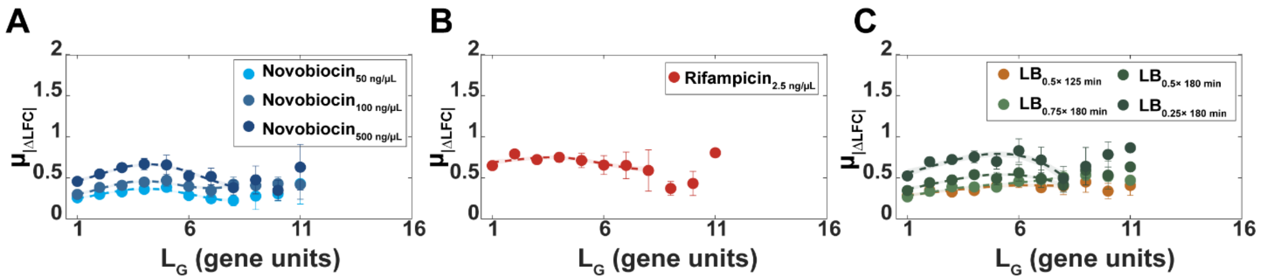


**Figure S7.** $\boldsymbol{\mu_{|\Delta LFC|}(L_G)}$ **of pairs of genes in the same operon**. Data from the 427 operons that are not subject to TF regulation. **(A)** Novobiocin stresses **(B)** Rifampicin stress **(C)** Shifts in RNAP concentrations due to shifts in media richness. Vertical error bars are the standard errors of the mean (SEM). The parameters of the best fitting curves and $R^2$ values are shown in Supplementary Table S3.

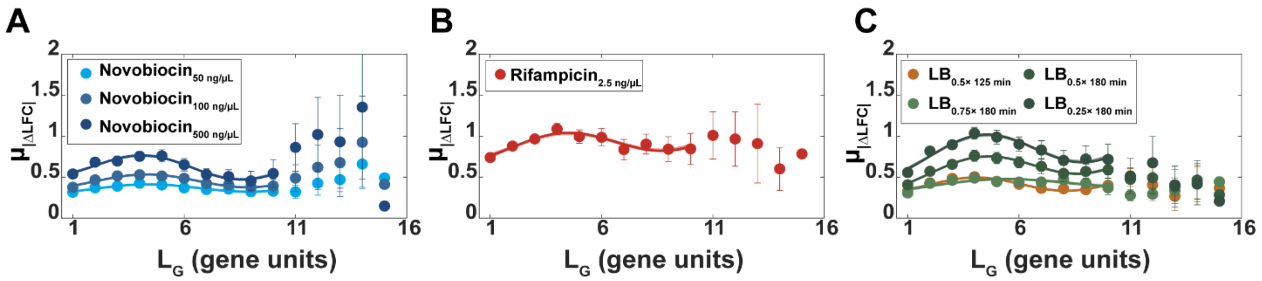


**Figure S8. $\mu_{|\Delta LFC|}(L_G)$ of pairs of genes in the same operon**. Data from the 406 operons that are subject to direct TF regulation. **(A)** Novobiocin stresses **(B)** Rifampicin stress **(C)** Shifts in RNAP concentrations due to shifts in media richness. Vertical error bars are the standard errors of the mean (SEM). The parameters of the best fitting curves and $R^2$ values are shown in Supplementary Table S4.

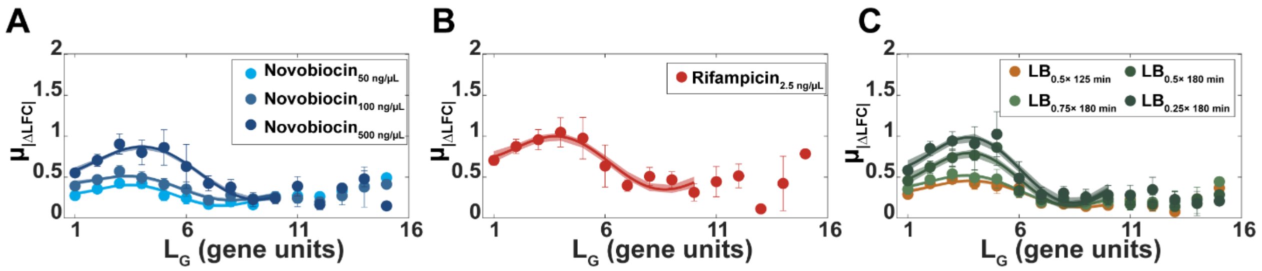


**Figure S9.** $\boldsymbol{\mu_{|\Delta LFC|}(L_G)}$ **of pairs of genes in the same operon**. Data from the 56 operons that are subject to self-TF regulation. **(A)** Novobiocin stresses **(B)** Rifampicin stress **(C)** Shifts in RNAP concentrations due to shifts in media richness. Vertical error bars are the standard errors of the mean (SEM). The parameters of the best fitting curves and $R^2$ values are shown in Supplementary Table S5.

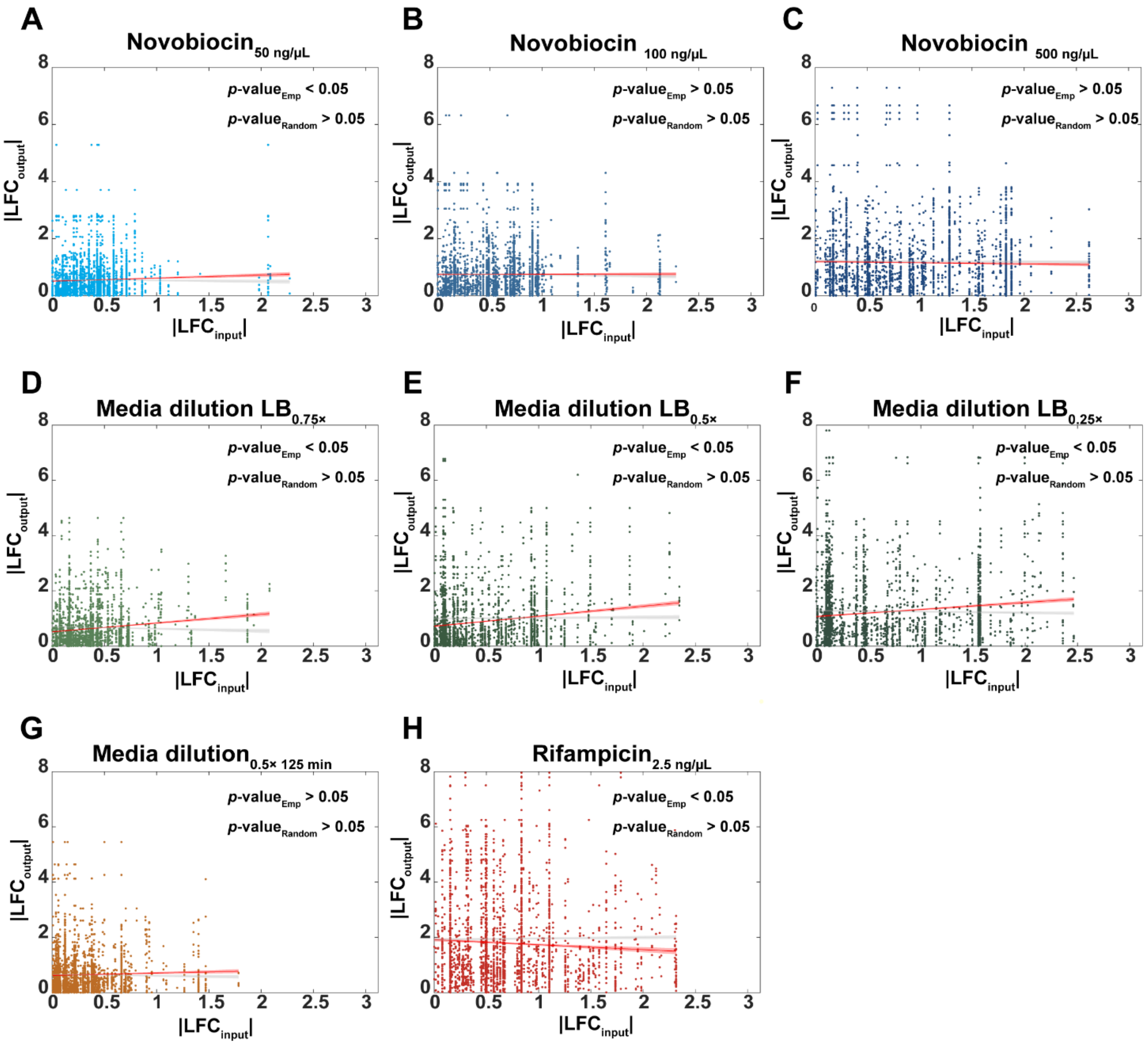


**Figure S10. Scatter plots between the |LFC|'s of pairs of genes, one gene coding for a TF input and the other gene is the output of that TF**. Each dot is one pair of genes. Also shown is the best fitting line of those dots (dark red) and its 95% confidence bounds (red shadow), along with the *p*-value under the null hypothesis that the line is horizontal (*p*-value$_{\text{Emp}}$). In addition, we randomized which genes are input and output of each TF and repeated the calculations 1000 times. From this, we obtained an average best fitting line and its p-value under the null hypothesis that the line is horizontal (*p*-values$_{\text{random}}$), using 'fitlm' in MATLAB. For t-statistic *p*-values > 0.05, we cannot reject the null hypothesis that the slope is 0. The shadow areas are the 95% confidence intervals. (**A-H**) correspond to data from a specific condition, listed on top of the graph, specifically.

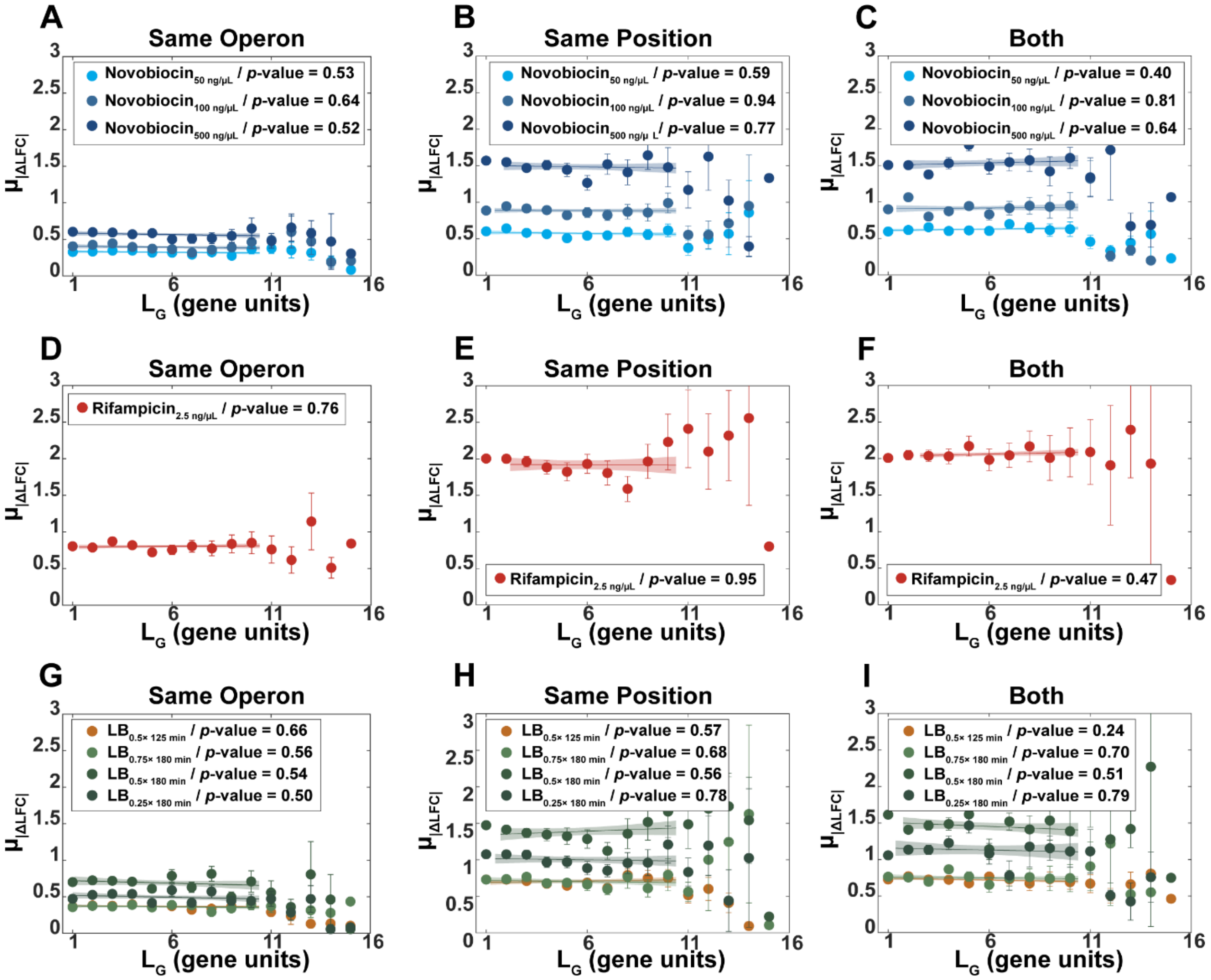


**Figure S11. Null models of $\mu_{|\Delta LFC|}(L_G)$ (A, D, G)**. We randomized the position of each gene in the same operon. **(B, E, H)**. We randomized the operon of each gene, but not their position. (**C, F, I**) We randomized both the operon of each gene as well as their position. Each data point is the average result from 1000 independent iterations. Vertical error bars are the standard errors of the mean (SEM). The *p*-value of each line is from a statistical test of whether the linear fit differs from a horizontal line, using 'fitlm' in MATLAB. For t-statistic *p*-values > 0.05, we cannot reject the null hypothesis that the slope is 0. The shadow areas are the 95% confidence intervals.

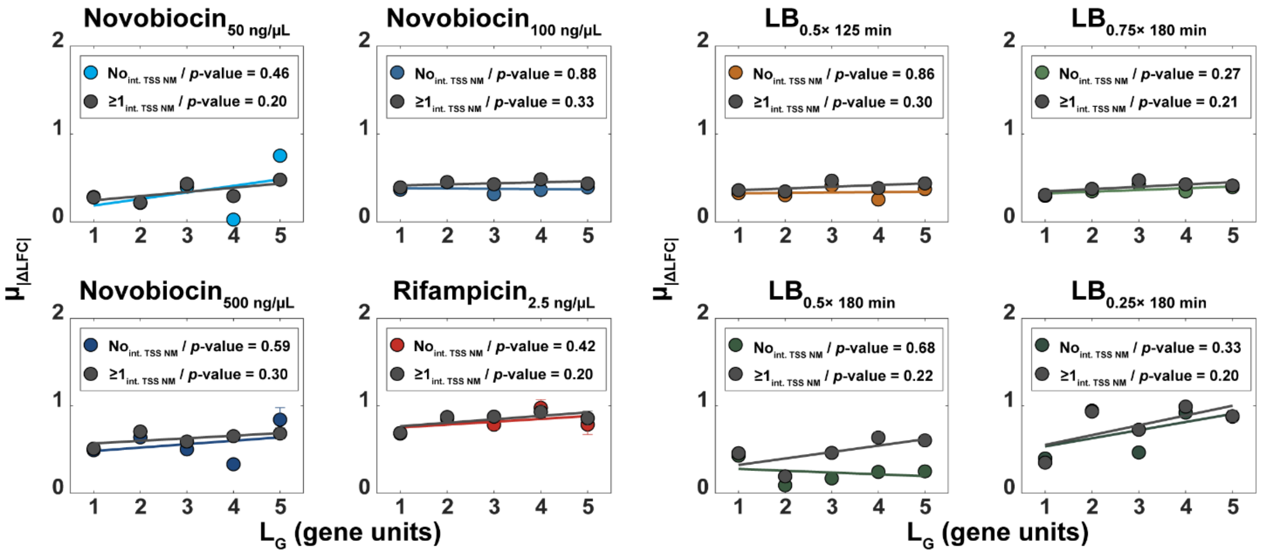


**Figure S12. Null models of $\mu_{|\Delta LFC|}(L_G)$ for with and without internal promoters**. Average absolute difference between the LFCs of pairs of genes in same operon ($\mu_{|\Delta LFC|}$) with and without internal TSS's in between (colored and gray lines, respectively). Data plotted only for $L_G < 6$ since there are very few pairs of genes without TSSs in between for $L_G > 5$. Vertical error bars are the standard errors of the mean (SEM). Also shown are the best linear fits and corresponding *p*-values. For *p*-value < 0.05, the t test statistics reject the null hypothesis that the best fitting line is not different from a horizontal line. The shadow areas are the 95% confidence intervals.

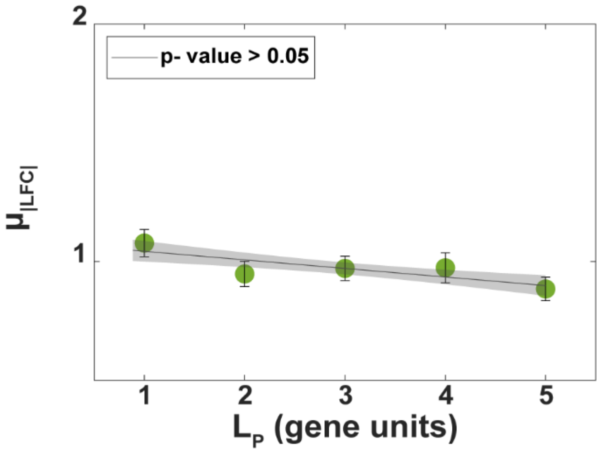


**Figure S13. Null models of $\mu_{|LFC|}$ ($L_p$).** Average absolute response strength, $\mu_{|LFC|}$, of the first gene immediately downstream of a TSS plotted as a function of $L_P$ (number of genes controlled by that TSS prior to any other downstream TSS, illustrated in Figure 1C) for the random model operons created by shuffling the position of the internal promoter. Vertical error bars are the standard errors of the mean (SEM). Also shown are the best linear fits and corresponding *p*-values. For *p*-value < 0.05, the t test statistics rejects the null hypothesis that the best fitting line is not different from a horizontal line. The shadow areas are the 95% confidence intervals.

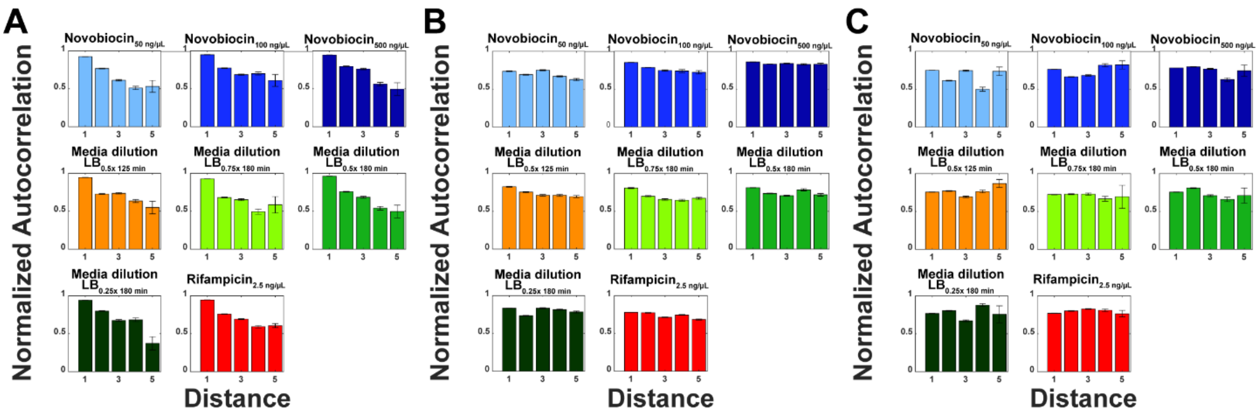


**Figure S14**: **Normalized unbiased autocorrelations of the response strengths of genes in the same operon, for each stress condition, as a function of the gene's distance from its closest upstream promoter**. **(A)** Genes and their *nearest* upstream promoter. **(B)** Genes and *any* upstream promoter, provided that there is another internal promoter in between them. **(C)** Genes and their nearest upstream promoter as in (A), however the distances from each gene to its promoter were randomly shuffled. Distances larger than 6 were not considered, since there are not sufficient data points for (A). The error bars are the standard error of the mean.

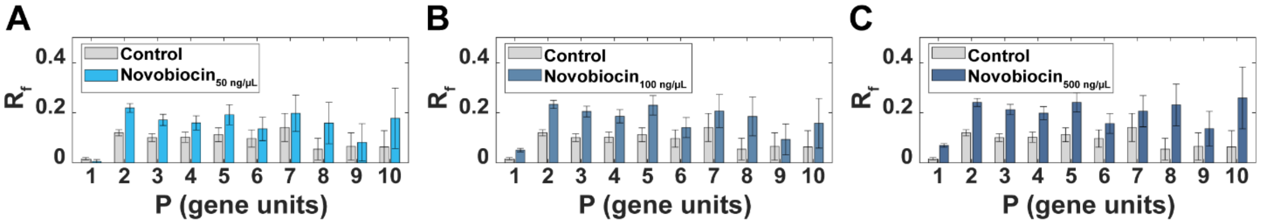


**Figure S15: Rates of RNAP premature terminations, $R_f$ for different concentrations of Novobiocin.** Average differences between the number of RNA reads from the starting region of a gene and the number of RNA reads from the ending region of the same gene, normalized by the number of reads of the starting region. Data is plotted as a function of the position (*P*) of the genes, relative to the primary TSS. The error bars are the standard error of the sum. (**A, B, C**) correspond to different novobiocin concentrations, listed on the legends of the graphs, specifically.

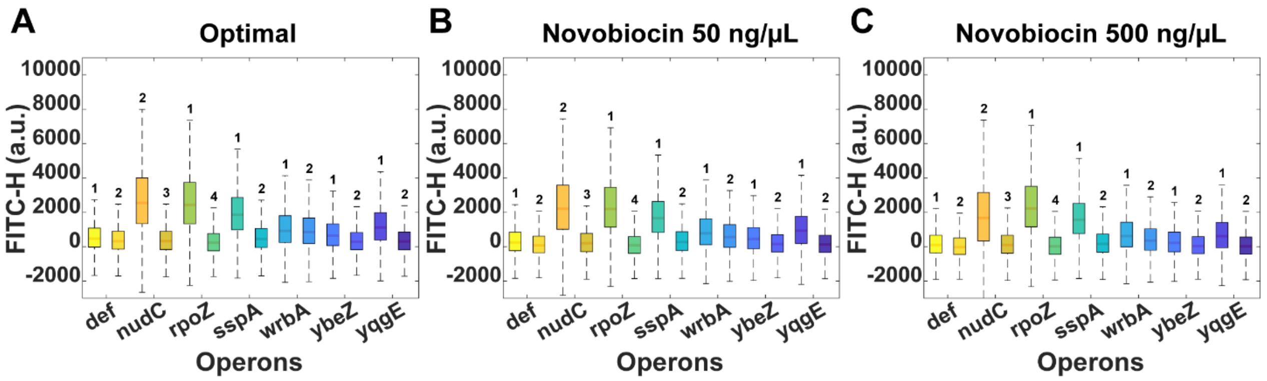


**Figure S16**. **Box plots of single-cell gene expression levels of two genes of the same operon**. Data measured by flow-cytometry, FITC-H. Each box is from one gene, while each pair of bars is from the same operon (named in the x-axis). The number on top of each bar corresponds to the position of the gene in the operon. In each box, the center indicates the median, and the bottom and top edges of the vertical line indicate the 25th and 75th percentiles, respectively. (**A, B, and C**) correspond to data from a specific condition, listed on top of the graph, specifically.

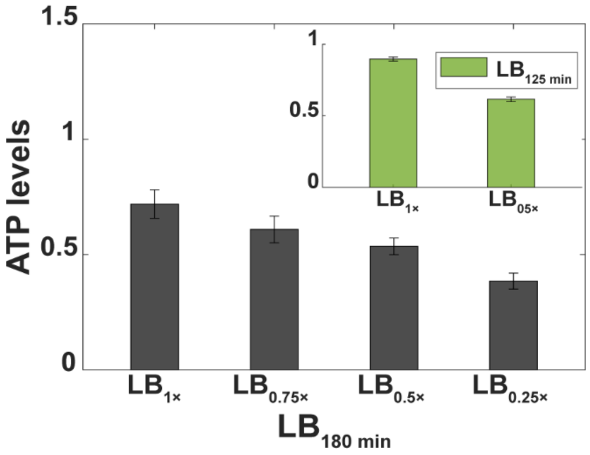


**Figure S17. Cellular ATP levels measured using the QUEEN-2m sensor**. Measurements conducted 180 minutes after placing the cells in tailored media (Methods section ‘RNA-seq’). We performed 3 replicates per condition. The inset is the same measurement at 125 minutes. The error bars are the standard error of the mean (SEM) calculated using error propagation method as described in methods section ‘Statistical tests’. Shown are the ratios between the 513 nm emission intensities at the two excitation wavelengths, used as a proxy for cellular ATP concentrations (Methods section ‘cellular ATP levels’).

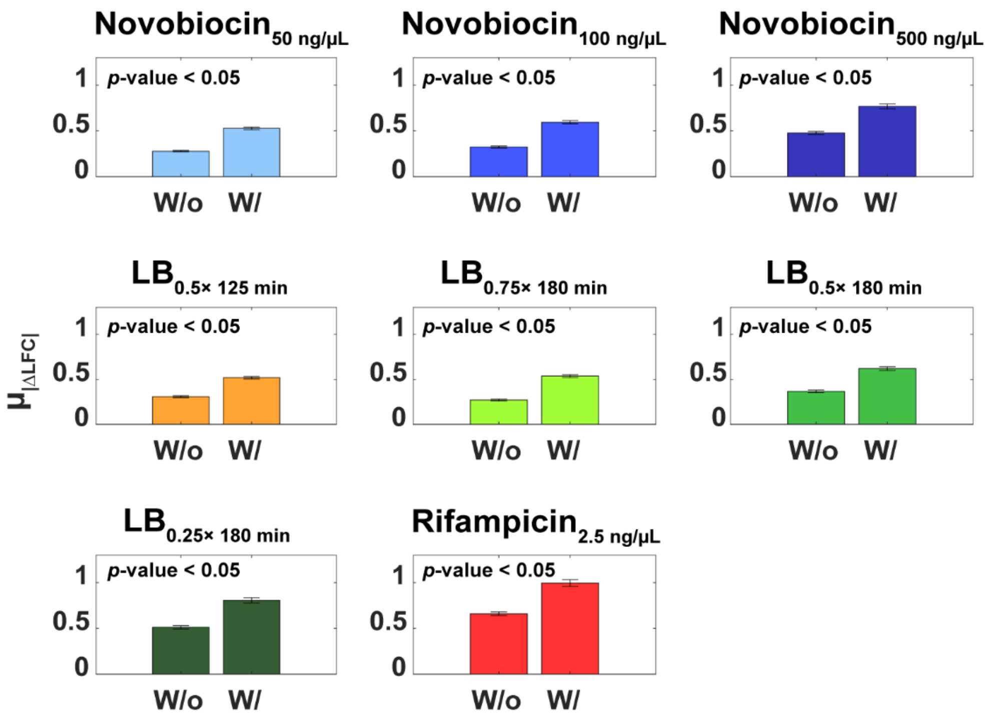


**Figure S18. $\mu_{|\Delta LFC|}$ between pairs of adjacent genes with and without a *pTTS* in between.** Average absolute difference in the response strength between adjacent gene pairs ($L_G = 1$) with ('W/') and without ('W/o') premature transcription termination sites (***pTTS***). Vertical error bars are the standard errors of the mean (SEM). The *p*-values are from t test statistics assessing whether two distributions are from independent random samples from normal distributions with equal means and equal, but unknown, variances. For *p*-value < 0.05, the t test statistics rejects the null hypothesis that the two distributions have equal means, at the 5% significance level.

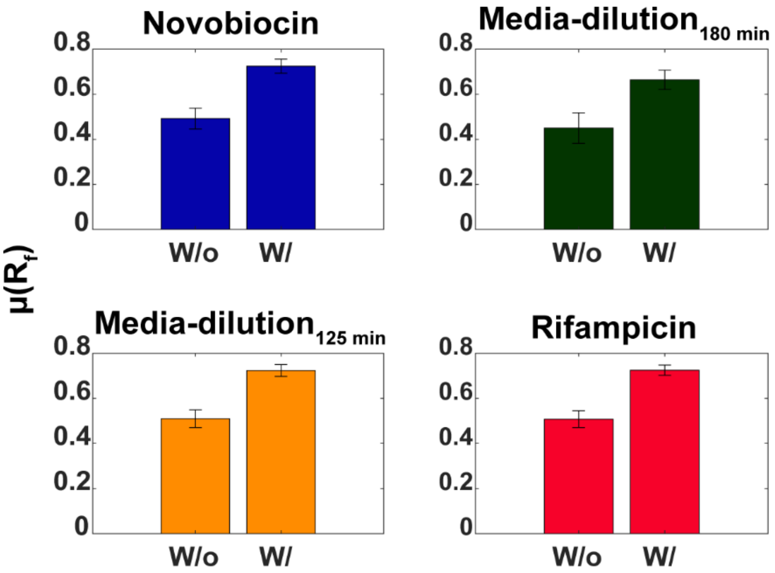


**Figure S19.** ***μ(R$_f$)* of genes with and without a *pTTS* in their coding region.** Average differences between the number of RNA reads from the starting region of a gene and the number of RNA reads from the ending region of the same gene, normalized by the number of reads from the starting region. Data is plotted with (W/) or without (W/o) *pTTS* sites in between them, while the vertical error bars are the standard errors of the mean (SEM). The data for "Novobiocin" and for "Media-dilution at 180 minutes" are, in both cases, averages from the three different stresses, respectively (Methods section 'RNA-seq').

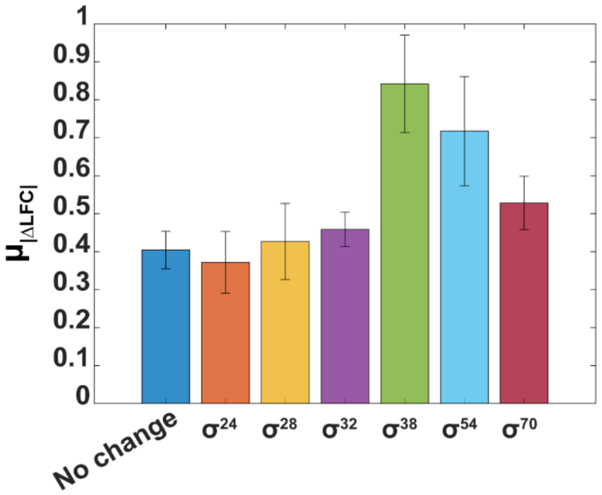


**Figure S20. $\mu_{|\Delta LFC|}$ between adjacent genes in operons separated by a promoter whose σ factor preference differs from the preference of the upstream promoter.** Data shown as a function of the σ factor preference of the upstream promoter and obtained from all stress conditions. Vertical error bars are the standard errors of the mean (SEM). The dark blue line is the control condition, when there is no change between the σ factor preference of the upstream and downstream promoters.

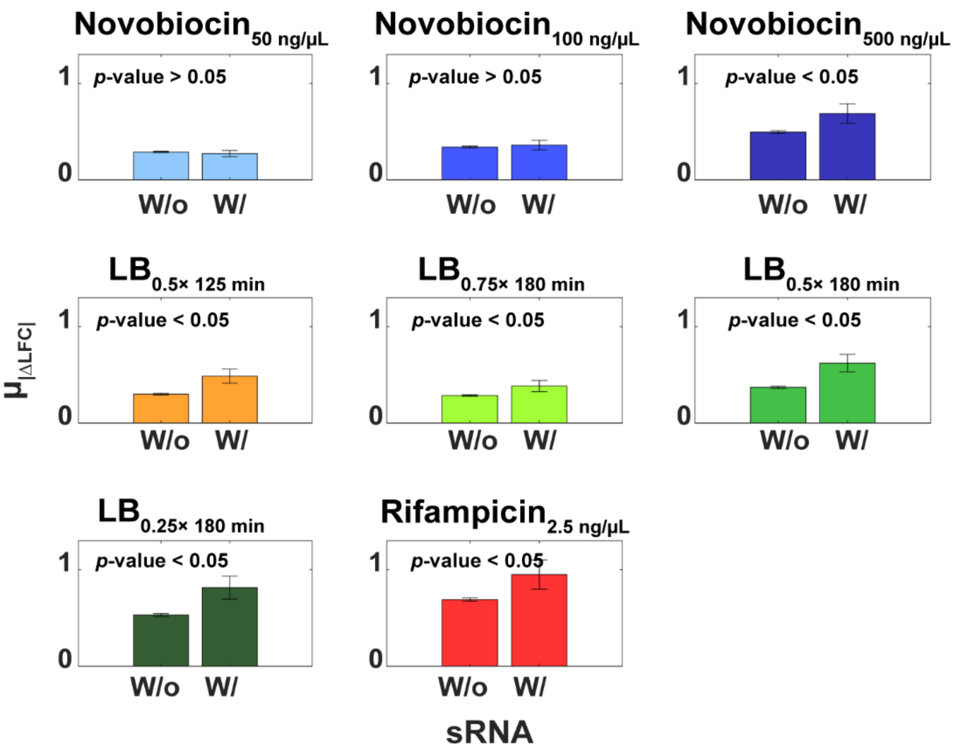


**Figure S21. $\mu_{|\Delta LFC|}$ of adjacent genes with at least one the genes having an sRNA binding site region.** Average absolute difference in the response strength between adjacent gene pairs of $L_G = 1$ with (W/) and without (W/o) sRNA binding sites in between them. Vertical error bars are the standard errors of the mean (SEM). The $p$-values are from t test statistics assessing whether two distributions are from independent random samples from normal distributions with equal means and equal, but unknown, variances. For $p$-value < 0.05, the t test statistics rejects the null hypothesis that the two distributions have equal means, at the 5% significance level.

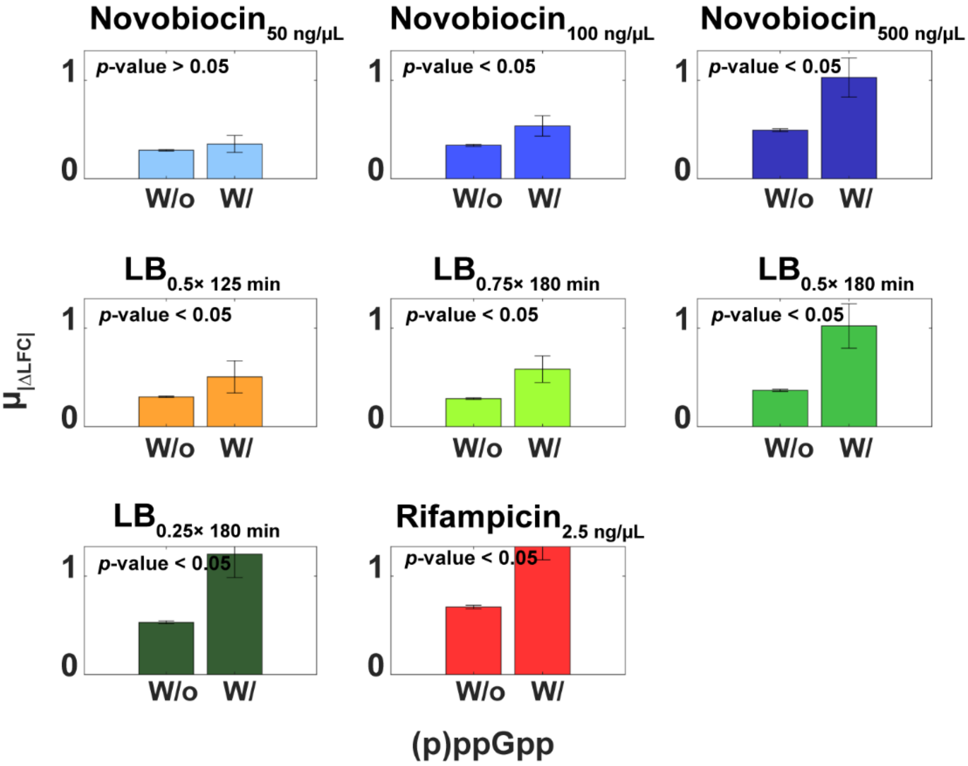


**Figure S22. $\mu_{|\Delta LFC|}$ of genes with and without (p)ppGpp sensitivity.** Average absolute difference in the response strength between adjacent gene pairs of $L_G = 1$ with (W/) and without (W/o) (p)ppGpp sensitivity between them. Vertical error bars are the standard errors of the mean (SEM). The *p*-values are from t test statistics assessing whether two distributions are from independent random samples from normal distributions with equal means and equal, but unknown, variances. For *p*-value < 0.05, the t test statistics rejects the null hypothesis that the two distributions have equal means, at the 5% significance level.

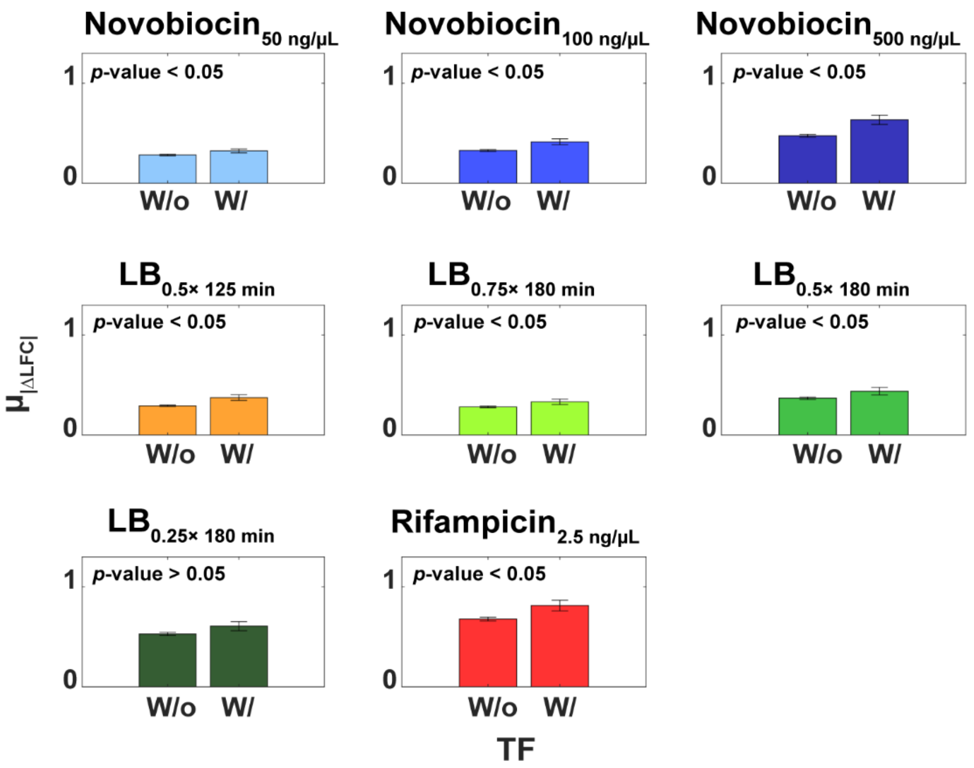


**Figure S23.** $\mu_{|\Delta LFC|}$ **of genes with and without transcription factor (TF) binding sites in between them.** Average absolute difference in the response strength between adjacent gene pairs of $L_G = 1$ with (W/) and without (W/o) **TF** binding sites in between them. Vertical error bars are the standard errors of the mean (SEM). The *p*-values are from t test statistics assessing whether two distributions are from independent random samples from normal distributions with equal means and equal, but unknown, variances. For *p*-value < 0.05, the t test statistics rejects the null hypothesis that the two distributions have equal means, at the 5% significance level.

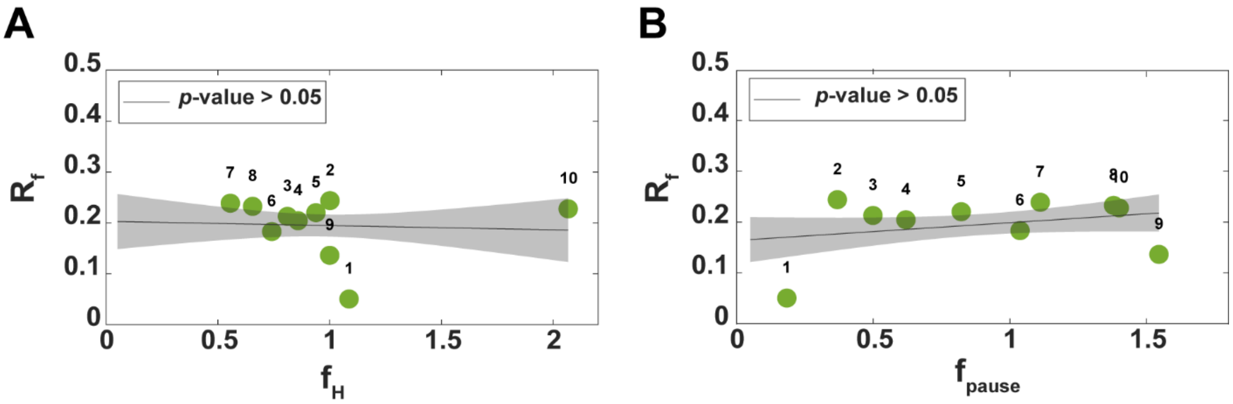


**Figure S24.** Average differences in numbers of RNA reads from the starting and the ending regions of each gene, normalized by the former, $\boldsymbol{R_f}$. **(A)** $R_f$ as a function of the frequency ($f_H$) of homopolymeric A/T tracts in the sequence of the gene. Again, frequency stands for the total number of tracts in a position, divided by the total number of operons with a gene in that position. The *p*-values of the lines are from statistical tests of whether the linear fits differ from horizontal lines, using 'fitlm' in MATLAB. For t-statistic $p$-values > 0.05, we cannot reject the null hypothesis that the slope is 0. The shadow areas are the 95% confidence intervals. In both figures, the numbers on top of the circles inform on the position in the operon of the genes corresponding to the circles. **(B)** $R_f$ plotted as a function of the frequency ($f_{pause}$) of pause sequences found in genes at given distances from the primary promoter (in gene units). This frequency is estimated from the total number of pause sequences found at genes, divided by the total number of operons with a gene in that position (Methods section ‘Estimating genome-wide differences in reads between the start and end of genes in operons’ in the main manuscript’). The pause sequences considered are reported in (*69*).

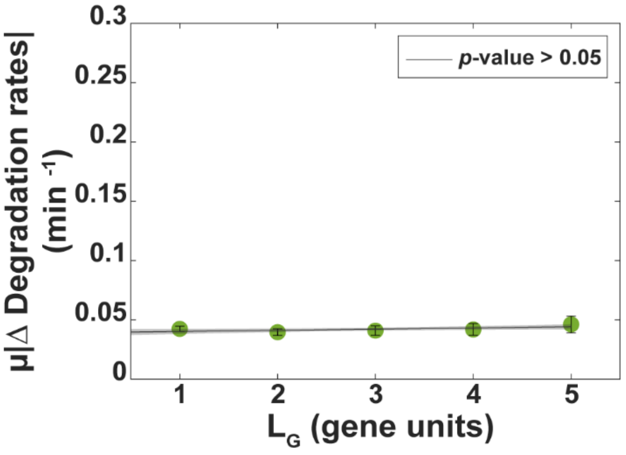


**Figure S25. Differences in RNA degradation rates as a function of distances between genes in operons without internal promoters in between.** Average difference between the degradation rate of RNAs in the same operon plotted as a function of the distance between the genes ($L_G$) coding for the RNAs. The *p*-values of the lines are from statistical tests of whether the linear fits differ from horizontal lines, using 'fitlm' in MATLAB. For t-statistic *p*-values > 0.05, we cannot reject the null hypothesis that the slope is 0. The shadow areas are the 95% confidence intervals.

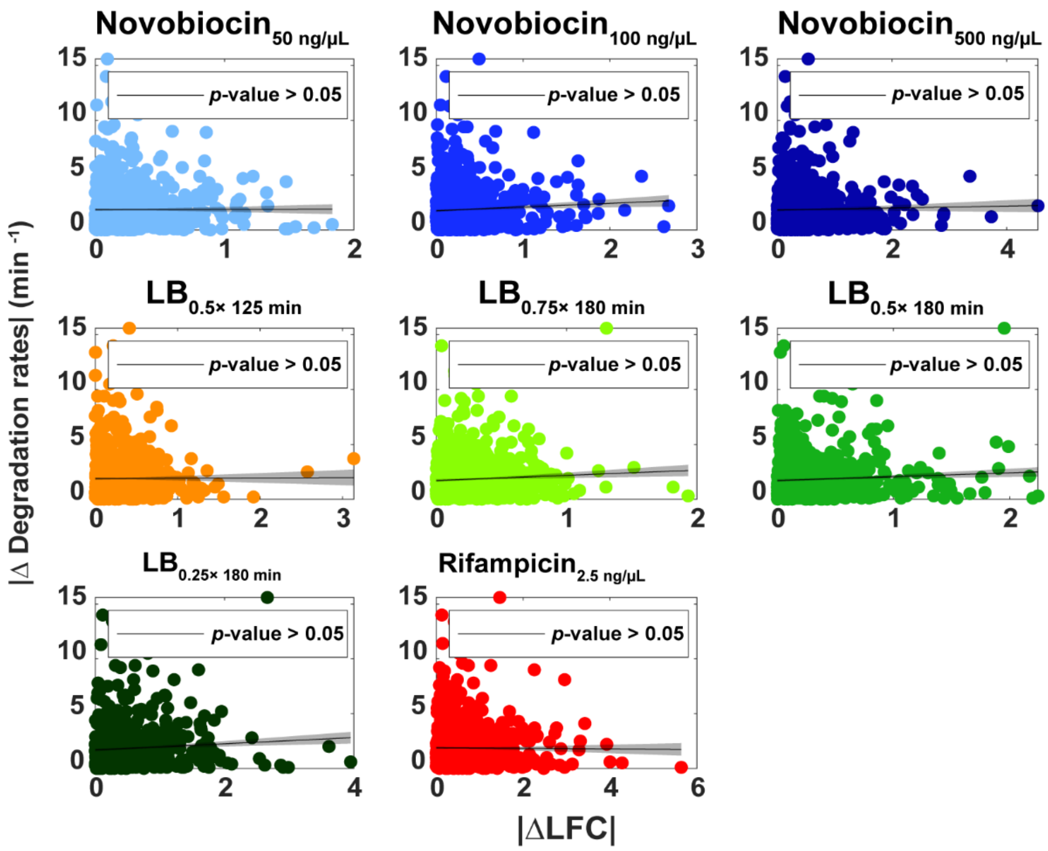


**Figure S26. |*ΔLFC*| between pairs of genes plotted against their absolute differences in degradation rates (|*Δ*|).** Average absolute difference in the response strength between gene pairs of $L_G = 1$ plotted against their differences in degradation rates (*80*). Also shown are the best linear fits and corresponding *p*-values. For *p*-value < 0.05, the t test statistics rejects the null hypothesis that the best fitting line is not different from a horizontal line. The shadow areas are the 95% confidence intervals. A few, rare, data points have differences in degradation rates above 15. These points are not shown for easier visualization of the results. Nevertheless, they were considered when we estimated the linear fits.

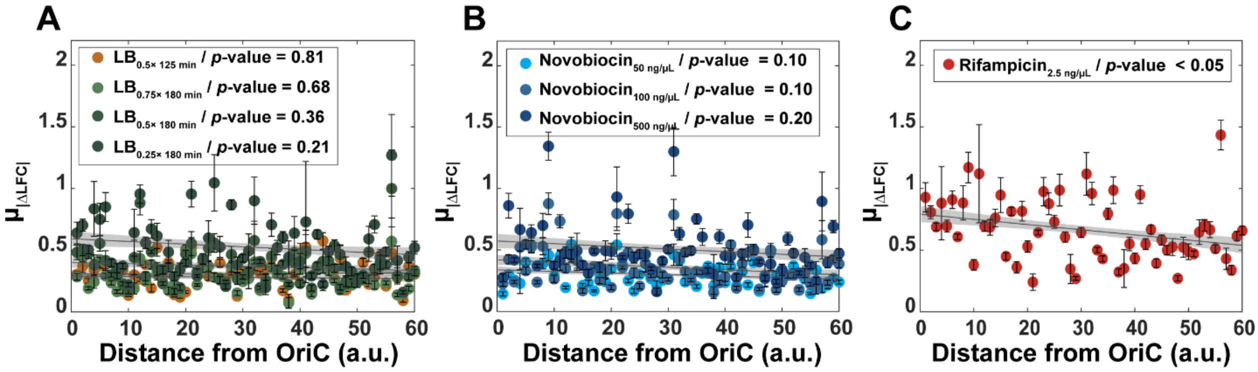


**Figure S27. Average operon response strengths to the genome wide stresses and distance from origin of DNA replication.** (A, B, C) $\mu_{|\Delta LFC|}$ stands for the |ΔLFC| of nearest neighbor genes in the same operons averaged over all pairs and all operons within specific DNA regions. In the x axis, each arbitrary unit (a.u.) from the origin of replication (OriC) corresponds to a cohort of 38680 base pairs. For *p*-value < 0.05, the *t* test statistics rejects the null hypothesis that the best fitting line is not different from a horizontal line. The shadow areas are the 95% confidence intervals.

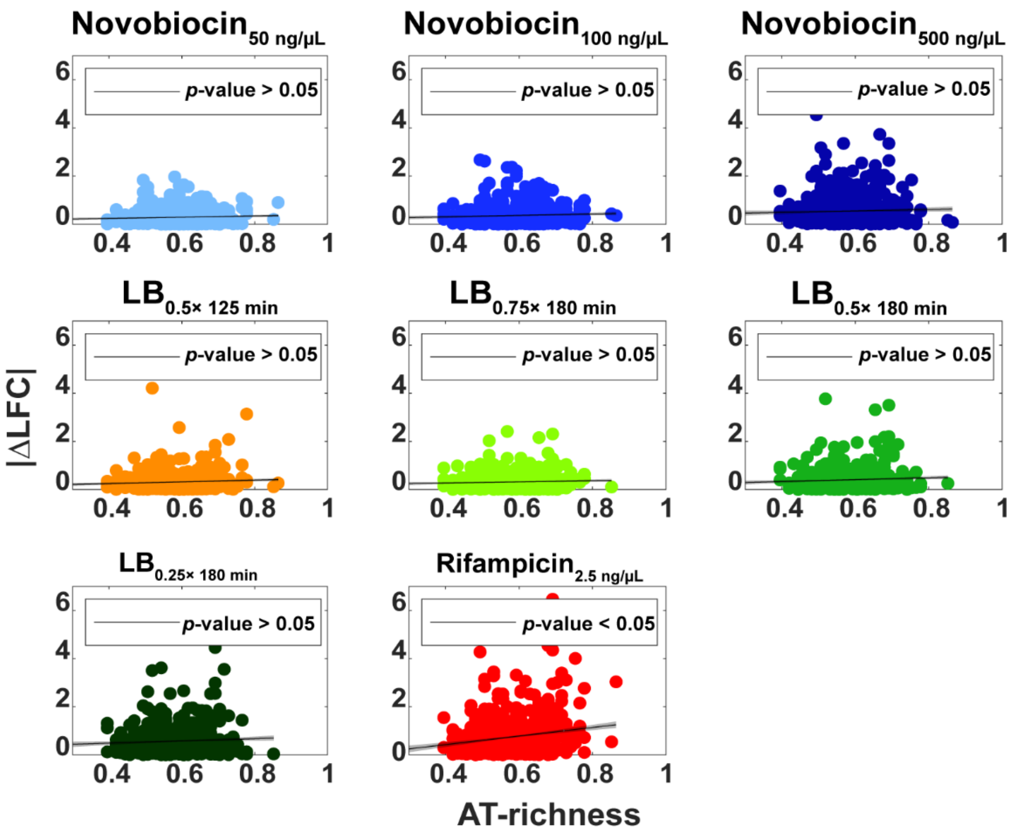


**Figure S28. |ΔLFC| between upstream and downstream genes from promoters plotted against that promoter's AT-richness.** Average absolute difference in the response strength between adjacent gene pairs of $L_G = 1$ plotted against the AT richness of the promoter. Also shown are best linear fits and corresponding $p$-values. For $p$-value $< 0.05$, the t test statistics rejects the null hypothesis that the best fitting line is not different from a horizontal line. The shadow areas are the 95% confidence intervals.

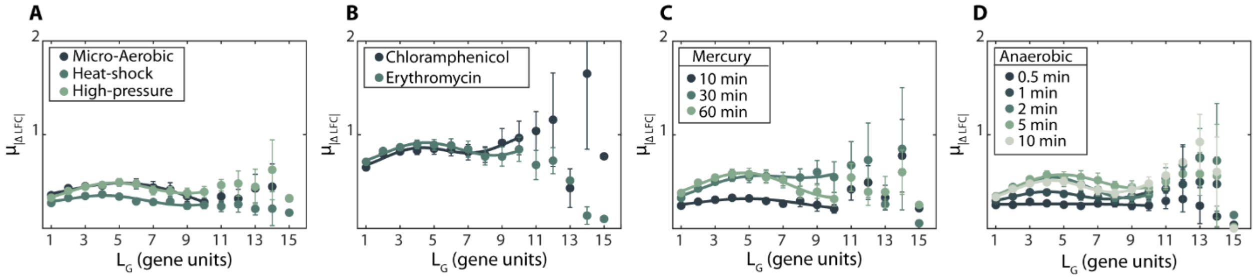


**Figure S29: Average absolute differences in the response strengths of genes in the same operon ($\mu_{|\Delta LFC|}$) as a function of the distance between genes in the same operon, $L_G$, for *E. coli* cells under other stresses**. From **(A)** to **(D),** each data is the $\mu_{|\Delta LFC|}$ of all pairs of genes of the same operon distanced by $L_G$ genes. The error bars are the standard error of the mean. Also shown are the best fits of functions of the form (*Eq 1*) along with their $R^2$ values. See Supplementary Table S11, for the best fitting parameter values. In all cases, $R^2$ is larger than 0.75 (average $R^2$ of 0.90). Finally, the data on heat-shock responses was obtained by AIR-seq. Data from (*37–42*). In **(C)** and **(D)**, the time values correspond to when measurements were conducted following the perturbation.

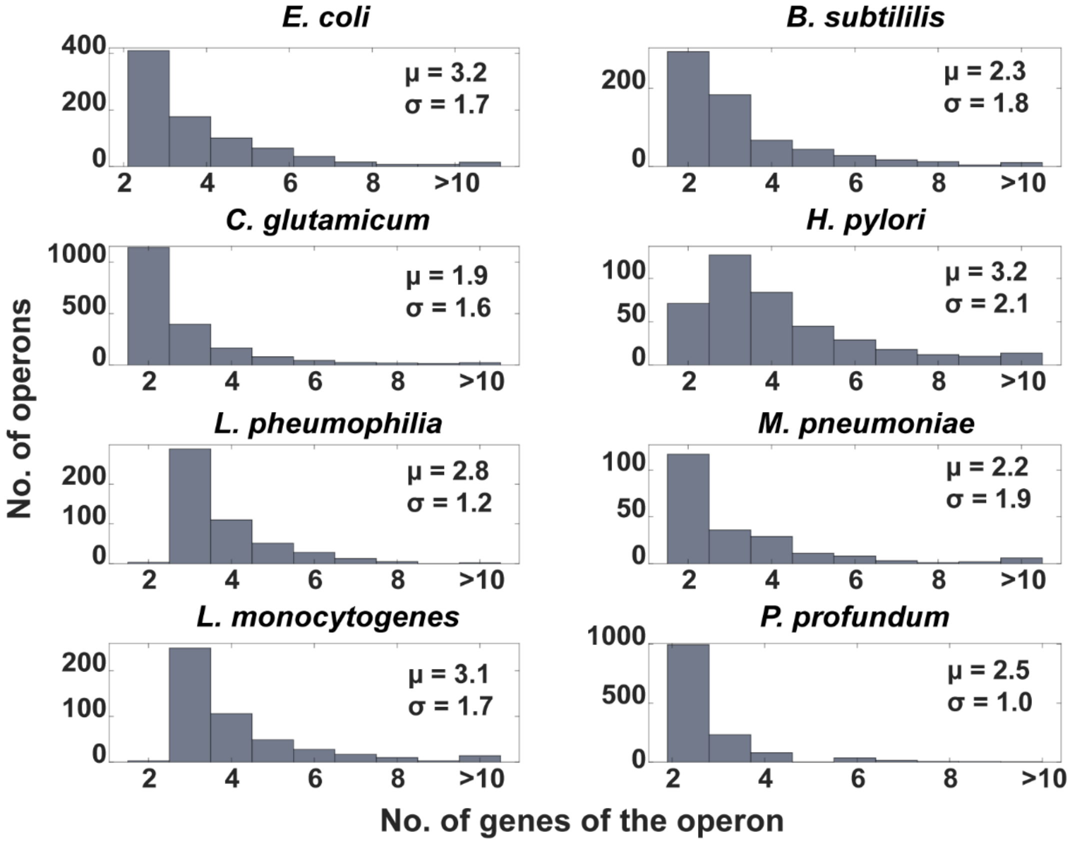


**Figure S30. Distribution of the number of operons as a function of their length (number of genes).** Data for 8 bacterial species. 'Size' stands for the number of genes of the operon while μ is the mean and σ is the standard deviation of the distribution. A 2-sample Kolmogorov-Smirnov (KS) test (Supplementary Table S12) found that, for 5 out of the 7 species, the distribution could not be distinguished from *E. coli*, shown in the top left.

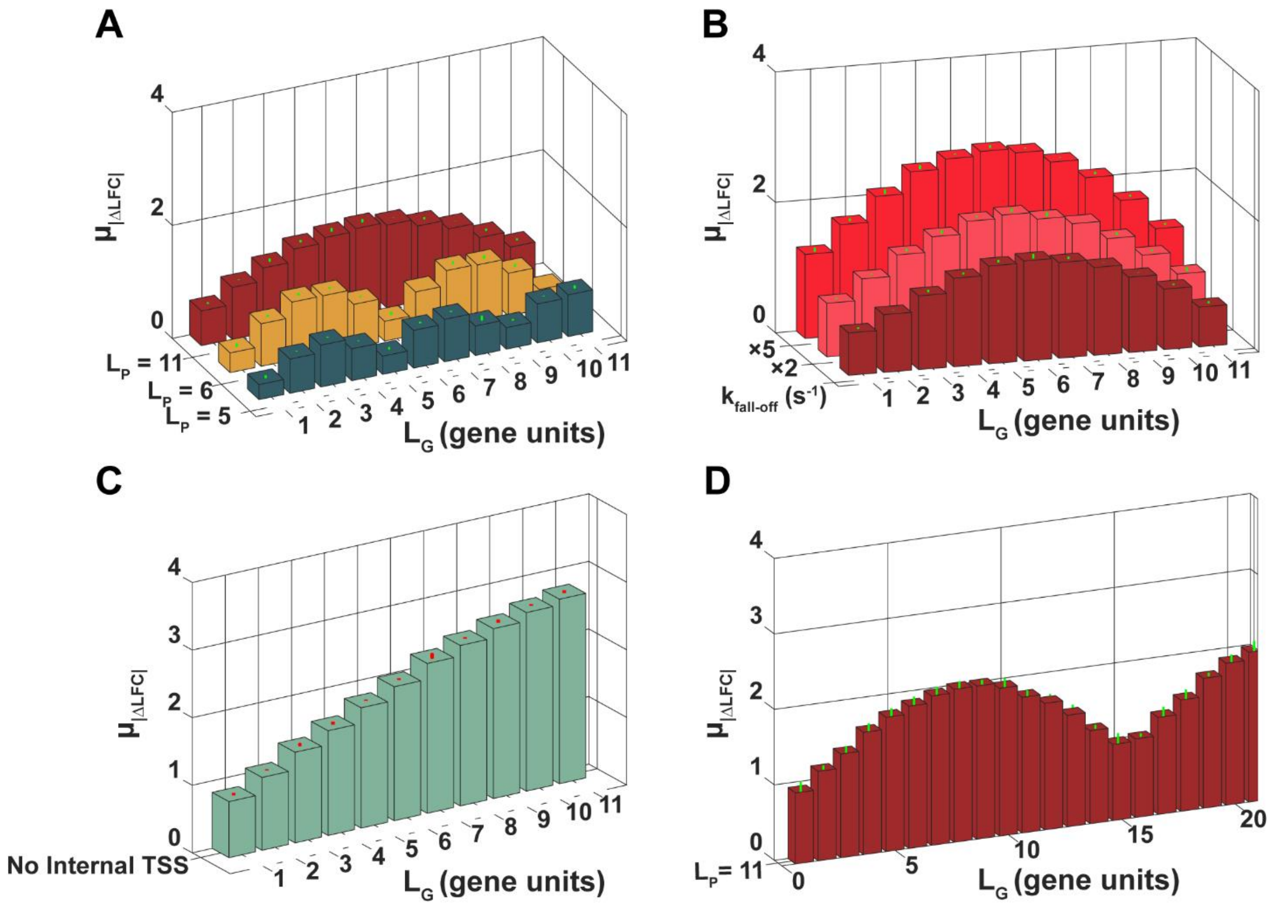


**Figure S31**. **$\mu_{|\Delta RNA|}$(*$L_G$*) of model operons (22 genes-long). (A)** Results for three models that differ in the number of genes between internal promoters of the operon ($L_P$). Since we kept the length of the operon (number of genes) constant, we altered $L_P$ by changing the number of internal promoters. Then, by fitting a sine-function to each curve (equation 1 in the main manuscript), we estimated the spatial frequency and amplitudes to be (0.13, 0.79), (0.1, 1.04), and (0.05, 2.64) for $L_P$ = 5, 6, and 11, respectively. **(B)** Results for three models differing in rates of spontaneous RNAP premature terminations, $k_{fall\text{-}off}$ (in all models, assuming $L_P$ = 11, for simplicity). We estimated the periods and amplitudes to be (0.05, 1.52), (0.05, 1.95), and (0.05, 73.7) for $k_{fall\text{-}off}$ rates equal to 0.0015, 2×0.0015, and 5×0.0015, respectively. **(C)** Result for a model without internal promoters. (**D**) Model operon with $L_P$ = 11 where the primary promoter does not change its dynamics with the stress, while the internal promoter responsiveness increases its transcription rate by 100%. Error bars are the standard error of the mean (not visible in most cases). See Supplementary Methods 'Stochastic Model' and Supplementary Table S16 for additional details on the models and simulations. Note that the y-axes differ in size. We used model operons with 22 genes to enhance the possible values of ***$L_G$***, which facilitates observing the sine curves.

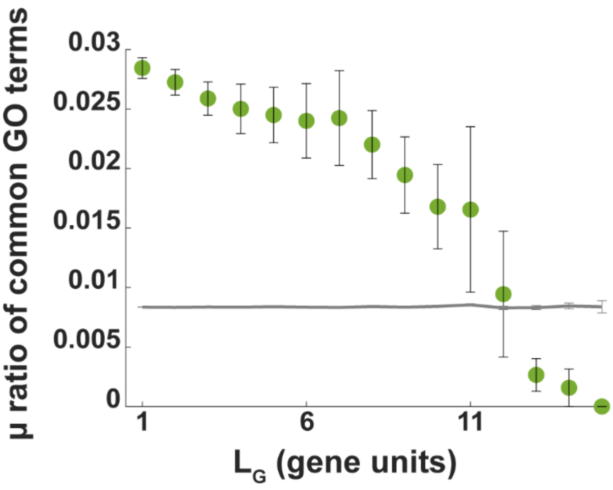


**Figure S32**. **Average ($\mu$) genome-wide ratio of common GO terms of pairs of genes in the same operon**. Data plotted as a function of the $L_G$ of the pair of genes. The error bars are the standard error of the mean. The gray, best-fitting line was obtained from estimations using the null model (Methods section 'Gene Ontology'). Note the increased uncertainty for $L_G > 10$.

**Supplementary Tables**

**Table S1. Parameters of the best fitting sine function**. We used a sine function of the form $g(L_G) = A_0 + A.L_G + B.\sin(2\pi.f.L_G)$. Also shown are the $R^2$ values to evaluate if the sinusoidal curve is well fitted, along with the adjusted (adj.) $R^2$ values, which consider the number of parameters used to fit the curve. Related to Figure 2A-C in the main manuscript.

| **Novobiocin** | | | |
|---|---|---|---|
| | **50 ng/µL** | **100 ng/µL** | **500 ng/µL** |
| **$A_0$** | 0.331 | 0.410 | 0.604 |
| **A** | 0.004 | 0.006 | -0.002 |
| **B** | 0.052 | 0.077 | 0.136 |
| **f** | 0.11 | 0.10 | 0.11 |
| **$R^2$** | 0.898 | 0.926 | 0.976 |
| **adj. $R^2$** | 0.847 | 0.889 | 0.964 |

| **Media dilution** | | | | |
|---|---|---|---|---|
| | **0.5× (125 min)** | **0.75× (180 min)** | **0.5× (180 min)** | **0.25× (180 min)** |
| **$A_0$** | 0.368 | 0.355 | 0.477 | 0.694 |
| **A** | 0.006 | 0.006 | 0.023 | 0.02 |
| **B** | 0.065 | 0.075 | 0.126 | 0.182 |
| **f** | 0.11 | 0.09 | 0.10 | 0.11 |
| **$R^2$** | 0.916 | 0.739 | 0.946 | 0.928 |
| **adj. $R^2$** | 0.874 | 0.657 | 0.919 | 0.892 |

| **Rifampicin (2.5 ng/µL)** | |
|---|---|
| **$A_0$** | 0.800 |
| **A** | 0.010 |
| **B** | 0.117 |
| **f** | 0.10 |
| **$R^2$** | 0.823 |
| **adj. $R^2$** | 0.734 |

**Table S2. Parameters of the best fitting sine function**. We used a sine function of the form $g(L_G) = A_0 + A.L_G + B.\sin(2\pi.f.L_G)$. Also shown are the $R^2$ values to evaluate if the sinusoidal curve is well fitted, along with the adjusted (adj.) $R^2$ values, which consider the number of parameters used to fit the curve. Related to Supplementary Figure S6.

| | |
|---|---|
| **$A_0$** | 179.70 |
| **A** | -23.75 |
| **B** | 49.11 |
| **f** | 0.17 |
| **$R^2$** | 0.668 |
| **adj. $R^2$** | 0.598 |

**Table S3. Parameters of the best fitting sine function**. We used a sine function of the form $g(L_G) = A_0 + A.L_G + B.\sin(2\pi.f.L_G)$. Also shown are the $R^2$ values to evaluate if the sinusoidal curve is well fitted, along with the adjusted (adj.) $R^2$ values, which consider the number of parameters used to fit the curve. Related to Supplementary Figure S7.

| **Novobiocin** | | | |
|---|---|---|---|
| | **50 ng/µL** | **100 ng/µL** | **500 ng/µL** |
| **$A_0$** | 0.299 | 0.351 | 0.545 |
| **A** | 0.004 | 0.014 | 0.005 |
| **B** | 0.071 | 0.066 | 0.136 |
| **f** | 0.11 | 0.12 | 0.11 |
| **$R^2$** | 0.913 | 0.847 | 0.935 |
| **adj. $R^2$** | 0.847 | 0.732 | 0.887 |

| **Media richness depletion** | | | | |
|---|---|---|---|---|
| | **0.5× (125 min)** | **0.75× (180 min)** | **0.5× (180 min)** | **0.25× (180 min)** |
| **$A_0$** | 0.368 | 0.291 | 0.494 | 0.694 |
| **A** | 0.006 | 0.031 | 0.008 | 0.020 |
| **B** | 0.065 | 0.012 | 0.454 | 0.182 |
| **f** | 0.11 | 0.13 | 0.10 | 0.10 |
| **$R^2$** | 0.734 | 0.943 | 0.868 | 0.785 |
| **adj. $R^2$** | 0.637 | 0.92 | 0.77 | 0.624 |

| **Rifampicin (2.5 ng/µL)** | |
|---|---|
| **$A_0$** | 0.728 |
| **A** | 0.001 |
| **B** | 0.052 |
| **f** | 0.12 |
| **$R^2$** | 0.700 |
| **adj. $R^2$** | 0.653 |

**Table S4. Parameters of the best fitting sine function**. We used a sine function of the form $g(L_G) = A_0 + A.L_G + B.\sin(2\pi.f.L_G)$. Also shown are the $R^2$ values to evaluate if the sinusoidal curve is well fitted, along with the adjusted (adj.) $R^2$ values, which consider the number of parameters used to fit the curve. Related to Supplementary Figure S8.

| **Novobiocin** | | | |
|---|---|---|---|
| | **50 ng/µL** | **100 ng/µL** | **500 ng/µL** |
| **$A_0$** | 0.331 | 0.447 | 0.645 |
| **A** | 0.001 | 0.003 | -0.002 |
| **B** | 0.048 | 0.081 | 0.136 |
| **f** | 0.10 | 0.10 | 0.11 |
| **$R^2$** | 0.875 | 0.950 | 0.952 |
| **adj. $R^2$** | 0.813 | 0.926 | 0.928 |

| Media richness depletion | | | | |
|---|---|---|---|---|
| | **0.5× (125 min)** | **0.75× (180 min)** | **0.5× (180 min)** | **0.25× (180 min)** |
| **$A_0$** | 0.408 | 0.384 | 0.523 | 0.736 |
| **A** | 0.001 | 0.006 | 0.022 | 0.020 |
| **B** | 0.083 | 0.066 | 0.153 | 0.215 |
| **f** | 0.12 | 0.10 | 0.10 | 0.10 |
| **$R^2$** | 0.912 | 0.618 | 0.966 | 0.941 |
| **adj. $R^2$** | 0.868 | 0.587 | 0.949 | 0.912 |

| **Rifampicin (2.5 ng/µL)** | |
|---|---|
| **$A_0$** | 0.851 |
| **A** | 0.014 |
| **B** | 0.141 |
| **f** | 0.10 |
| **$R^2$** | 0.838 |
| **adj. $R^2$** | 0.757 |

**Table S5. Parameters of the best fitting sine function**. We used a sine function of the form $g(L_G) = A_0 + A.L_G + B.\sin(2\pi.f.L_G)$. Also shown are the $R^2$ values to evaluate if the sinusoidal curve is well fitted, along with the adjusted (adj.) $R^2$ values, which consider the number of parameters used to fit the curve. Related to Supplementary Figure S9.

| **Novobiocin** | | | |
|---|---|---|---|
| | **50 ng/µL** | **100 ng/µL** | **500 ng/µL** |
| **$A_0$** | 0.331 | 0.497 | 0.734 |
| **A** | 0.004 | 0.028 | -0.003 |
| **B** | 0.090 | 0.090 | 0.241 |
| **f** | 0.13 | 0.13 | 0.11 |
| **$R^2$** | 0.893 | 0.903 | 0.956 |
| **adj. $R^2$** | 0.840 | 0.856 | 0.935 |

| **Media richness depletion** | | | | |
|---|---|---|---|---|
| | **0.5× (125 min)** | **0.75× (180 min)** | **0.5× (180 min)** | **0.25× (180 min)** |
| **$A_0$** | 0.401 | 0.472 | 0.65 | 0.839 |
| **A** | -0.020 | -0.021 | -0.035 | 0.005 |
| **B** | 0.115 | 0.126 | 0.246 | 0.289 |
| **f** | 0.12 | 0.12 | 0.12 | 0.12 |
| **$R^2$** | 0.932 | 0.951 | 0.869 | 0.911 |
| **adj. $R^2$** | 0.899 | 0.927 | 0.803 | 0.867 |

| **Rifampicin (2.5 ng/µL)** | |
|---|---|
| **$A_0$** | 0.901 |
| **A** | -0.004 |
| **B** | 0.224 |
| **f** | 0.11 |
| **$R^2$** | 0.878 |
| **adj. $R^2$** | 0.817 |

**Table S6. $\mu_{|\Delta LFC|}$ is higher between genes not in the same operon.** $\mu_{|\Delta LFC|}$ between the most downstream gene of each operon and the gene after the end of the operon. For comparison, the second column shows $\mu_{|\Delta LFC|}$ between pairs of genes in the same operon.

| **Conditions** | **$\mu_{|\Delta LFC|}$ between the most downstream gene of each operon and its nearest neighbor gene, outside of the operon ($L_G$ = 0)** | **$\mu_{|\Delta LFC|}$ between next near neighbor genes in the same operon ($L_G$ = 0)** |
|---|---|---|
| **Novobiocin$_{50\ ng/\mu L}$** | 0.51 | 0.28 |
| **Novobiocin$_{100\ ng/\mu L}$** | 0.75 | 0.34 |
| **Novobiocin$_{500\ ng/\mu L}$** | 1.24 | 0.50 |
| **LB$_{0.5\times(125\ min)}$** | 0.58 | 0.30 |
| **LB$_{0.75\times(180\ min)}$** | 0.63 | 0.29 |
| **LB$_{0.5\times(180\ min)}$** | 0.82 | 0.38 |
| **LB$_{0.25\times(180\ min)}$** | 1.18 | 0.54 |
| **Rifampicin$_{2.5\ ng/\mu L}$** | 1.70 | 0.70 |

**Table S7. Parameters of the best fitting function**. We tested $f(x) = A + B.x$ and $g(L_G) = A_0 + A.L_G + B.\sin(2\pi.f.L_G)$. $R^2$ evaluates the goodness of fit. The adjusted (adj.) $R^2$ further considers the number of parameters used to fit the curve. In all cases, the best model had both the highest $R^2$ value, as well as the lowest AIC and BIC. Related to Figures 2D-2E.

<table>
<tr><th colspan="20">Novobiocin (No internal TSSs)</th></tr>
<tr><td colspan="5"></td><td colspan="5">50 ng/µL</td><td colspan="5">100 ng/µL</td><td colspan="5">500 ng/µL</td></tr>
<tr><td colspan="5">A</td><td colspan="5">0.31</td><td colspan="5">0.35</td><td colspan="5">0.51</td></tr>
<tr><td colspan="5">B</td><td colspan="5">0.04</td><td colspan="5">0.08</td><td colspan="5">0.13</td></tr>
<tr><td colspan="5">AIC</td><td colspan="5">-35.51</td><td colspan="5">-39.87</td><td colspan="5">-29.19</td></tr>
<tr><td colspan="5">BIC</td><td colspan="5">-36.29</td><td colspan="5">-40.65</td><td colspan="5">-29.98</td></tr>
<tr><td colspan="5">adj. $R^2$</td><td colspan="5">0.872</td><td colspan="5">0.986</td><td colspan="5">0.953</td></tr>
<tr><th colspan="20">Media richness depletion (No internal TSS)</th></tr>
<tr><td colspan="4"></td><td colspan="4">0.5× (125 min)</td><td colspan="4">0.75× (180 min)</td><td colspan="4">0.5× (180 min)</td><td colspan="4">0.25× (180 min)</td></tr>
<tr><td colspan="4">A</td><td colspan="4">0.32</td><td colspan="4">0.31</td><td colspan="4">0.39</td><td colspan="4">0.56</td></tr>
<tr><td colspan="4">B</td><td colspan="4">0.07</td><td colspan="4">0.06</td><td colspan="4">0.11</td><td colspan="4">0.20</td></tr>
<tr><td colspan="4">AIC</td><td colspan="4">-36.54</td><td colspan="4">-31.53</td><td colspan="4">-37,67</td><td colspan="4">-29.64</td></tr>
<tr><td colspan="4">BIC</td><td colspan="4">-37.32</td><td colspan="4">-32.31</td><td colspan="4">-38.45</td><td colspan="4">-30.42</td></tr>
<tr><td colspan="4">adj. $R^2$</td><td colspan="4">0.96</td><td colspan="4">0.87</td><td colspan="4">0.98</td><td colspan="4">0.98</td></tr>
<tr><th colspan="20">Rifampicin 2.5 ng/µL (No internal TSS)</th></tr>
<tr><td colspan="10">A</td><td colspan="10">0.68</td></tr>
<tr><td colspan="10">B</td><td colspan="10">0.15</td></tr>
<tr><td colspan="10">AIC</td><td colspan="10">-33.92</td></tr>
<tr><td colspan="10">BIC</td><td colspan="10">-34.70</td></tr>
<tr><td colspan="10">adj. $R^2$</td><td colspan="10">0.98</td></tr>
<tr><th colspan="20">Novobiocin ( ≥ 1 internal TSS)</th></tr>
<tr><td colspan="5"></td><td colspan="5">50 ng</td><td colspan="5">100 ng</td><td colspan="5">500 ng</td></tr>
<tr><td colspan="5">$A_0$</td><td colspan="5">0.39</td><td colspan="5">0.38</td><td colspan="5">0.53</td></tr>
<tr><td colspan="5">A</td><td colspan="5">-0.07</td><td colspan="5">0.01</td><td colspan="5">0.03</td></tr>
<tr><td colspan="5">B</td><td colspan="5">0.31</td><td colspan="5">0.06</td><td colspan="5">0.05</td></tr>
<tr><td colspan="5">f</td><td colspan="5">0.05</td><td colspan="5">0.06</td><td colspan="5">0.07</td></tr>
<tr><td colspan="5">AIC</td><td colspan="5">-40.93</td><td colspan="5">-38.10</td><td colspan="5">-36.99</td></tr>
<tr><td colspan="5">BIC</td><td colspan="5">-42.50</td><td colspan="5">-39.66</td><td colspan="5">-38.55</td></tr>
<tr><td colspan="5">adj. $R^2$</td><td colspan="5">0.88</td><td colspan="5">0.94</td><td colspan="5">0.90</td></tr>
<tr><th colspan="20">Media richness depletion ( ≥ 1 internal TSS)</th></tr>
<tr><td colspan="4"></td><td colspan="4">0.5× (125min)</td><td colspan="4">0.75× (180min)</td><td colspan="4">0.5× (180min)</td><td colspan="4">0.25× (180min)</td></tr>
<tr><td colspan="4">$A_0$</td><td colspan="4">0.34</td><td colspan="4">0.36</td><td colspan="4">0.42</td><td colspan="4">0.63</td></tr>
<tr><td colspan="4">A</td><td colspan="4">0.01</td><td colspan="4">0.01</td><td colspan="4">0.02</td><td colspan="4">0.04</td></tr>
<tr><td colspan="4">B</td><td colspan="4">0.05</td><td colspan="4">0.06</td><td colspan="4">0.09</td><td colspan="4">0.12</td></tr>
<tr><td colspan="4">f</td><td colspan="4">0.08</td><td colspan="4">0.08</td><td colspan="4">0.07</td><td colspan="4">0.09</td></tr>
<tr><td colspan="4">AIC</td><td colspan="4">-42.98</td><td colspan="4">-36.01</td><td colspan="4">-31.31</td><td colspan="4">-27.29</td></tr>
<tr><td colspan="4">BIC</td><td colspan="4">-44.55</td><td colspan="4">-37.58</td><td colspan="4">-32.87</td><td colspan="4">-28.86</td></tr>
<tr><td colspan="4">adj. $R^2$</td><td colspan="4">0.94</td><td colspan="4">0.89</td><td colspan="4">0.86</td><td colspan="4">0.79</td></tr>
<tr><th colspan="20">Rifampicin 2.5 ng/µL ( ≥ 1 internal TSS)</th></tr>
<tr><td colspan="10">$A_0$</td><td colspan="10">1.04</td></tr>
<tr><td colspan="10">A</td><td colspan="10">-0.24</td></tr>
<tr><td colspan="10">B</td><td colspan="10">0.91</td></tr>
</table>

| | |
|---|---|
| **f** | 0.05 |
| **AIC** | -26.58 |
| **BIC** | -28.14 |
| **adj. $R^2$** | 0.69 |

**Table S8. Slopes of the best fitting lines (m) of $\mu_{|\Delta LFC|}(L_G)$.** Compared are pairs of genes in the same operon without TSSs in between them, and pairs of genes in the same operon without TSSs in between them and that are supercoiling sensitive.

| **Conditions** | ***Slope m (no TSS but SS)*** | ***Slope m (no TSS and no SS)*** |
|---|---|---|
| **Novobiocin$_{50\ ng/\mu L}$** | 0.04 | 0.04 |
| **Novobiocin$_{100\ ng/\mu L}$** | 0.08 | 0.05 |
| **Novobiocin$_{500\ ng/\mu L}$** | 0.13 | 0.05 |
| **LB$_{0.5\times\ 125\ min}$** | 0.07 | 0.03 |
| **LB$_{0.75\times\ 180\ min}$** | 0.06 | 0.02 |
| **LB$_{0.5\times\ 180\ min}$** | 0.11 | 0.04 |
| **LB$_{0.25\times\ 180\ min}$** | 0.20 | 0.14 |
| **Rifampicin$_{2.5\ ng/\mu L}$** | 0.15 | 0.08 |

**Table S9. Expression levels of synthetic genetic constructs.** Mean and standard error of the mean (SEM) expression levels of synthetic gene constructs under different conditions.

| Conditions | Constructs | Mean (μ) | Standard error of the mean (SEM) |
|---|---|---|---|
| **Control** | Lac | 456.9 | 2.0 |
| **Control** | Tet | 4850.6 | 16.6 |
| **Control** | Tet-Lac | 439.6 | 1.4 |
| **Novobiocin$_{50\ ng/\mu L}$** | Lac | 358.1 | 2.5 |
| **Novobiocin$_{50\ ng/\mu L}$** | Tet | 4363.2 | 27.0 |
| **Novobiocin$_{50\ ng/\mu L}$** | Tet-Lac | 361.7 | 1.7 |
| **Novobiocin$_{100\ ng/\mu L}$** | Lac | 328.9 | 2.0 |
| **Novobiocin$_{100\ ng/\mu L}$** | Tet | 4021.1 | 27.4 |
| **Novobiocin$_{100\ ng/\mu L}$** | Tet-Lac | 310.2 | 2.1 |
| **Novobiocin$_{500\ ng/\mu L}$** | Lac | 291.2 | 2.7 |
| **Novobiocin$_{500\ ng/\mu L}$** | Tet | 3845.1 | 27.3 |
| **Novobiocin$_{500\ ng/\mu L}$** | Tet-Lac | 237.8 | 2.0 |
| **LB$_{0.75\times}$** | Lac | 357.3 | 3.1 |
| **LB$_{0.75\times}$** | Tet | 4458.2 | 48.6 |
| **LB$_{0.75\times}$** | Tet-Lac | 343.8 | 2.3 |
| **LB$_{0.5\times}$** | Lac | 304.4 | 3.2 |
| **LB$_{0.5\times}$** | Tet | 4175.9 | 42.5 |
| **LB$_{0.5\times}$** | Tet-Lac | 273.1 | 1.90 |
| **LB$_{0.25\times}$** | Lac | 252.8 | 2.53 |
| **LB$_{0.25\times}$** | Tet | 4102.7 | 26.3 |
| **LB$_{0.25\times}$** | Tet-Lac | 244.0 | 6.8 |
| **Rifampicin$_{2.5\ ng/\mu L}$** | Lac | 128.4 | 0.8 |
| **Rifampicin$_{2.5\ ng/\mu L}$** | Tet | 4188.0 | 14.4 |
| **Rifampicin$_{2.5\ ng/\mu L}$** | Tet-Lac | 193.2 | 0.7 |

**Table S10. DNA sequences that are known to enhance premature terminations**. The table shows the frequency at which specific sequences are found along the genes in operons for 0 and 1 degrees of freedom (Methods section ‘Identifying premature terminations and pause sequences’).

| **Sequence (Reference)** | ***μ* Frequency (0 degrees of freedom)** | ***μ* Frequency (1 degree of freedom)** |
|---|---|---|
| **CGGGTAGATCCG** | 0 | 0.0012 |
| **GGTGAAACCGCA** | 0.0048 | 0.0156 |
| **CGGTAAAGTGTA** | 0.0012 | 0.0024 |
| **CGTATCACTGCG** | 0.0024 | 0.0048 |
| **CGATGTGTGCTG** | 0.0012 | 0.0072 |
| **TCCGCCCGCATA** | 0 | 0.0024 |
| **CATCTTTTGACA** | 0.0012 | 0.0024 |
| **AGGCCGCCGTAT** | 0.0012 | 0.0096 |
| **ACCACCATCATC** | 0.0012 | 0.0108 |
| **AAGACATTCAGA** | 0.0012 | 0.0048 |
| **AGATCGACCTGT** | 0 | 0.0024 |
| **TTGAAAAAGTTA** | 0.0012 | 0.0048 |
| **AATCCGTGATAA** | 0.0012 | 0.0036 |
| **TAACAAGCTGCA** | 0.0012 | 0.0048 |
| **TOTAL** | 0.01800 | 0.0768 |
| **Homopolymetric Sequences** | **Frequency (0 degrees of freedom)** | **Frequency (1 degree of freedom)** |
| **AAAAAAAAA** | 0.0048 | 0.2209 |
| **TTTTTTTTT** | 0.0048 | 0.2761 |
| **TOTAL** | 0.0096 | 0.4969 |

**Table S11. Parameters of the best fitting sine function**. Supplementary Figure S29. We used a sine function of the form $g(L_G) = A_0 + A.L_G + B.\sin(2\pi.f.L_G)$. Also shown are the $R^2$ values to evaluate if the sinusoidal curve is well fitted, along with the adjusted (adj.) $R^2$ values, which consider the number of parameters used to fit the curve.

| **Antibiotics** | | |
|---|---|---|
| | **Chloramphenicol** | **Erythromycin** |
| **$A_0$** | 0.72 | 0.78 |
| **A** | 0.02 | 0.01 |
| **B** | 0.06 | 0.09 |
| **f** | 0.13 | 0.11 |
| **$R^2$** | 0.90 | 0.90 |
| **adj. $R^2$** | 0.85 | 0.85 |

| **Environment** | | | |
|---|---|---|---|
| | **Micro-Aerobic** | **Heat-shock** | **High-Pressure** |
| **$A_0$** | 0.41 | 0.31 | 0.37 |
| **A** | -0.010 | -0.003 | 0.008 |
| **B** | 0.12 | 0.04 | 0.07 |
| **f** | 0.08 | 0.11 | 0.10 |
| **$R^2$** | 0.96 | 0.83 | 0.95 |
| **adj. $R^2$** | 0.94 | 0.75 | 0.93 |

| **Anaerobic** | | | | | |
|---|---|---|---|---|---|
| | **0.5 min** | **1 min** | **2 min** | **5 min** | **10 min** |
| **$A_0$** | 0.26 | 0.33 | 0.42 | 0.41 | 0.39 |
| **A** | -0.005 | 0.004 | 0.001 | 0.010 | 0.010 |
| **B** | 0.02 | 0.04 | 0.11 | 0.10 | 0.08 |
| **f** | 0.10 | 0.13 | 0.10 | 0.10 | 0.12 |
| **$R^2$** | 0.46 | 0.89 | 0.95 | 0.95 | 0.96 |
| **adj. $R^2$** | 0.42 | 0.83 | 0.93 | 0.93 | 0.95 |

| **Mercury** | | | |
|---|---|---|---|
| | **10 min** | **30 min** | **60 min** |
| **$A_0$** | 0.28 | 0.38 | 0.46 |
| **A** | -0.002 | 0.020 | -0.001 |
| **B** | 0.04 | 0.06 | 0.12 |
| **f** | 0.09 | 0.10 | 0.10 |
| **$R^2$** | 0.78 | 0.90 | 0.95 |
| **adj. $R^2$** | 0.68 | 0.85 | 0.92 |

**Table S12. 2-sample K-S tests comparing the distribution of the number of operons of a given number of genes of *E. coli* with the ones of other species**. For *p*-values < 0.05, we reject the null hypothesis that the two distributions are the same.

| **Species** | **Mean *p*-values** |
|---|---|
| *Bacillus subtillis* | 0.87 |
| *Corynebacterium glutamicum* | 0.72 |
| *Helicobacter pylori* | 0.06 |
| *Legionella pneumophila* | < 0.05 |
| *Listeria monocytogenes* | 0.62 |
| *Mycoplasma pneumoniae* | 0.52 |
| *Photobacterium profundum* | < 0.05 |

**Table S13. Parameters of the best fitting sine function**. Related to Figure 5A-D. We used a sine function of the form $g(L_G) = A_0 + A.L_G + B.\sin(2\pi.f.L_G)$. Also shown are the $R^2$ values to evaluate if the sinusoidal curve is well fitted, along with the adjusted (adj.) $R^2$ values, which consider the number of parameters used to fit the curve.

| | **Antimicrobe** | **Biofilm** |
|---|---|---|
| **$A_0$** | 0.20 | 1.3 |
| **A** | 0.007 | -0.020 |
| **B** | 0.05 | 0.20 |
| **f** | 0.08 | 0.10 |
| **$R^2$** | 0.91 | 0.86 |
| **adj. $R^2$** | 0.86 | 0.79 |
| | **Amphotericin** | **Berberine** |
| **$A_0$** | 0.21 | 0.77 |
| **A** | 0.002 | -0.020 |
| **B** | 0.03 | 0.25 |
| **f** | 0.11 | 0.10 |
| **$R^2$** | 0.75 | 0.93 |
| **adj. $R^2$** | 0.63 | 0.90 |
| **Salinity** | | |
| | **30 min** | **60 min** |
| **$A_0$** | 1.48 | 1.22 |
| **A** | -0.060 | -0.010 |
| **B** | 0.14 | 0.11 |
| **f** | 0.13 | 0.13 |
| **$R^2$** | 0.92 | 0.86 |
| **adj. $R^2$** | 0.89 | 0.79 |
| | **xyn-A mutant** | **non-motile** |
| **$A_0$** | 0.28 | 0.30 |
| **A** | -0.070 | -0.002 |
| **B** | 0.67 | 0.05 |
| **f** | 0.10 | 0.09 |
| **$R^2$** | 0.84 | 0.88 |
| **adj. $R^2$** | 0.76 | 0.82 |

**Table S14. Parameters of the best fitting sine function**. Related to Figure 5E-F. We used a sine function of the form $g(L_G) = A_0 + A.L_G + B.\sin(2\pi.f.L_G)$. Also shown are the $R^2$ values to evaluate if the sinusoidal curve is well fitted, along with the adjusted (adj.) $R^2$ values, which consider the number of parameters used to fit the curve.

| | **Glucose** | **Indole** |
|---|---|---|
| **$A_0$** | 0.27 | 0.36 |
| **A** | 0.007 | 0.008 |
| **B** | 0.15 | 0.08 |
| **f** | 0.12 | 0.09 |
| **$R^2$** | 0.91 | 0.88 |
| **adj. $R^2$** | 0.87 | 0.82 |
| | **Pipecolic** | **Proline** |
| **$A_0$** | 2.85 | 0.26 |
| **A** | -2.600 | -0.003 |
| **B** | 0.09 | 0.07 |
| **f** | 0.12 | 0.12 |
| **$R^2$** | 0.77 | 0.88 |
| **adj. $R^2$** | 0.65 | 0.82 |

**Table S15 Parameters of the best fitting sine function**. Related to Figure 5G-H. We used a sine function of the form $g(L_G) = A_0 + A.L_G + B.\sin(2\pi.f.L_G)$. Also shown are the $R^2$ values to evaluate if the sinusoidal curve is well fitted, along with the adjusted (adj.) $R^2$ values, which consider the number of parameters used to fit the curve.

| pH 5.3 vs pH 7 (1 hour) | | | |
|---|---|---|---|
| | **Wild Type** | **rdxA mutant** | **ArsS mutant** |
| **$A_0$** | 0.49 | 0.43 | 0.30 |
| **A** | 0.02 | 0.02 | 0.001 |
| **B** | 0.07 | 0.03 | 0.07 |
| **f** | 0.07 | 0.10 | 0.07 |
| **$R^2$** | 0.91 | 0.90 | 0.95 |
| **adj. $R^2$** | 0.86 | 0.85 | 0.93 |
| **pH 5.3 vs pH 7 (7 hours) WT** | | | |
| **$A_0$** | 0.57 | | |
| **A** | 0.02 | | |
| **B** | 0.10 | | |
| **f** | 0.10 | | |
| **$R^2$** | 0.81 | | |
| **adj. $R^2$** | 0.72 | | |

**Table S16. Parameter values for simulations.**

| **Parameter** | **Value ($s^{-1}$)** | **Reference** |
|---|---|---|
| $\mathbf{k_{cc}}$ | 0.0013 | (*65*) |
| $\mathbf{k_{oc}}$ | 0.0052 | (*65*) |
| $\mathbf{k_{esc}}$ | 1 | (*96*) |
| $\mathbf{k_{el}}$ | nucleotide length of the gene × $42^{-1}$ | (*67*, *102*) |
| $\mathbf{k^{CC}_{col}}$, $\mathbf{k^{OC}_{col}}$ | 20 | Rates set to allow equal chances of fall-off of elongating and promoter-bound RNAPs, when colliding, regarding if the RNAP is in CC or OC. |
| $\mathbf{k_{fall}}$ | For each gene, we produce a random number from Absolute (Normal(0.0015, 0.001)) in the control, and Absolute(Normal(0.0003, 0.001)) in the stress condition. | (*97*, *103*) for the rate in the control condition. Other rates are estimated based on the empirical data here reported. |

**Table S17. Spatial frequency (*f*) of the best fitting sinusoidal curve of the form $g(L_G) = A_0 + A.L_G + B.\sin(2\pi.f.L_G)$ of $\mu_{|\Delta LFC|}(L_G)$.** Considered are operons with three or more genes with a given number of TSS upstream from the primary TSS.

| **Conditions** | ***f*** **( ≥ 3 TSS)** | ***f*** **( < 3 TSS)** |
|---|---|---|
| **Novobiocin$_{50\ ng/\mu L}$** | 0.09 | 0.08 |
| **Novobiocin$_{100\ ng/\mu L}$** | 0.09 | 0.07 |
| **Novobiocin$_{500\ ng/\mu L}$** | 0.09 | 0.08 |
| **LB$_{0.5\times(125\ min)}$** | 0.09 | 0.09 |
| **LB$_{0.75\times(125\ min)}$** | 0.10 | 0.12 |
| **LB$_{0.5\times(125\ min)}$** | 0.11 | 0.09 |
| **LB$_{0.25\times(125\ min)}$** | 0.10 | 0.07 |
| **Rifampicin$_{2.5\ ng/\mu L}$** | 0.09 | 0.07 |

**Table S18. List of YFP-fusion strains (*32*) used in this study.**

| No. | Strain name | Genotype | Source |
|---|---|---|---|
| 1 | MG1655 | λ-, rph-1 | Yale CGSC (CGSC # 6300) |
| 2 | SX1031 | F-, Δ(argF-lac)169, gal-490, Δ(modF-ybhJ)803, λ[cI857 Δ(cro-bioA)],def-791-YFP(::cat), IN(rrnD-rrnE)1, rph-1 | Yale CGSC (CGSC #12586) |
| 3 | SX1834 | F-, Δ(argF-lac)169, gal-490, Δ(modF-ybhJ)803, λ[cI857 Δ(cro-bioA)],fmt-791-YFP(::cat), IN(rrnD-rrnE)1, rph-1 | Yale CGSC (CGSC #13389) |
| 4 | SX1681 | F-, Δ(argF-lac)169, gal-490, Δ(modF-ybhJ)803, λ[cI857 Δ(cro-bioA)],hemE-791-YFP(::cat), IN(rrnD-rrnE)1, rph-1 | Yale CGSC (CGSC #13236) |
| 5 | SX1295 | F-, Δ(argF-lac)169, gal-490, Δ(modF-ybhJ)803, λ[cI857 Δ(cro-bioA)],nfi-791-YFP(::cat), IN(rrnD-rrnE)1, rph-1 | Yale CGSC (CGSC #12850) |
| 6 | SX1463 | F-, Δ(argF-lac)169, gal-490, Δ(modF-ybhJ)803, λ[cI857 Δ(cro-bioA)],rpoZ-793-YFP(::cat), IN(rrnD-rrnE)1, rph-1 | Yale CGSC (CGSC #13018) |
| 7 | SX1068 | F-, Δ(argF-lac)169, gal-490, Δ(modF-ybhJ)803, λ[cI857 Δ(cro-bioA)],spoT-791-YFP(::cat), IN(rrnD-rrnE)1, rph-1 | Yale CGSC (CGSC #12623) |
| 8 | SX1139 | F-, Δ(argF-lac)169, gal-490, Δ(modF-ybhJ)803, λ[cI857 Δ(cro-bioA)],sspA-791-YFP(::cat), IN(rrnD-rrnE)1, rph-1 | Yale CGSC (CGSC #12694) |
| 9 | SX1075 | F-, Δ(argF-lac)169, gal-490, Δ(modF-ybhJ)803, λ[cI857 Δ(cro-bioA)],sspB-792-YFP(::cat), IN(rrnD-rrnE)1, rph-1 | Yale CGSC (CGSC #12630) |
| 10 | SX1718 | F-, Δ(argF-lac)169, gal-490, Δ(modF-ybhJ)803, λ[cI857 Δ(cro-bioA)],wrbA-791-YFP(::cat), IN(rrnD-rrnE)1, rph-1 | Yale CGSC (CGSC #13273) |
| 11 | SX1975 | F-, Δ(argF-lac)169, gal-490, Δ(modF-ybhJ)803, λ[cI857 Δ(cro-bioA)],yccJ-791-YFP(::cat), IN(rrnD-rrnE)1, rph-1 | Yale CGSC (CGSC #13530) |
| 12 | SX1158 | F-, Δ(argF-lac)169, gal-490, Δ(modF-ybhJ)803, λ[cI857 Δ(cro-bioA)],ybeZ-797-YFP(::cat), IN(rrnD-rrnE)1, rph-1 | Yale CGSC (CGSC #12713) |
| 13 | SX1146 | F-, Δ(argF-lac)169, gal-490, Δ(modF-ybhJ)803, λ[cI857 Δ(cro-bioA)],ybeY-796-YFP(::cat), IN(rrnD-rrnE)1, rph-1 | Yale CGSC (CGSC #12701) |
| 14 | SX1885 | F-, Δ(argF-lac)169, gal-490, Δ(modF-ybhJ)803, λ[cI857 Δ(cro-bioA)],yqgE-791-YFP(::cat), IN(rrnD-rrnE)1, rph-1 | Yale CGSC (CGSC #13440) |
| 15 | SX2009 | F-, Δ(argF-lac)169, gal-490, Δ(modF-ybhJ)803, λ[cI857 Δ(cro-bioA)],yqgF-792-YFP(::cat), IN(rrnD-rrnE)1, rph-1 | Yale CGSC (CGSC #13564) |

**Table S19**. **List of the variables, and their definitions, used in the study.**

| **Variable** | **Definition** |
|---|---|
| **LFC** | $\log_2$ fold change |
| $\boldsymbol{P}$ | Position of each gene in an operon. The first gene downstream from the primary promoter is assigned to be in position 1 (illustrated in Figure 1C). |
| $\boldsymbol{L_G}$ | Distance between genes in the same operon (in ‘gene’ units) |
| $\boldsymbol{L_P}$ | Distances between consecutive promoters in the same operon (measured in ‘gene’ units as illustrated in Figure 1C) |
| $\mathbf{\lvert\Delta LFC\rvert}$ | Absolute difference in the $\log_2$ fold change between gene pairs of the same operon |
| $\boldsymbol{\mu_{\lvert\Delta LFC\rvert}}$ | Mean absolute differences in the $\log_2$ fold change between gene pairs of the same operon |
| $\mathbf{\mu\lvert\Delta pn\rvert}$ | Mean absolute differences in the protein numbers between gene pairs of the same operon |
| $\boldsymbol{r_f}$ | Rate of premature termination for each gene in the operon |
| $\boldsymbol{R_f}$ | Operon-wide average rates of premature termination |
| $\boldsymbol{f_H}$ | frequency of Homopolymeric AT tracts in the sequence of the gene |
| $\boldsymbol{f_{pause}}$ | frequency of pause sequences in the gene |
| **PSB** | Positive supercoiling buildup |
| **SS** | Positive supercoiling sensitive genes |
| **TF** | Transcription factor |
| **TSS** | Transcription start site |
| **TTS** | Transcription termination site |
| ***pTTS*** | Transcription termination sites reported in (*36*) |
| **Terms used in the model** | |
| $\boldsymbol{k_{cc}}$ | Rate of closed complex formation. |
| $\boldsymbol{k_{oc}}$ | Rate of Open complex formation. |
| $\boldsymbol{k_{esc}}$ | Rate of promoter escape. |
| $\boldsymbol{k_{el}}$ | Rate of RNAP elongation. |
| $\boldsymbol{k^{CC}_{col}}$ | Rate of RNAP fall-off due to collision between downstream RNAP sitting on the closed complex and upstream elongating RNAP. |
| $\boldsymbol{k^{OC}_{col}}$ | Rate of RNAP fall-off due to collision between downstream RNAP sitting on the open complex and upstream elongating RNAP. |
| $\boldsymbol{k_{fall}}$ | Rate of RNAP fall-off. |
| $\boldsymbol{\mu_{\lvert\Delta RNA\rvert}}$ | Mean absolute differences in the RNA numbers between gene pairs. |
| $\boldsymbol{A_0}$ | y axis intercept of the mathematical model |
| $\boldsymbol{A}$ | Coefficient of linear term of the mathematical model |
| $\boldsymbol{B}$ | Coefficient of linear term of the mathematical model |
| $\boldsymbol{f}$ | Spatial frequency of the mathematical model |
